# Extent, heritability, and functional relevance of single cell expression variability in highly homogeneous populations of human cells

**DOI:** 10.1101/574426

**Authors:** Daniel Osorio, Xue Yu, Yan Zhong, Guanxun Li, Peng Yu, Erchin Serpedin, Jianhua Huang, James J. Cai

## Abstract

Because of recent technological developments, single-cell assays such as single-cell RNA sequencing (scRNA-seq) have become much more widely available and have achieved unprecedented resolution in revealing cell heterogeneity. The extent of intrinsic cell-to-cell variability in gene expression, or *single cell expression variability* (scEV), has thus been increasingly appreciated. However, it remains unclear whether this variability is functionally important and, if so, what its implications are for multi-cellular organisms. We therefore analyzed multiple scRNA-seq data sets from lymphoblastoid cell lines (LCLs), lung airway epithelial cells (LAECs), and dermal fibroblasts (DFs). For each of the three cell types, we estimated scEV in homogeneous populations of cells; we identified 465, 466, and 291 highly variable genes (HVGs), respectively. These HVGs were enriched with specific functions precisely relevant to the cell types, from which the scRNA-seq data used to identify HVGs were generated—e.g., HVGs identified in lymphoblastoid cells were enriched in cytokine signaling pathways, LAECs collagen formation, and DFs keratinization. HVGs were deeply embedded in gene regulatory networks specific to corresponding cell types. We also found that scEV is a heritable trait, partially determined by cell donors’ genetic makeups. Furthermore, across genes, especially immune genes, levels of scEV and between-individual variability in gene expression were positively correlated, suggesting a potential link between the two variabilities measured at different organizational levels. Taken together, our results support the “variation is function” hypothesis, which postulates that scEV is required for higher-level system function. Thus, we argue that quantifying and characterizing scEV in relevant cell types may deepen our understating of normal as well as pathological cellular processes.

## Introduction

Cells are fundamental units of cellular function. Cells in multi-cellular organisms can be organized into groups, or cell types, based on shared features that are quantifiable. A multicellular organism is usually composed of cells of many different types—each is a distinct functional entity differing from the other. Within the same cell type, cells are nearly identical and are considered to carry the same function.

The recent development of single cell RNA sequencing (scRNA-seq) technologies has brought the increasingly high-resolution measurements of gene expression in single cells (Zhang et al. 2019). This power has been widely adopted to refine the categories of known cell types and analyze complex tissues systematically and reproducibly (Buettner et al. 2015). The power of scRNA-seq has also been harnessed to identify novel cellular states among the same type of cells (Trapnell 2015).

Cells of the same type and the same state may still show marked intrinsic cell-to-cell variability in gene expression or *single cell expression variability* (scEV), even under the same environmental conditions (Ko 1992; Fiering et al. 2000; Raj and van Oudenaarden 2008). The importance of this intrinsic variability is increasingly appreciated (Eldar and Elowitz 2010; Pelkmans 2012). Changes in the magnitude of scEV have been associated with development (Wernet et al. 2006; Chang et al. 2008; Faure et al. 2017; Kumar et al. 2017), aging (Martinez-Jimenez et al. 2017; Wiley et al. 2017), and pathological processes (Segerstolpe et al. 2016; Azizi et al. 2018).

Dueck and colleagues (Dueck et al. 2016) put forward the so-called “variation is function” hypothesis, saying that scEV *per se* might be crucial for population-level function. They used the term “single cell variation or variability” to refer to diversity within an ensemble that has been previously defined as being generally homogeneous, rather than diversity of cell types that are clearly distinct and already recognized. The main focus of their question is to ask how the individual cells with different gene expression levels may interact to causally generate higher-level function. If the hypothesis turns out to be true, it means that the intrinsic cell-to-cell variability is an indicator of a diversity of hidden functional capacities, which facilitate the collective behavior of cells. This collective behavior is essential for the function and normal development of cells and tissues (Raj et al. 2010; Tay et al. 2010). The loss of this collective cellular behavior may result in disease. Thus, investigation of the intrinsic cell-to-cell variability may contribute to the understanding of pathological processes associated with disease development.

It is worth noting that the level of intrinsic cell-to-cell variability needs to be measured within a highly homogenous population of cells. This is because many micro-environmental perturbations and stochastic factors at the cellular level are known to change the scEV. These factors may include local cell density, cell size, shape and rate of proliferation, cell cycle, and so on (Snijder et al. 2009; McDavid et al. 2014; Kernfeld et al. 2018; Miragaia et al. 2018; Mitchell et al. 2018). To work on the cell-to-cell variability, these confounding factors have to be controlled.

Exponential scaling of scRNA-seq made it feasible to study scEV across thousands of cells (Svensson et al. 2018), and quantify scEV based on measures of statistical dispersion such as the coefficient of variation (CV)(Geiler-Samerotte et al. 2013; Mar 2019). The sheer number of cells sequenced in a “typical” droplet-based scRNA-seq experiment allows us to filter out for a sizable number of highly homogeneous cells, based on the similarity between their global transcriptional profiles. With these selected cells, we are able to test the “variation is function” hypothesis systematically. Furthermore, using established statistical methods, we are able to control for many sources of technical variation that may confound the measurement of scEV to obtain an unbiased estimate. For instance, single-molecule capture efficiency, 3’ end bias due to single-cell RNA library preparation protocol, and low expression of genes are examples of known sources of technical variation (Marinov et al. 2014), which should be controlled for using statistical means.

The characterization of the impact of scEV on cell function requires the understanding of which genes show greater or less cell-to-cell variability in their expression. These feature genes may carry valuable information that can facilitate the elucidation of underlying regulatory networks (Li and You 2013). Once these genes are identified, a follow-up question is whether they are tissue- or cell type-specific—i.e., whether the same genes will be identified for different tissues or cell types. Our working hypothesis is in line with the “variation is function” hypothesis, that is, different tissues or cell types have different sets of highly variable genes (HVGs), and these HVGs should be enriched with functions that reflect the biological functions of respective tissues or the cell types. To test this, we analyzed three scRNA-seq data sets generated for three different cell types. Each data set contains thousands of cells. For each cell type, we selected a highly homogenous population of cells, with the help of a newly developed dimensionality reduction method, called potential of heat-diffusion for affinity-based trajectory embedding (PHATE)(Moon et al. 2018). We estimated scEV among selected cells for each of these cell types and further systematically characterized functions of identified HVGs. We show that HVGs are highly specific to cell types, i.e., different cell types have different sets of HVGs; functions of HVGs precisely mirror the functions of the corresponding cell types. We explored both the level of scEV and potential mechanism behind the variability across cells, allowing us to understand a previously unexplored aspect of gene regulation in humans.

## Results

### Single-cell RNA sequencing and selection of highly homogenous cells

In this study, we experimented with three different human cell types, namely, lymphoblastoid cell line (LCL), lung airway epithelial cell (LAEC), and dermal fibroblast (DF). We estimated single cell expression variability (scEV) for each of these cell types, individually.

To obtain the scRNA-seq data for LCL, we cultured GM12878, an LCL strain widely used in genomic research, prepared cells using a 10X Genomics Chromium Controller, and sequenced a total of 7,045 cells (Osorio et al. 2019). This data has been deposited in the NCBI GEO database (Accession number GSE126321). For the other two cell types, LAEC and DF, we obtained the scRNA-seq data for 3,863 and 2,553 cells from the studies of (Habiel et al. 2018) and (Hagai et al. 2018), respectively (see **Data Availability**). All scRNA-seq data sets of the three cell types were produced using 10X Genomics droplet-based solution and made use of unique molecular identifiers (UMIs)(Kivioja et al. 2011).

For each cell type, we employed a data analysis procedure, a filter pipeline on scRNA-seq data, to select highly homogenous cells (**Materials and Methods**). These selected cells are a representative population of each the cell type. The main steps of the filter pipeline are depicted in **Supplementary Fig. S1**. Briefly, we first excluded mitochondrial DNA-encoded genes from the analysis. We then excluded cells in the S- or G2/M phases and only retained G1-phase cells. We also excluded cells with library size smaller than 55 percentile or greater than 99 percentile. Finally, we used PHATE to produce the embedding plot of remaining cells to inspect between-cell structure driven by heterogeneity in gene expression. PHATE is a visualization method that captures both local and global nonlinear structure in data by an information-geometry distance between data points (Moon et al. 2018). As seen from the PHATE projection (**Fig. 1A**), several “arms” of cells show the structure of the cell-to-cell relationship. Based on the observation, we manually picked one “core” cell, at the root of arms of cells, in the middle of cell cloud where the majority of cells are clustered. The core cell and 999 nearest cells around the core cell are then selected to form the final cell population, which is used for subsequent data analyses.

**Fig. 1.**
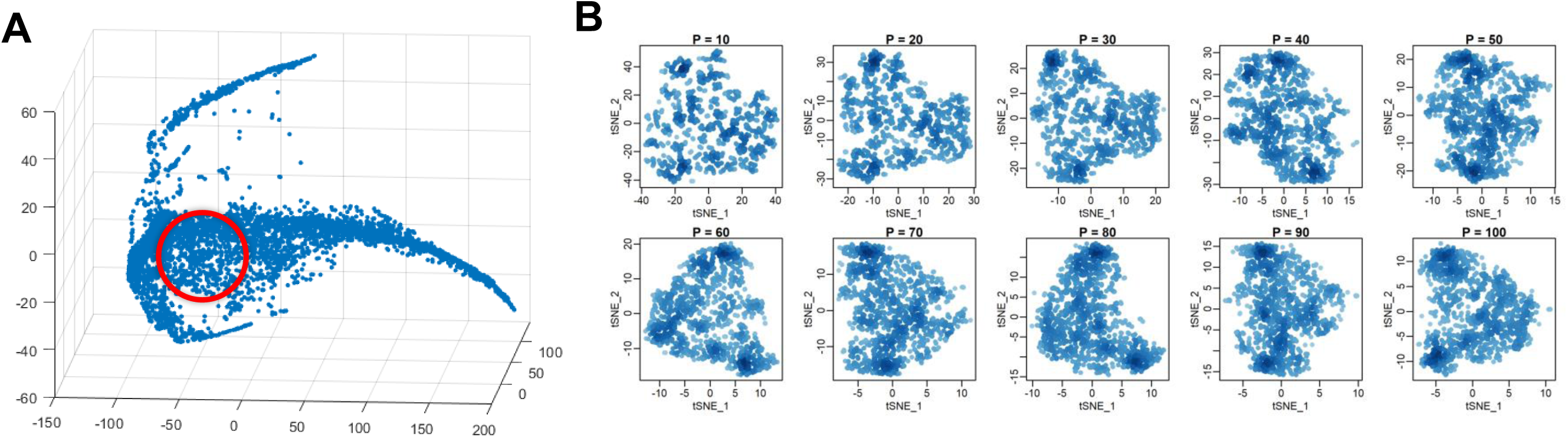
Selection of a highly homogenous cell population for variability analysis. (**A**) Three-dimensional PHATE embedding plot for G1-phase cells of GM12878. Each point represents a single cell in the three-dimensional space. The red circle indicates the approximate positions of 1,000 selected cells. (**B**) Embedding plots generated for the 1,000 selected cells with t-SNE algorithm with a series of perplexity values.

To examine the homogeneity of selected cells, we used t-distributed stochastic neighbor embedding (t-SNE)(van der Maaten and Hinton 2008) to position all 1,000 selected cells in the two-dimensional t-SNE space. Compared to PHATE, t-SNE is a more commonly used nonlinear visualization algorithm for revealing structures in high-dimensional data, emphasizing local neighborhood structure within the data. When running t-SNE, we experimented with a series of perplexity values to produce multiple plots for the same population of selected cells. t-SNE is known to be sensitive to hyperparameters (Becht et al. 2018). In general, when different parameter values are given, t-SNE tends to produce different cell clustering plots. But, for our selected cells, no structure is observed in any of these t-SNE embedding plots (**Fig. 1B**). The same results were obtained for the other two cell types. Thus, we confirm that cells selected with our filter pipeline are highly homogenous populations of representative cells for each cell type.

### Identification of highly variable genes

Highly variable genes (HVGs) are expressed variably across homogeneous cells of the same type. For each cell type, we used the method of (Brennecke et al. 2013) to identify HVGs from scRNA-seq data of the homogeneous population of selected cells. In this method, the relationship between the squared coefficient of variation (CV^2^) of genes and their average expression (μ) is considered. The relationship between log-transformed CV^2^ and log-transformed μ is fitted with a Generalized Linear Model (GLM), and the expected CV^2^ for a given μ is calculated with the fitted curve. The log-transformed ratio between observed CV^2^ and expected CV^2^ [=log(observed CV^2^)- log(expected CV^2^], called “residual variability”, is used as the measurement of scEV. Since the expected CV^2^ captures the variability originated from technical noise, the residual variability is considered to be an unbiased measure of biological variability. In total, we identified 465, 466, and 291 HVGs for LCL, LAEC, and DF, respectively (**Supplementary Tables S1-3**), after controlling for false discovery rate (FDR) at 0.01 (**Materials and Methods**). To visualize expression variability of genes, we plot CV^2^ against μ, both on the logarithmic scale, for LCL (**Fig. 2A**). Each dot represents a gene; all genes together give a characteristic cloud showing the μ and CV^2^ of gene expression. Genes above the GLM fitting curve, e.g., *IGKC, CCL3, LTB*, and *FTL*, are more variable than expectation, whereas genes below the curve, e.g., *TMEM9B* and *RPL17*, are less variable (**Fig. 2B**).

**Fig. 2.**
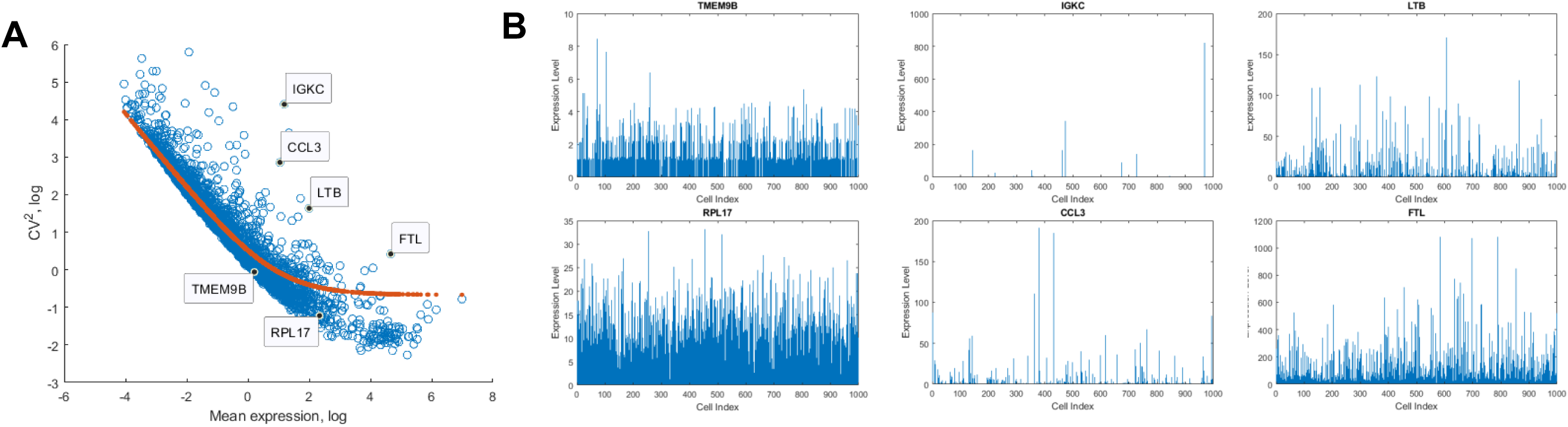
Identification of highly variable genes (HVGs). (**A**) The relationship between CV^2^ and mean expression of genes in LCL GM12878. The red line shows the trend for the GLM fit curve between CV^2^ and mean expression and used to identify HVGs. For each gene, the residual variability is calculated as the difference between observed CV^2^ and expected CV^2^ from the fitted curve. (**B**) Expression profiles of selected HVGs and lowly variable genes across cells. Cells are unsorted and remain a random order. Each vertical line is a cell and the height of line indicates the level of gene expression (UMI counts) in the cell.

### Cell-type origin determines the function of highly variable genes

To assess the biological functions of HVGs in different cell types, we performed the enrichment analyses (**Materials and Methods**). The categories of enriched gene ontology (GO) terms and pathways of the three cell types are largely distinct and reflect the respective cell functions of each cell type (**Table 1**). Two exemplary LCL HVGs are *CCL22* and *IFI27*. Collectively, more than expected number of LCL HVGs (FDR<0.01) are involved in *cytokine-* or *interferon-signaling pathways*, and also, more generally, *innate immune system*. LAEC HVGs, including genes, *COL1A1, MMP1*, and *IL17C*, are more likely to be involved in the processes of *collagen formation* and *extracellular matrix organization* (FDR<0.01 for both). DF HVGs, including *KRT14, ACAN,* and *FLG*, are more likely to be involved in *keratinization, regulation of cell proliferation*, as well as *extracellular structure organization* (FDR<0.01 for all). DF HVGs also include *SFRP2, DPP4,* and *LSP1*, which are marker genes defining major fibroblast subpopulations in human skin (Tabib et al. 2018). Taken together, these results show different cell types have different sets of HVGs, suggesting scEV implies cell function.

**Table 1.** Representative HVGs identified in the three cell types: LCL, LAEC, and DF, and the results of functional enrichment analyses. Genes are sorted by residual variability. The top 50 genes with the highest residual variability values are selected as representative HVGs.

| Cell type | Highly variable genes (HVGs), top 50 | Enriched GO terms, top 10 | Enriched Reactome pathways, top 10 |
| --- | --- | --- | --- |
| Lymphoblastoid Cell line (LCL) | *ANKRD37 ATF3 BIN1 BMP4 CAMP CCL22 CCL3 CCL3L3 CCL4 CCL4L2 CCR7 CD69 CD7 CD83 CDKN1A CTSC CYP1B1 DHRS9 DUSP2 FSCN1 HIST1H1C IER3 IFI27 IGHG1 IGHG3 IGHM IGKC ITM2A KCNMA1 LINC00176 LINC01588 LMNA LTA LTB MAL MIER2 MIR155HG MYC NFKBIA PMCH PRSS2 RGS1 RGS16 RGS2 RP11-291B21.2 S100A4 SFN TNFAIP2 TUBB4B WFDC2* | 1. signal transduction (GO:0007165)<br>2. response to stimulus (GO:0050896)<br>3. immune response (GO:0006955)<br>4. response to biotic stimulus (GO:0009607)<br>5. immune system process (GO:0002376)<br>6. response to external biotic stimulus (GO:0043207)<br>7. response to external stimulus (GO:0009605)<br>8. cytokine-mediated signaling pathway (GO:0019221)<br>9. defense response (GO:0006952)<br>10. response to chemical (GO:0042221) | 1. Immune System (R-HSA-168256)<br>2. Chemokine receptors bind chemokines (R-HSA-380108)<br>3. Interferon alpha/beta signaling (R-HSA-909733)<br>4. Cytokine Signaling in Immune system (R-HSA-1280215)<br>5. Interferon Signaling (R-HSA-913531)<br>6. Peptide ligand-binding receptors (R-HSA-375276)<br>7. G alpha (i) signalling events (R-HSA-418594)<br>8. Innate Immune System (R-HSA-168249)<br>9. Interferon gamma signaling (R-HSA-877300)<br>10. Cell Cycle (R-HSA-1640170) |
| Lung airway epithelial cell (LAEC) | *AMTN ANKRD1 AREG CCL2 CCL5 CCL7 COL1A1 COL1A2 COL3A1 COL6A1 COL6A3 CRCT1 CTGF CXCL5 CXCL6 FBXO32 GREM1 HAS2 IFNL1 IFNL2 IFNL3 IGFBP5 IGFL1 IL17C IL23A KRT14 KRT6B KRT81 LY6D MEG3 MMP1 MSMB OVOS2 PI3 POSTN PPBP RP11-338I21.1 S100A7 S100A8 S100A9 SERPINB2 SERPINB3 SERPINB4 SLC15A2 SPARC SUGCT SULF1 TEX26-AS1 TNFAIP6 TSLP* | 1. regulation of multicellular organismal process (GO:0051239)<br>2. regulation of signaling receptor activity (GO:0010469)<br>3. response to stimulus (GO:0050896)<br>4. regulation of cell proliferation (GO:0042127)<br>5. developmental process (GO:0032502)<br>6. extracellular matrix organization (GO:0030198)<br>7. response to chemical (GO:0042221)<br>8. response to organic substance (GO:0010033) | 1. Extracellular matrix organization (R-HSA-1474244)<br>2. Assembly of collagen fibrils and other multimeric structures (R-HSA-2022090)<br>3. Cytokine Signaling in Immune system (R-HSA-1280215)<br>4. Collagen formation (R-HSA-1474290)<br>5. Signaling by Interleukins (R-HSA-449147)<br>6. Chemokine receptors bind chemokines (R-HSA-380108)<br>7. Peptide ligand-binding receptors (R-HSA-375276)<br>8. Collagen biosynthesis and modifying enzymes (R-HSA-1650814) |
|  |  | 9. regulation of developmental process (GO:0050793)<br>10. regulation of response to stimulus (GO:0048583) | 9. Integrin cell surface interactions (R-HSA-216083)<br>10. Class A/1 (Rhodopsin-like receptors) (R-HSA-373076) |
| Dermal fibroblast (DF) | *ACAN ACTA2 ACTC1 CEMIP*<br>*CLU COMP CTSC CXCL1*<br>*DCN DKK1 FLG G0S2 GAL*<br>*HIST1H4C IGFBP5 IGFBP7*<br>*IL1RL1 KCNMA1 KRT14*<br>*KRT17 KRT19 KRT34 KRT81*<br>*KRTAP1-5 KRTAP2-3 LCE1F*<br>*LUM MGP MMP1 MMP3*<br>*MT1X NMB OLFM2 PCP4*<br>*PENK PGF PI16 POSTN*<br>*PPP1R14A PTTG1 PTX3*<br>*RARRES2 RGCC SCG5*<br>*SERPINE2 SFRP2 SFRP4*<br>*STMN2 TFPI2 TNFRSF11B* | 1. regulation of signaling receptor activity (GO:0010469)<br>2. developmental process (GO:0032502)<br>3. keratinization (GO:0031424)<br>4. anatomical structure development (GO:0048856)<br>5. regulation of cell proliferation (GO:0042127)<br>6. regulation of multicellular organismal process (GO:0051239)<br>7. extracellular matrix organization (GO:0030198)<br>8. extracellular structure organization (GO:0043062)<br>9. response to oxygen-containing compound (GO:1901700)<br>10. multicellular organismal process (GO:0032501) | 1. Extracellular matrix organization (R-HSA-1474244)<br>2. Regulation of Insulin-like Growth Factor (IGF) transport and uptake by Insulin-like Growth Factor Binding Proteins (IGFBPs) (R-HSA-381426)<br>3. ECM proteoglycans (R-HSA-3000178)<br>4. Hemostasis (R-HSA-109582)<br>5. Platelet degranulation (R-HSA-114608)<br>6. Dissolution of Fibrin Clot (R-HSA-75205)<br>7. Response to elevated platelet cytosolic Ca<sup>2+</sup> (R-HSA-76005)<br>8. Negative regulation of TCF-dependent signaling by WNT ligand antagonists (R-HSA-3772470)<br>9. GPCR ligand binding (R-HSA-500792)<br>10. Peptide ligand-binding receptors (R-HSA-375276) |

If two cell types have shared function, then we expect to see the overlap in their HVG-associated functions. This indeed is the case: there are overlaps between enriched functions between the three cell types we examined here. For example, *cytokine signaling pathway* is enriched for both LCL and LAEC, and *extracellular structure organization* is enriched for both LAEC and DF. Meanwhile, across all three cell types, there are 13 shared HVGs genes: *CDC20, CLEC2B, CLIC3, CTSC, HES1, MT1E, NPW, SOX4, STMN1, TK1, TRIB3*, and *UCHL1*, showing highly diverse cellular and molecular functions.

### HVGs as part of the regulatory network with high cell-type specificity

Next, we set out to test whether HVGs are co-expressed and thus tend to form co-expression networks (Mantsoki et al. 2016). We first imputed the expression matrix and then constructed the co-expressed network using the top 50 HVGs for each cell type. For LCLs, the network contains two main modules centered on *NFKBIA* and *IGHG1*, respectively (**Fig. 3A**). *NFKBIA* encodes NF-κB inhibitor that interacts with REL dimers to inhibit NF-κB/Rel complexes (Courtois et al. 2003; Lopez-Granados et al. 2008). For LAECs, two modules are centered on *IL23A*/*TNFAIP6* and *COL1A1* (**Fig. 3B**); for DF, *KRTAP2-3* and *IGFBP7* (**Fig. 3C**). Thus, functions of “hub” genes in HVG co-expression networks are closely relevant to the function of corresponding cell type. These results are another line of evidence that scEV implies cell function.

**Fig. 3.**
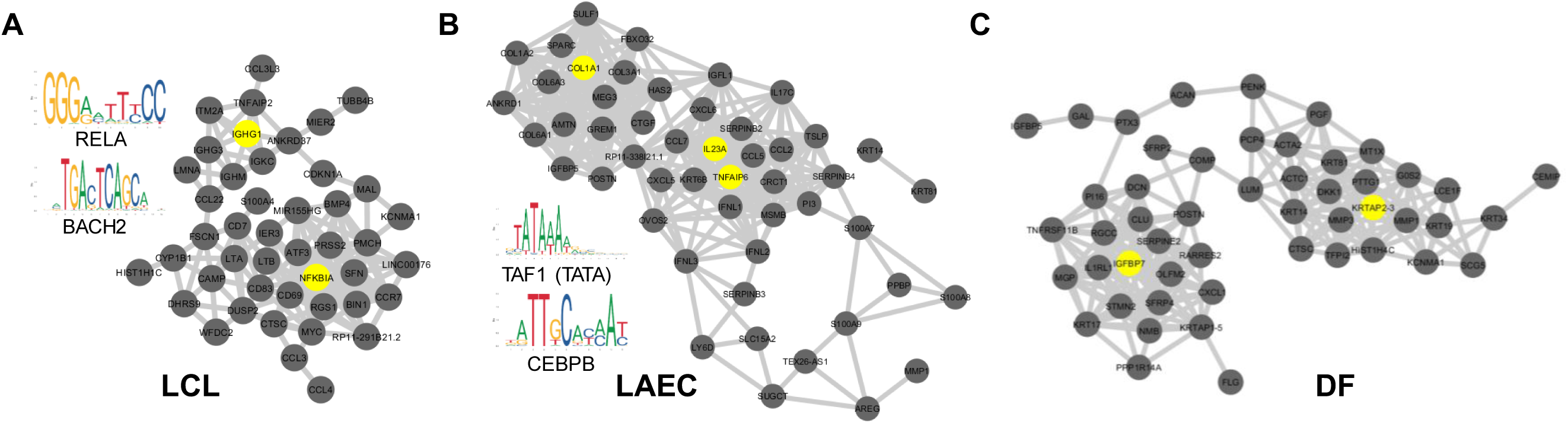
Co-expression networks of top HVGs. (**A**) Co-expression network between most-variable HVGs of LCL and two enriched binding motifs identified in these HVGs. (B) and (C) are for LAEC and DF, respectively.

The transcription of multiple HVGs may be involved in the same underlying regulatory activities, giving rise to the co-expression network as we observed. Thus, we wondered whether scEV in several different HVGs is driven by activities of one or few common TFs. To address this question, we searched for upstream regulators of the HVGs defined by our analysis (**Materials and Methods**). We identified significant enriched TF binding motifs upstream of HVGs, four for LCL and five for LAEC (**Supplementary Table S4**). No significantly enriched motif was identified for DF. The known motifs of LCL HVGs include that of NF-κB subunit gene, *RELA*, and that of *BACH2* (**Fig. 3A**). The known motifs of LAEC HVGs include TATA box and that of *CEBPB* (**Fig. 3B**).

To further explore the involvement of HVGs in cell type-specific regulatory network, we focused on LCL HVGs in a well-studied gene regulatory network that orchestrates B cell fate dynamics (Sciammas et al. 2011; Nutt et al. 2015; Roy et al. 2019). This known regulatory network involves eight genes, including three LCL HVGs: *PRDM1* (or Blimp-1), *AICDA* (or AID), and *IRF4*, two key regulatory genes with binding motifs enriched in targeting LCL HVGs (see above): *RELA* and *BACH2*, and three other key regulators: *BCL6, PAX5*, and *REL* (cRel)(**Fig. 4A**). We examined the inter-relationship between across-cell expressions of three LCL HVGs (**Fig. 4B**). The scatter plot shows that the directionality of the correlation between *AICDA* and *IRF4* depends on the expression level of *PRDM1*. Among cells with relatively low expression of *PRDM1*, expressions of *AICDA* and *IRF4* are negatively correlated. Whereas, among cells in which *PRDM1* is highly expressed, expressions of *AICDA* and *IRF4* are positively correlated. This nonlinear relationship between expressions of HVGs suggests they are embedded in a tightly regulated expression network. Thus, we examined the all-by-all correlation between expressions of all eight genes in this regulatory network using the imputed data of the homogenous LCLs (**Fig. 4C**). By comparing the sign of correlation coefficient of each pair of genes with the regulatory directionality of the gene pair in the model network, we found that the correlation matrix can be used to correctly recover 16 out of 18 direct regulatory relationships. The result suggests that, even in this highly homogenous population of LCLs, cells retain gene regulatory network activities that orchestrate cell fate dynamics as in their original B cells.

**Fig. 4.**
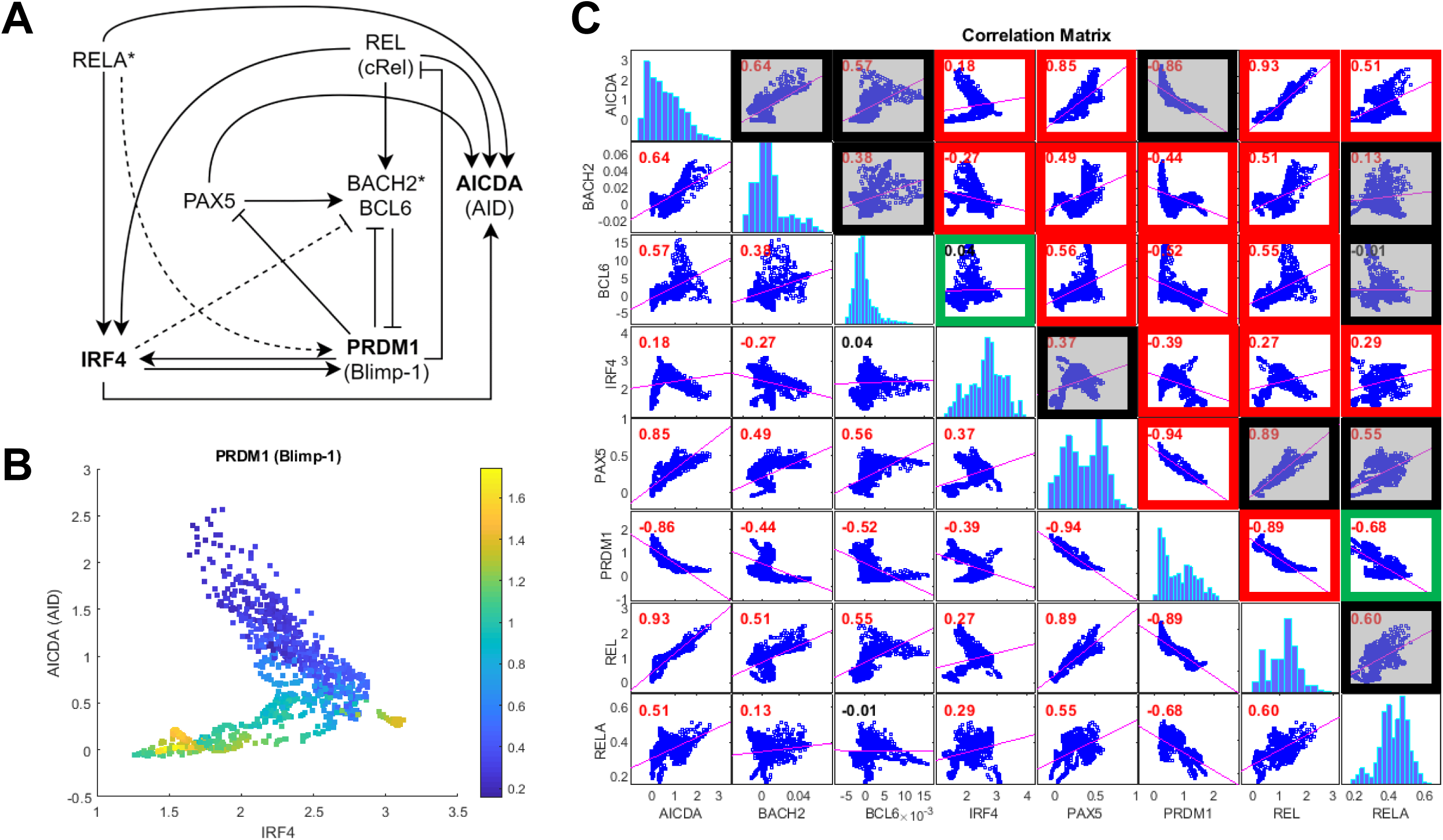
Gene regulatory network and correlation matrix of LCL HVGs. (**A**) NF-κB regulatory network model for activated B cell (ABC)-antibody secreting cell (ASC) differentiation, modified from (Roy et al. 2019). Bold font indicates HVGs; asterisk indicates the upstream TFs targeting HVGs; solid line dashed line indicates the regulatory relationship supported by the correlation between two corresponding genes, and the dashed line indicates regulatory relationship not supported by the expression correlation between genes. (**B**) Scatter plot of cells, showing the correlation between expression levels of three HVGs: *IRF4, AICDA* (AID) and *PRDM1* (Blimp-1). The color bar indicates the expression level of *PRDM1* (Blimp-1). (**C**) Correlation matrix between expression levels of eight genes involved in the model. Red boxes indicate that the direction of correlation between two genes is consistent with the direction of the relationship between the two in the regulatory model. Green boxes indicate inconsistency, while black boxes indicate no direct relationship according to the model. P-value in red indicated high significance (p < 0.01 after Bonferroni correction).

### Single cell expression variability is more similar between genetically related samples than unrelated samples

Given that scEV is important for cell function, we thought the level of scEV may be genetically determined. If true, then we expect that the similarity in scEV between cell lines derived from two genetically related individuals is higher than that between cell lines derived from two unrelated individuals. To test this, we performed the scRNA-seq with another LCL—GM18502, derived from a donor of African ancestry, unrelated to GM12878. We processed GM18502 along with GM12878 in the same batch (**Materials and Methods**), along with a technical replicate sample made from a 1:1 mixture of the two (Osorio et al. 2019). For comparison, we also obtained scRNA-seq data from the study of (Zhang et al. 2019) for another LCL GM12981. The donor of GM12891 is the father of GM12878. We estimated the scEV for these two additional scRNA-seq data sets: one from GM18502 (unrelated) and the other from GM12891 (father), using the same procedure applied to GM12878. To measure the correlation between scEV of different samples, we used both the Spearman correlation coefficient (SCC) and Pearson correlation coefficient (PCC). Across genes, the correlation between residual variability estimated from GM12878 and that from GM12891 (daughter-father) is ρ=0.92 (SCC) or r=0.94 (PCC)(**Supplementary Fig. S3**). That is to say, 85% (=r^2^) of the variance in scEV across genes of a daughter can be explained by that of a father. In contrast, only 77% of the variance can be explained by that of an unrelated individual—the correlation between GM12878 and GM18502 is ρ=0.87 (SCC) or r=0.89 (PCC)(**Supplementary Fig. S3**).

Note that, the similarity between these two related samples, GM12878 and GM12891 (daughter and father), might have been underestimated. This is due to the gender difference between the two samples was not taken into account. Furthermore, the two scRNA-seq data sets of them were produced in different batches: GM12878 by us in this study and GM12891 by (Zhang et al. 2019). The batch effect could also influence the daughter-father correlation downward. Nevertheless, we still observed a stronger correlation between the two related samples compared to that between the two unrelated samples. These results suggest that scEV is likely to be highly heritable.

When the correlation tests were performed across cell types, much weaker correlations were observed: the correlation between LCL and LAEC is 0.60 (SCC) or 0.65 (PCC), and that between LCL and DF is 0.57 (SCC) or 0.70 (PCC).

### Single cell expression variability is positively correlated with between-individual expression variability

Next, we examined the relationship between scEV and inter-individual expression variability. We distinguish between the two different types of variabilities at different organizational levels. Specifically, the former is cell-to-cell variability in a population of cells, and the latter is inter-individual variability at the human population level. We again focused on LCLs, for which population-scale gene expression data are available from the Geuvadis RNA-seq project of 1,000 Genomes samples. The bulk RNA-seq data was downloaded as normalized expression matrix of FPKM values. We retained data for all LCLs of European ancestry (CEU)(Lappalainen et al. 2013). With the residual variability estimated from scRNA-seq of GM12878 and that estimated from the CEU population, we tested the correlation between the two estimates across genes. When the test was conducted with all genes (n=8,424), we obtained a significant but weak positive correlation (SCC, ρ = 0.19, *p* = 1.2×10^-9^). We wondered whether this positive correlation was driven by subsets of genes. To identify these gene sets, we conducted the correlation tests for the GO-defined gene sets one by one. Across all gene sets tested, the average SCC for gene sets defined by GO biological process (BP) and molecular function (MF) terms are on average ρ=0.28 and ρ=0.23, respectively. Strikingly, we found a small number of gene sets that produced SCC much higher than averages. The functions of these gene sets include *B-cell activation involved in immune response* (GO: 0002322), cytokine receptor activity (GO:0004896), *cellular response to drug* (GO: 0035690) and *regulation of tyrosine phosphorylation of stat protein* (GO: 0042509)(**Fig. 5**), as well as *leukocyte chemotaxis* (GO: 0030595) and *phospholipase activity* (GO: 0004620)(for more examples, see **Supplementary Fig. S4**). Thus, for these gene sets, scEV may contribute to the establishment of between-individual expression variability.

**Fig. 5.**
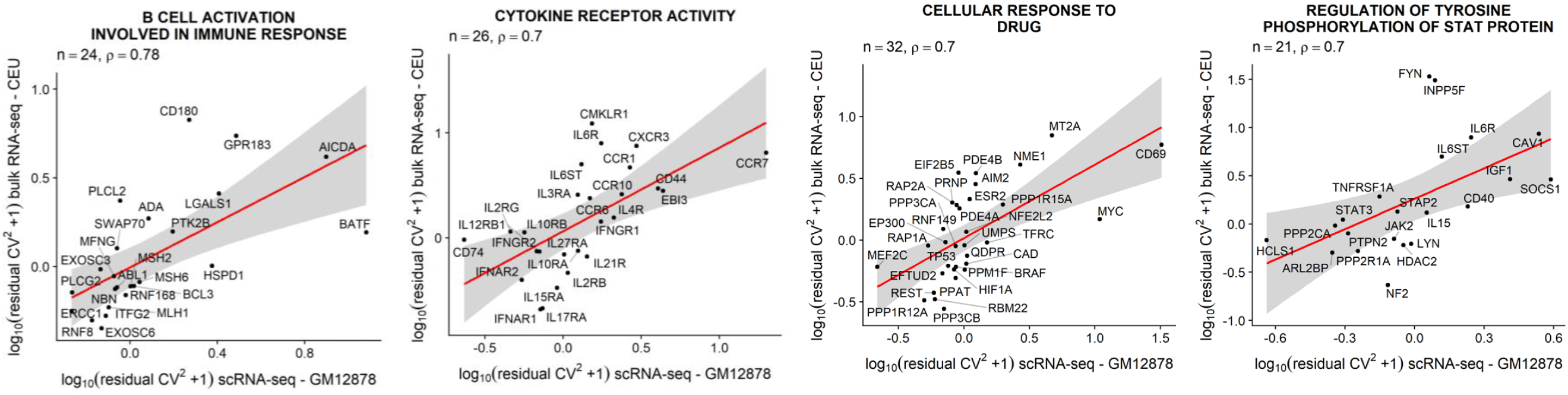
Correlation between scEV (i.e., residual variability estimated from LCL GM12878) and the population-level expression variability (measured in LCLs derived from unrelated individuals of European ancestry, CEU) between genes of selected gene sets. More examples can be found in **Supplementary Fig. S3**.

### No enriched functions associated with HVGs identified in human induced pluripotent stem cells (iPSCs)

Finally, we argued, if scEV is the indicator of cell type-specific function then scEV in undifferentiated cells should not be associated with any cellular functions. To test this, we obtained the scRNA-seq data from the study of (Nguyen et al. 2018)(see **Data Availability**). The data was generated from WTC-CRISPRi human iPSCs (Mandegar et al. 2016). Same as other cell types we examined in this study, these iPSCs were also prepared using the 10X Genomics Chromium controller. The released data contains five samples. We used the first batch (sample 1) of the data to perform the HVG detection and function enrichment tests, using the same procedure applied to other cell types. When plotting the relationship between log-transformed CV^2^ and log-transformed average expression (μ), we found almost no genes showing the deviated CV2 from the regression curve (**Supplementary Fig. S5**). That is, for the majority of genes, scEV can be explained by technical noise and sampling stochasticity. Or, in other words, iPSCs lack biological variability in single cell expression. Accordingly, we found no significant GO terms or enriched pathways associated with nearly random HVGs in iPSCs. This negative result is consistent with our prediction, based on the “variation is function” hypothesis, for iPSCs, which are not expected to be associated with any cellular function found in differentiated cells.

## Discussion

Single cell expression variability (or scEV) is also called gene expression noise, implying the stochastic nature of transcriptional activities in cells (Kaern et al. 2005; Raser and O’Shea 2005). Interrogating scEV data has provided insights into gene regulatory architecture (Mar et al. 2011; Chalancon et al. 2012), and manipulating the magnitude of scEV, through using noise enhancers or scEV-modulating chemicals, has been an approach to achieve drug synergies (Dar et al. 2014). Understanding the origin and functional implications of scEV has long been appreciated (Ko 1992; Fiering et al. 2000; Raj and van Oudenaarden 2008; Ecker et al. 2018).

In this study, we focused on scEV in human cells. More specifically, we wanted to characterize different expression levels of genes within a highly homogeneous population of genetically identical (or nearly isogenic) cells under the same environmental conditions. To this end, we set out to quantify scEV in highly homogeneous populations of a sizable number of viable cells. Working with cells of the same type, for example, LCL, we start by preprocessing data from thousands of cells. We found that, even though we have firstly preprocessed the data and retained only cells with similar library size and in the same cell cycle phase, it is not enough. There are still marked substructures, shown as branches of cells, in the embedding cloud of cells (**Fig. 1A**), as revealed by the new embedding algorithm (Moon et al. 2018). Retrospectively, we applied the trajectory analysis and found out that one of the longest branches contains cells with elevated expression of immunoglobulin genes (**Supplementary Fig. S5**).

Similarly, marked substructures were observed in the embedding plots of the other two cell types, LAEC and DF. Genes that were differentially expressed and drove the formation of branches of LAECs and DFs were different from those in LCL cells. Thus, there is no single or a small set of marker genes that can be used to capture cellular heterogeneity across different cell types, making the definition of populations of homogenous cells a tedious task. Our work might be the first focusing on comparing scEV in highly homogeneous cell populations across cell types.

We showed that scEV estimated from homogeneous populations of selected cells for different cell types carries information on cell type-specific function. Information on molecular functions of cells and biological processes of a given cell type can be extracted from a set of highly variable genes (HVGs), bearing significant biological meaning [see also (Dueck et al. 2015)]. HVGs detected in different cell types do not overlap and can reveal the subtle differences in cellar functions between cell types. These conclusions are reached based on our investigation of three cell types and their corresponding HVGs.

First, LCLs are usually established by in vitro infection of human peripheral blood lymphocytes with Epstein-Barr virus. The viral infection selectively immortalizes resting B cells, giving rise to an actively proliferating B cell population (Neitzel 1986). B cells genetically diversity by rearranging the immunoglobulin locus to produce diverse antibody repertories that allow the immune system to recognize foreign molecules and initiate differential immune responses (Tonegawa 1983; Papavasiliou et al. 1997; Mitchell et al. 2018). LCLs are produced through the rapid proliferation of few EBV-driven B cells from the blood cell population (Ryan et al. 2006). Thus, scRNA-seq data sets of LCLs offer a “snapshot” of highly diverse immunoglobulin rearrangement profiles in a much larger population of polyclonal B cells established in donors of these cell lines. Therefore, it is not unexpected to see quite a few immunoglobulin genes in the top list of HVGs identified in LCLs. In addition to these immunoglobulin genes, a number of other immune genes, especially C-C motif chemokine ligands (CCLs) and C-C motif chemokine receptors (CCRs), are in the list of HVGs of LCL. These genes play important roles in allowing the coordination of the activity of individual cells through intercellular communication, essential for the immune system maintains robustness (Altan-Bonnet and Mukherjee 2019). The HVG co-expression network analysis revealed the key role of the NF-κB pathway in facilitating communications between immune cells (Tay et al. 2010; Mitchell et al. 2018). More strikingly, we were able to reconstruct nearly entire NF-κB regulatory network, underlying differentiation of activated B cells and antibody-secreting cells, by using the correlation and anti-correlation relationships between expressions of HVGs and their regulatory genes.

Second, LAEC is a key cell type playing important roles in lung tissue remodeling, and pulmonary inflammatory and immune responses (Hiemstra et al. 2015). The airway epithelium, playing a critical role in conducting air to and from the alveoli, is a dynamic tissue that normally undergoes slow but constant turnover. In the event of mild to moderate injury, the airway epithelium responds vigorously to re-establish an epithelial sheet with normal structure and function. HVGs identified in LAECs, which are enriched with genes involved in collagen formation, regulation of cell proliferation, and extracellular matrix organization, accurately elucidate this aspect of functions of the airway epithelium. LAECs are also central to the defense of the lung against pathogens and particulates that are inhaled from the environment. This aspect of functions is also reflected in the enriched functionality of LAEC HVGs.

Third, DFs are responsible for generating connective tissue and play a critical role in normal wound healing (Tracy et al. 2016). DFs are also commonly used in immunological studies (Zhao et al. 2012; Ivashkiv and Donlin 2014; Hagai et al. 2018). HVGs identified in DFs again accurately reflect these primary aspects of DF functions, including extracellular matrix organization, keratinization, and regulation of signaling receptor activity. DF HVGs do have several categories of enriched functions overlap with those of LAEC, which is not unexpected, given that DF and LAEC have functional overlaps (Sacco et al. 2004).

Our results support the “variation is function” hypothesis, proposed by (Dueck et al. 2016), suggesting that the aggregate cellular function may depend on scEV. Dueck and colleagues also laid down several scenarios, including bet hedging, response distribution, fate plasticity, and so on, in which the establishment of the relationship between scEV and cell function could be attained. Our analytical framework using scRNA-seq data may be utilized in appropriate systems to test the plausibility of these different scenarios. If scEV is an accountable and credible surrogate of cell function, as we have shown in this study, then quantifying and characterizing scEV may become a first-line approach for understanding the function of cell types and tissues. Indeed, when we applied this framework to scRNA-seq data from human iPSCs, we observed no enriched gene functions and no regulatory pathways/networks associated with HVGs in iPSCs. This anti-example, showing no variation no function, further validates the “variation is function” hypothesis.

Through sequencing LCLs from three donors, we were able to compare the overall scEV across genes between related and unrelated LCL pairs. Our results suggest that scEV is a heritable trait and its relative magnitude across genes is genetically determined. In theory, the heritability of scEV can be estimated with more LCL samples from different donors of different levels of relatedness. A pairwise similarity matrix between LCLs in scEV can be regressed with the genetic relationship matrix between LCL donors, using a Haseman-Elston regression-type analysis (Haseman and Elston 1972), to quantify the heritability. The normalization between data from different LCLs (i.e., batch effect correction) can be achieved using the method of mutual nearest neighbors (Haghverdi et al. 2018), canonical correlation analysis (Butler et al. 2018) or manifold alignment (Amodio and Krishnaswamy 2018). With a population-scale scRNA-seq data set in the future, we will be able to identify mutations associated with increased scEV. At this moment, we still do not have such a resource of scRNA-seq data for samples from a sizable number of human individuals.

Nevertheless, we have shown that, across certain sets of genes, scEV is positively correlated with population-level expression variability. This correlation provides a new possibility to design single cell assays with one sample to approximate the population variability of certain genes’ expression. This new method may be used to study disease-causing expression dysregulation because it has been a number of cases that increased population-level expression variability has been linked with diseases (Ho et al. 2008; Li et al. 2010; Ecker et al. 2015; Guan et al. 2016; Huang et al. 2018).

In a visionary perspective article, Pelkmans (2012) pointed out that “Embracing this cell-to-cell variability as a fact in our scientific understanding requires a paradigm shift, but it will be necessary.” Indeed, scRNA-seq technologies have brought revolution to gene expression analysis. The technical development gives us a new approach beyond the capacity of traditional methods that rely on experimental measurements of population-average behavior of cells to conceive regulatory network models and signal processing pathways. More importantly, for traditional methods, by averaging information across many cells, differences among cells, which may be important in explaining mechanisms, can be lost. Given the large degree of cell-to-cell expression variability even between genetically homogeneous cells, conclusions reached as for such with traditional average-based methods may be of low-resolution, incomplete, and sometimes misleading (Tay et al. 2010; Bendall and Nolan 2012; Li and You 2013; Trapnell 2015). We have shown that scEV in highly homogeneous populations of human cells is widespread, is heritable, and implies cell function. We conclude that single cell variability and the information it contains are the key to a deepened understanding of cells and their functions. Careful assessment and characterization of cell-to-cell expression variability in relevant cell types will facilitate the study of normal cell functions as well as pathological cell processes.

## Materials and Methods

### LCL cell culture and scRNA-seq experiment

Two lymphoblastoid cell lines (LCLs), GM12878 (CEU) and GM18502 (YRI), were purchased from the Coriell Institute for Medical Research. They were cultured in the RPMI-1640 medium supplied with 2mM L-glutamine and 20% of non-inactivated fetal bovine serum, incubated at 37°C under 5% CO2 atmosphere. For maintenance, cells were subcultured every three days by adding fresh medium. For single cell sequencing, each cell line was subcultured with 200,000 viable cells/mL. To minimize the growth differences between those two cell lines, we plotted the growth curve by counting the viable cells which are not stained with 0.4% trypan blue every day. Both cell lines were harvested for single cell sample preparation and sequencing at day four (stationary phase) following the Sample Preparation Demonstrated Protocol and Single Cell 3’ Reagent Kits v2 User Guide provided by 10X Genomics. Briefly, cells were mixed well in each flask, and 1 mL of cell suspensions from each cell line were taken out. The cells were washed three times by centrifuging, suspending and resuspending in 1X PBS with 0.04% BSA. Viable cells were then counted using an automated cell counter (Thermo Fisher Scientific, Carlsbad, CA). Cells (∼5000 per cell line) were then pelleted and resuspended in the nuclease-free water based on cell suspension volume calculator table, followed by GEM (Gel Bead-In-Emulsions) generation and barcoding, the post GEM-RT cleanup, cDNA amplification, and library construction and sequencing. The experiments were conducted at the Texas A&M Institute for Genome Sciences and Society. The sequencing was conducted in the North Texas Genome Center facilities using a Novaseq 6000 sequencer (Illumina, San Diego, CA). Raw reads for each cell were analyzed using CellRanger (v2.0.0, 10X Genomics) and the outputs were aligned to the human reference genome (GRCh38) to obtain the counts.

### Non-LCL scRNA-seq data sets

The scRNA-seq data for lung airway epithelial cells (LAECs) was downloaded from the GEO database using accession number GSE115982. The original data was generated in the study of (Habiel et al. 2018) for CCR10^-^ and CCR10^+^ LAECs. We used the data generated from the CCR10^-^ cells with the sample identifier GSM3204305. The scRNA-seq data for primary dermal fibroblasts (DFs) was generated in the study of (Hagai et al. 2018). We downloaded the data for unstimulated DFs from the ArrayExpress database using accession number E-MTAB-5988. We also downloaded scRNA-seq data (GEO accession number GSE111912) generated in the study of (Zhang et al. 2019) for LCL sample GM12891. All of these data sets were produced using the 10X Genomics scRNA-seq solutions.

### Selection of highly homogeneous populations of cells

We used a supervised data analysis method to select highly homogeneous cells based on the scRNA-seq expression profile of each cell. The procedure is summarized in a flowchart (**Supplementary Fig. S1**). The main steps are as follows. We used Seurat (v0.2)(Butler et al. 2018) to assign each cell into a cell cycle phase and excluded cells that are not considered to be in G1-phase. We removed genes encoded in the mitochondrial genome from the analysis. We then selected and retained cells with library size between 50 and 95 percentiles. We used PHATE (Moon et al. 2018) to generate embedding plot of all remaining cells and inspected the distributions of cells in the three-dimensional plot, and manually picked one “core” cell. Finally, an additional 999 cells that are closest to the core cell, according to the Euclidean distances between cells, were selected to form the final 1,000-cell population. This selection procedure was applied to each of the three cell types independently.

### Identification of HVGs

Highly variable genes (HVG) is based on the assumption that genes with high variance relative to their mean expression are due to biological effects rather than just technical noise. We used the method proposed in (Brennecke et al. 2013), which is implemented in function `sc_hvg` of the scGEApp package (https://github.com/jamesjcai/scGEApp)(Cai 2019). This method starts by adjusting the library size and assumes that the observed mean expression 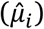 and the observed 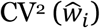 of gene *i* among cells have the following relationship:

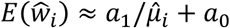

and

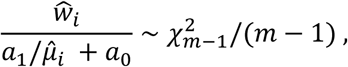

where *m* is the number of cells. The values of *a*_0_ and *a*_1_ are estimated by generalized linear regression (GLM). The residual term 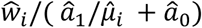 for each gene is used to test if the observed CV^2^ is significantly larger than the expected CV^2^ via a chi-squared test. Multiple testing p-value adjustment was performed by controlling the false discovery rate (FDR)(Benjamini and Hochberg 1995).

### Function enrichment analyses

To identify overrepresented biological functions of HVGs in different cell types, we performed the GO enrichment analysis using Enrichr (Chen et al. 2013; Kuleshov et al. 2016) and GOrilla (Eden et al. 2009). Enrichr was conducted for HVGs (FDR<0.01) against the rest of the expressed genes with respect to pathways collected in the Reactome pathway knowledgebase (Fabregat et al. 2018). GOrilla was performed with the list of genes sorted in descending order of their residual variability.

### Analyses of Co-expression network and regulatory regions of HVGs

MAGIC (van Dijk et al. 2018) was used to impute the expression matrix. The co-expression networks were constructed using SBEToolbox (Konganti et al. 2013). The motif analysis of the regulatory regions associated with the HVGs was performed using the GREAT (McLean et al. 2010). Genomic coordinates for the HVG genes from the Human Reference Genome (hg19) were downloaded from the Ensembl Biomart (Smedley et al. 2009) and converted into bed format using an in-house script. Identified motifs were searched against the JASPAR database (Khan et al. 2018) to match the binding sites of corresponding TFs.

## Data availability

The data sets produced in this study and computer code are available:

1. LCLs GM12878 and GM18502 scRNA-seq data in the Gene Expression Omnibus (GEO) database with accession number GSE126321: https://www.ncbi.nlm.nih.gov/geo/query/acc.cgi?acc=GSE126321
2. LCL GM12891 scRNA-seq data in the Gene Expression Omnibus (GEO) database with accession number GSE111912: https://www.ncbi.nlm.nih.gov/geo/query/acc.cgi?acc=GSE111912
3. LAEC scRNA-seq data in the Gene Expression Omnibus (GEO) database with accession number GSE115982: https://www.ncbi.nlm.nih.gov/geo/query/acc.cgi?acc=GSE115982
4. DF scRNA-seq data in ArrayExpress database with accession number E-MTAB-5988: https://www.ebi.ac.uk/arrayexpress/experiments/E-MTAB-5988
5. Human iPSC scRNA-seq data in ArrayExpress database with accession number E-MTAB-6687: https://www.ebi.ac.uk/arrayexpress/experiments/E-MTAB-6687
6. Computer codes used to analyze data: https://github.com/cailab-tamu/LCL_scRNA-seq

## Supplementary figure legends

**Fig. S1.** Flowchart of selection of a highly homogeneous population of cells.

**Fig. S2.** Correlation analyses of scEV between two genetically related LCL samples and between two genetically unrelated LCL samples.

**Fig. S3.** Correlation between scEV and population-level expression variability across genes of functional sets.

**Fig. S4.** PHATE 3-D embedding plot for cells colored according to *IGLC2* expression level in cells.

**Fig. S5.** The relationship between CV^2^ and mean expression of genes in human iPSCs.

## Acknowledgements

We thank Andrew Hillhouse and Chris Blazier for help with single cell preparation and raw data processing, Michael Criscitiello and Simon Mitchell for helpful discussions, and Barbara Gastel for writing suggestions.

## Author contributions

JH and JJC conceived and designed the project; DO and XY cultured the cells; DO, YZ, GL, PY, ES, JH, and JJC analyzed the data; DO and JJC wrote the manuscript. All authors reviewed the manuscript.

