## Supplementary material for "Extent, heritability, and functional relevance of single cell expression variability in highly homogeneous populations of human cells": Suppl Tables

### Single cell expression variability in highly homogeneous populations of human cells is widespread, heritable, and implies cell function

Daniel Osorio[1,#], Xue Yu[2,#], Yan Zhong[3,#], Guanxun Li[3], Peng Yu[4], Erchin Serpedin[4], Jianhua Huang[3,*], James J. Cai[1,4,5,*]

[1]Department of Veterinary Integrative Biosciences; [2]Department of Veterinary Pathobiology; [3]Department of Statistics; [4]Department of Electrical and Computer Engineering; [5]Interdisciplinary Program of Genetics; Texas A&M University, College Station, TX 77843, USA.

[#]These authors contributed equally to this work.

#### Supplementary Tables

**Supplementary Table S1. HVGs of LCL GM12878 (FDR<0.01).**

| id | genes | u | cv2 | residualcv2 | fitratio | pval | fdr |
|---|---|---|---|---|---|---|---|
| 1 | IGKC | 3.279132 | 81.83502 | 4.185526 | 65.72806 | 0 | 0 |
| 2 | WFDC2 | 0.145432 | 328.9569 | 3.611274 | 37.01318 | 0 | 0 |
| 3 | IGHM | 3.83006 | 38.27298 | 3.467405 | 32.05346 | 0 | 0 |
| 4 | CCL4 | 0.613502 | 76.75264 | 3.315936 | 27.54818 | 0 | 0 |
| 5 | CCL22 | 0.336603 | 112.033 | 3.249625 | 25.78067 | 0 | 0 |
| 6 | CCL4L2 | 0.248003 | 135.3981 | 3.189026 | 24.26478 | 0 | 0 |
| 7 | IGHG1 | 0.879858 | 35.99707 | 2.789436 | 16.27184 | 0 | 0 |
| 8 | RGS1 | 0.386765 | 52.73496 | 2.604946 | 13.5305 | 0 | 0 |
| 9 | CCL3 | 2.841433 | 17.42626 | 2.595852 | 13.40801 | 0 | 0 |
| 10 | CCL3L3 | 0.279217 | 65.97229 | 2.568705 | 13.04892 | 0 | 0 |
| 11 | BIN1 | 0.696029 | 27.70819 | 2.381194 | 10.81781 | 0 | 0 |
| 12 | LINC00176 | 0.90831 | 21.68926 | 2.30171 | 9.991251 | 0 | 0 |
| 13 | CD69 | 0.548075 | 27.29328 | 2.203893 | 9.060216 | 0 | 0 |
| 14 | CYP1B1 | 0.128323 | 83.99067 | 2.13275 | 8.438037 | 0 | 0 |
| 15 | KCNMA1 | 0.050237 | 198.4927 | 2.110967 | 8.256225 | 0 | 0 |
| 16 | PRSS2 | 0.044945 | 198.6256 | 2.004232 | 7.420392 | 0 | 0 |
| 17 | CD7 | 0.031185 | 277.0247 | 1.981667 | 7.254825 | 0 | 0 |
| 18 | IGHG3 | 0.637732 | 18.46284 | 1.9173 | 6.802569 | 0 | 0 |
| 19 | ATF3 | 0.060945 | 124.9399 | 1.833405 | 6.255147 | 0 | 0 |
| 20 | HIST1H1C | 5.572301 | 6.692024 | 1.806432 | 6.088682 | 0 | 0 |
| 21 | ITM2A | 0.688372 | 14.19716 | 1.705278 | 5.502913 | 0 | 0 |
| 22 | CAMP | 0.198917 | 36.10813 | 1.678864 | 5.359464 | 0 | 0 |
| 23 | DHRS9 | 0.18633 | 37.35663 | 1.655887 | 5.237722 | 0 | 0 |
| 24 | CCR7 | 1.774538 | 7.845156 | 1.624366 | 5.075202 | 0 | 0 |
| 25 | LTA | 0.267946 | 26.48256 | 1.621888 | 5.062638 | 0 | 0 |
| 26 | LTB | 7.381157 | 5.143045 | 1.590816 | 4.907754 | 0 | 0 |
| 27 | LINC01588 | 0.058874 | 96.39689 | 1.540991 | 4.669217 | 0 | 0 |

| | | | | | | | |
|---|---|---|---|---|---|---|---|
| 28 | TNFAIP2 | 0.055105 | 102.0179 | 1.534266 | 4.637921 | 0 | 0 |
| 29 | CD83 | 0.46288 | 15.67753 | 1.52757 | 4.606966 | 0 | 0 |
| 30 | RGS2 | 0.02954 | 181.5317 | 1.506009 | 4.5087 | 0 | 0 |
| 31 | SFN | 0.042664 | 124.2037 | 1.484344 | 4.412069 | 0 | 0 |
| 32 | MAL | 0.139676 | 40.37007 | 1.477027 | 4.379904 | 0 | 0 |
| 33 | DUSP2 | 1.468837 | 7.071851 | 1.436027 | 4.203959 | 0 | 0 |
| 34 | PMCH | 0.034682 | 137.8805 | 1.387602 | 4.005235 | 0 | 0 |
| 35 | LMNA | 0.693606 | 10.00946 | 1.36072 | 3.898999 | 0 | 0 |
| 36 | ANKRD37 | 1.588458 | 6.187685 | 1.338556 | 3.813533 | 6.91e-323 | 1.55e-320 |
| 37 | BMP4 | 0.145733 | 32.63714 | 1.30266 | 3.67907 | 1.73E-301 | 3.80E-299 |
| 38 | CTSC | 5.127163 | 4.096594 | 1.299318 | 3.666797 | 1.51E-299 | 3.22E-297 |
| 39 | NFKBIA | 2.071455 | 5.313563 | 1.297433 | 3.659889 | 1.86E-298 | 3.86E-296 |
| 40 | CDKN1A | 1.573795 | 5.831791 | 1.275124 | 3.579144 | 9.08E-286 | 1.84E-283 |
| 41 | RGS16 | 0.322842 | 16.02215 | 1.271473 | 3.5661 | 9.96E-284 | 1.96E-281 |
| 42 | MYC | 0.662088 | 9.439371 | 1.271459 | 3.566053 | 1.01E-283 | 1.96E-281 |
| 43 | IER3 | 0.190097 | 24.94214 | 1.269428 | 3.558817 | 1.37E-282 | 2.58E-280 |
| 44 | FSCN1 | 0.345521 | 14.96179 | 1.257065 | 3.515091 | 8.85E-276 | 1.63E-273 |
| 45 | IFI27 | 0.069111 | 61.65962 | 1.246983 | 3.479827 | 2.60E-270 | 4.70E-268 |
| 46 | S100A4 | 0.77263 | 8.282655 | 1.240495 | 3.457324 | 7.83E-267 | 1.38E-264 |
| 47 | MIR155HG | 1.409503 | 5.804966 | 1.218997 | 3.383793 | 1.57E-255 | 2.70E-253 |
| 48 | RP11-291B21.2 | 0.06744 | 61.24653 | 1.217002 | 3.377048 | 1.68E-254 | 2.85E-252 |
| 49 | MIER2 | 0.107068 | 38.38238 | 1.183488 | 3.265745 | 1.32E-237 | 2.19E-235 |
| 50 | TUBB4B | 0.758929 | 7.812086 | 1.170724 | 3.224327 | 2.22E-231 | 3.60E-229 |
| 51 | HSPB1 | 2.934978 | 4.132356 | 1.16681 | 3.211732 | 1.70E-229 | 2.71E-227 |
| 52 | TUBA1B | 4.204364 | 3.690298 | 1.151269 | 3.162204 | 4.11E-222 | 6.41E-220 |
| 53 | ENTPD2 | 0.03457 | 109.0478 | 1.149859 | 3.157747 | 1.89E-221 | 2.89E-219 |
| 54 | RSPH1 | 0.050121 | 74.62172 | 1.130425 | 3.096971 | 1.80E-212 | 2.71E-210 |
| 55 | DUSP4 | 0.186136 | 21.86224 | 1.119224 | 3.062478 | 2.07E-207 | 3.06E-205 |
| 56 | TIMD4 | 0.727041 | 7.534822 | 1.107223 | 3.025944 | 4.43E-202 | 6.41E-200 |
| 57 | FAM83D | 0.04374 | 82.9143 | 1.104333 | 3.01721 | 8.23E-201 | 1.17E-198 |

| | | | | | | | |
|---|---|---|---|---|---|---|---|
| 58 | IL4I1 | 0.302539 | 13.97623 | 1.082447 | 2.951895 | 2.24E-191 | 3.14E-189 |
| 59 | RARRES2 | 0.032028 | 107.9551 | 1.065316 | 2.901757 | 3.33E-184 | 4.58E-182 |
| 60 | PLK3 | 0.099306 | 36.12493 | 1.053114 | 2.866564 | 3.30E-179 | 4.46E-177 |
| 61 | MARCKSL1 | 1.653733 | 4.523364 | 1.043223 | 2.838351 | 3.16E-175 | 4.20E-173 |
| 62 | MACROD2 | 0.37391 | 11.24497 | 1.033409 | 2.81063 | 2.46E-171 | 3.21E-169 |
| 63 | ATP1B1 | 0.067846 | 49.24275 | 1.004572 | 2.730739 | 3.06E-160 | 3.94E-158 |
| 64 | SGK1 | 0.083907 | 40.25531 | 1.003899 | 2.728902 | 5.47E-160 | 6.94E-158 |
| 65 | TNF | 0.232107 | 16.02188 | 0.9988 | 2.715022 | 4.43E-158 | 5.53E-156 |
| 66 | C10orf99 | 0.100097 | 33.59288 | 0.987821 | 2.685377 | 5.05E-154 | 6.21E-152 |
| 67 | LINC01480 | 1.180651 | 5.030214 | 0.986607 | 2.682118 | 1.40E-153 | 1.70E-151 |
| 68 | IGLV6-57 | 0.364215 | 10.85912 | 0.978009 | 2.659158 | 1.86E-150 | 2.22E-148 |
| 69 | HMOX1 | 0.652307 | 7.096395 | 0.976255 | 2.654497 | 7.96E-150 | 9.36E-148 |
| 70 | IL32 | 0.128028 | 26.16786 | 0.964484 | 2.623433 | 1.24E-145 | 1.44E-143 |
| 71 | BATF | 0.915177 | 5.665865 | 0.963788 | 2.621608 | 2.19E-145 | 2.50E-143 |
| 72 | STEAP1 | 0.028884 | 106.7261 | 0.952881 | 2.59317 | 1.41E-141 | 1.59E-139 |
| 73 | INSIG1 | 0.747742 | 6.33211 | 0.951267 | 2.588988 | 5.11E-141 | 5.68E-139 |
| 74 | EGR2 | 0.103258 | 31.40006 | 0.949158 | 2.583534 | 2.73E-140 | 2.99E-138 |
| 75 | AICDA | 0.760627 | 6.219602 | 0.944171 | 2.570682 | 1.40E-138 | 1.51E-136 |
| 76 | HMGB2 | 1.294716 | 4.568666 | 0.937745 | 2.554216 | 2.13E-136 | 2.27E-134 |
| 77 | S100A10 | 0.217619 | 15.88443 | 0.935194 | 2.547709 | 1.54E-135 | 1.63E-133 |
| 78 | LGALS14 | 0.458972 | 8.518501 | 0.911315 | 2.487591 | 1.17E-127 | 1.21E-125 |
| 79 | NPIPB15 | 0.082273 | 36.99746 | 0.901016 | 2.462103 | 2.34E-124 | 2.40E-122 |
| 80 | PRKCDBP | 0.034718 | 83.96069 | 0.892571 | 2.441398 | 1.08E-121 | 1.10E-119 |
| 81 | ANXA1 | 0.121821 | 25.18031 | 0.880649 | 2.412465 | 5.42E-118 | 5.43E-116 |
| 82 | CLC | 0.041854 | 69.08249 | 0.879142 | 2.408831 | 1.57E-117 | 1.55E-115 |
| 83 | RP5-887A10.1 | 0.867276 | 5.284174 | 0.862083 | 2.368089 | 2.22E-112 | 2.17E-110 |
| 84 | BCL2A1 | 0.630299 | 6.444295 | 0.856842 | 2.35571 | 7.92E-111 | 7.65E-109 |
| 85 | TNFRSF18 | 0.059105 | 47.90335 | 0.845448 | 2.329021 | 1.68E-107 | 1.60E-105 |
| 86 | GPR34 | 0.070685 | 39.89794 | 0.833053 | 2.300332 | 5.88E-104 | 5.55E-102 |
| 87 | ID1 | 0.034407 | 79.10255 | 0.824199 | 2.280053 | 1.80E-101 | 1.68E-99 |

| | | | | | | | |
|---|---|---|---|---|---|---|---|
| 88 | STAG3 | 0.586888 | 6.543555 | 0.82344 | 2.278323 | 2.92E-101 | 2.69E-99 |
| 89 | HES1 | 0.064485 | 42.71778 | 0.814068 | 2.25707 | 1.12E-98 | 1.02E-96 |
| 90 | ZFP36L1 | 1.507996 | 3.737536 | 0.81063 | 2.249324 | 9.72E-98 | 8.76E-96 |
| 91 | HEY1 | 0.054765 | 49.16239 | 0.798318 | 2.221801 | 1.98E-94 | 1.77E-92 |
| 92 | NAALADL2-AS2 | 0.173094 | 16.67251 | 0.784356 | 2.190995 | 9.16E-91 | 8.08E-89 |
| 93 | B4GALT1 | 0.985705 | 4.522424 | 0.78136 | 2.184442 | 5.45E-90 | 4.75E-88 |
| 94 | CXCR4 | 0.055106 | 46.7748 | 0.754478 | 2.1265 | 3.13E-83 | 2.70E-81 |
| 95 | STMN1 | 1.487363 | 3.543647 | 0.750941 | 2.118993 | 2.29E-82 | 1.96E-80 |
| 96 | PHGDH | 0.207348 | 13.74494 | 0.748943 | 2.114763 | 7.01E-82 | 5.92E-80 |
| 97 | ID2 | 0.398587 | 8.000331 | 0.7423 | 2.100761 | 2.80E-80 | 2.34E-78 |
| 98 | JUNB | 1.379893 | 3.631655 | 0.739704 | 2.095315 | 1.17E-79 | 9.68E-78 |
| 99 | T | 0.017515 | 140.4125 | 0.735502 | 2.086529 | 1.16E-78 | 9.53E-77 |
| 100 | EGR3 | 0.028054 | 86.98154 | 0.719775 | 2.05397 | 5.37E-75 | 4.36E-73 |
| 101 | BIRC3 | 0.763761 | 4.931221 | 0.714648 | 2.043468 | 7.95E-74 | 6.39E-72 |
| 102 | ICAM1 | 0.158202 | 16.76304 | 0.709931 | 2.03385 | 9.28E-73 | 7.38E-71 |
| 103 | TCL1A | 0.919032 | 4.37611 | 0.707958 | 2.029842 | 2.58E-72 | 2.03E-70 |
| 104 | CHAC1 | 0.088846 | 28.31453 | 0.705677 | 2.025217 | 8.34E-72 | 6.51E-70 |
| 105 | SOCS3 | 0.068952 | 35.86295 | 0.702881 | 2.019562 | 3.50E-71 | 2.70E-69 |
| 106 | SOX4 | 0.364761 | 8.227901 | 0.701705 | 2.017189 | 6.38E-71 | 4.88E-69 |
| 107 | GZMH | 0.064235 | 38.22001 | 0.699103 | 2.011946 | 2.40E-70 | 1.82E-68 |
| 108 | FRMD4A | 0.388751 | 7.756317 | 0.692124 | 1.997954 | 8.07E-69 | 6.06E-67 |
| 109 | ABCB9 | 0.14963 | 17.29215 | 0.691171 | 1.996052 | 1.30E-68 | 9.67E-67 |
| 110 | CEBPD | 0.083832 | 29.30821 | 0.685677 | 1.985115 | 1.99E-67 | 1.47E-65 |
| 111 | DUSP11 | 0.588895 | 5.684222 | 0.685008 | 1.983787 | 2.77E-67 | 2.03E-65 |
| 112 | NFKBID | 0.131666 | 19.17556 | 0.679074 | 1.972052 | 5.09E-66 | 3.69E-64 |
| 113 | SELL | 0.612654 | 5.493063 | 0.677893 | 1.969724 | 9.04E-66 | 6.49E-64 |
| 114 | JUN | 0.274586 | 10.08388 | 0.676611 | 1.9672 | 1.69E-65 | 1.20E-63 |
| 115 | TRAF1 | 0.195168 | 13.28034 | 0.662095 | 1.93885 | 1.74E-62 | 1.23E-60 |
| 116 | CYS1 | 0.048282 | 48.38157 | 0.661089 | 1.9369 | 2.79E-62 | 1.95E-60 |
| 117 | PSAT1 | 0.300337 | 9.211261 | 0.659581 | 1.933982 | 5.67E-62 | 3.93E-60 |

| | | | | | | | |
|---|---|---|---|---|---|---|---|
| 118 | CHI3L2 | 1.043576 | 3.877033 | 0.659498 | 1.933822 | 5.89E-62 | 4.05E-60 |
| 119 | COCH | 0.276512 | 9.750708 | 0.648785 | 1.913214 | 8.44E-60 | 5.75E-58 |
| 120 | NAALADL2 | 0.066898 | 34.9208 | 0.647502 | 1.910762 | 1.52E-59 | 1.03E-57 |
| 121 | C1orf162 | 0.05587 | 41.27384 | 0.642583 | 1.901385 | 1.42E-58 | 9.53E-57 |
| 122 | H2AFZ | 7.158372 | 1.997749 | 0.640523 | 1.897474 | 3.60E-58 | 2.39E-56 |
| 123 | TUBB2A | 0.076712 | 30.10846 | 0.628996 | 1.875726 | 6.08E-56 | 4.01E-54 |
| 124 | AQP3 | 0.178457 | 13.84027 | 0.625075 | 1.868386 | 3.38E-55 | 2.21E-53 |
| 125 | MT2A | 0.819732 | 4.262495 | 0.612959 | 1.845885 | 6.21E-53 | 4.03E-51 |
| 126 | HERPUD1 | 5.862162 | 2.006135 | 0.611159 | 1.842567 | 1.33E-52 | 8.58E-51 |
| 127 | PTTG1 | 1.32013 | 3.263185 | 0.610943 | 1.842168 | 1.46E-52 | 9.33E-51 |
| 128 | CCL5 | 0.271863 | 9.502606 | 0.608996 | 1.838585 | 3.32E-52 | 2.10E-50 |
| 129 | CD24 | 0.182669 | 13.31199 | 0.606653 | 1.834282 | 8.88E-52 | 5.59E-50 |
| 130 | TNFAIP3 | 0.197931 | 12.34135 | 0.600985 | 1.823914 | 9.40E-51 | 5.87E-49 |
| 131 | VIM | 6.387519 | 1.949397 | 0.59757 | 1.817697 | 3.84E-50 | 2.38E-48 |
| 132 | GADD45A | 0.25067 | 10.02283 | 0.594656 | 1.812408 | 1.27E-49 | 7.78E-48 |
| 133 | IFIT1 | 0.489237 | 5.921364 | 0.594499 | 1.812123 | 1.35E-49 | 8.24E-48 |
| 134 | VPREB3 | 0.216376 | 11.29908 | 0.589665 | 1.803384 | 9.60E-49 | 5.81E-47 |
| 135 | EBI3 | 0.609183 | 4.978926 | 0.57575 | 1.778464 | 2.42E-46 | 1.45E-44 |
| 136 | PHLDA3 | 0.258031 | 9.557647 | 0.571356 | 1.770667 | 1.34E-45 | 7.97E-44 |
| 137 | HSPA5 | 8.422 | 1.806379 | 0.563253 | 1.756376 | 2.99E-44 | 1.77E-42 |
| 138 | STXBP6 | 0.054501 | 38.78829 | 0.556666 | 1.744846 | 3.59E-43 | 2.11E-41 |
| 139 | HAVCR2 | 0.060402 | 35.10514 | 0.555374 | 1.742592 | 5.82E-43 | 3.40E-41 |
| 140 | MS4A7 | 0.164012 | 13.90695 | 0.555249 | 1.742375 | 6.09E-43 | 3.53E-41 |
| 141 | NCALD | 0.160436 | 14.12009 | 0.550849 | 1.734724 | 3.12E-42 | 1.80E-40 |
| 142 | CDC20 | 0.172901 | 13.1933 | 0.549319 | 1.732073 | 5.49E-42 | 3.14E-40 |
| 143 | PLAC8 | 4.792886 | 1.962381 | 0.549259 | 1.73197 | 5.61E-42 | 3.18E-40 |
| 144 | IFIT3 | 0.686568 | 4.458841 | 0.545405 | 1.725306 | 2.30E-41 | 1.30E-39 |
| 145 | LAMP5 | 0.312625 | 7.93308 | 0.542675 | 1.720604 | 6.21E-41 | 3.47E-39 |
| 146 | GADD45B | 0.724297 | 4.284333 | 0.540222 | 1.716388 | 1.51E-40 | 8.37E-39 |
| 147 | ALOX5AP | 0.612989 | 4.780059 | 0.539232 | 1.714689 | 2.15E-40 | 1.19E-38 |

| | | | | | | | |
|---|---|---|---|---|---|---|---|
| 148 | TK1 | 0.344248 | 7.296137 | 0.53599 | 1.70914 | 6.86E-40 | 3.76E-38 |
| 149 | C1orf54 | 0.065172 | 32.00274 | 0.535365 | 1.708072 | 8.58E-40 | 4.67E-38 |
| 150 | ACY3 | 0.175732 | 12.79252 | 0.5328 | 1.703696 | 2.13E-39 | 1.15E-37 |
| 151 | WDR91 | 0.116351 | 18.54178 | 0.532515 | 1.70321 | 2.36E-39 | 1.27E-37 |
| 152 | IL2RA | 0.024912 | 80.65077 | 0.527782 | 1.695169 | 1.25E-38 | 6.65E-37 |
| 153 | CAV1 | 0.216287 | 10.62081 | 0.527405 | 1.694529 | 1.42E-38 | 7.54E-37 |
| 154 | DERL3 | 1.309743 | 3.009852 | 0.526194 | 1.692478 | 2.17E-38 | 1.14E-36 |
| 155 | FTL | 105.8097 | 1.525237 | 0.525995 | 1.692142 | 2.32E-38 | 1.22E-36 |
| 156 | PIM3 | 0.337883 | 7.316136 | 0.523931 | 1.688653 | 4.76E-38 | 2.47E-36 |
| 157 | DDIT4 | 1.142262 | 3.200818 | 0.517052 | 1.677076 | 5.05E-37 | 2.61E-35 |
| 158 | PRKAR1B | 0.20494 | 10.98053 | 0.514306 | 1.672477 | 1.28E-36 | 6.59E-35 |
| 159 | TRAF4 | 0.670467 | 4.379074 | 0.511735 | 1.668183 | 3.06E-36 | 1.56E-34 |
| 160 | SOCS1 | 0.657208 | 4.432823 | 0.510691 | 1.666442 | 4.34E-36 | 2.20E-34 |
| 161 | SLC25A4 | 1.979342 | 2.458911 | 0.509052 | 1.663713 | 7.51E-36 | 3.79E-34 |
| 162 | BIRC5 | 0.381354 | 6.552188 | 0.508578 | 1.662925 | 8.80E-36 | 4.41E-34 |
| 163 | ASB2 | 0.149661 | 14.35829 | 0.505427 | 1.657693 | 2.51E-35 | 1.25E-33 |
| 164 | LINC00158 | 0.076616 | 26.60064 | 0.503946 | 1.65524 | 4.09E-35 | 2.02E-33 |
| 165 | RP11-598F7.3 | 0.025176 | 77.81272 | 0.502301 | 1.652519 | 7.03E-35 | 3.46E-33 |
| 166 | FABP5 | 3.002173 | 2.105804 | 0.499577 | 1.648024 | 1.71E-34 | 8.38E-33 |
| 167 | CDKN2C | 1.41632 | 2.791193 | 0.48907 | 1.630799 | 5.05E-33 | 2.45E-31 |
| 168 | CCNB1 | 0.153231 | 13.80474 | 0.487236 | 1.627811 | 9.02E-33 | 4.36E-31 |
| 169 | C16orf74 | 0.253173 | 8.923557 | 0.486813 | 1.627123 | 1.03E-32 | 4.95E-31 |
| 170 | RP11-166B2.1 | 0.106839 | 19.12912 | 0.485115 | 1.624362 | 1.76E-32 | 8.40E-31 |
| 171 | MGST1 | 0.523172 | 5.047482 | 0.48316 | 1.62119 | 3.25E-32 | 1.54E-30 |
| 172 | NINJ1 | 0.184756 | 11.64312 | 0.482659 | 1.620377 | 3.80E-32 | 1.79E-30 |
| 173 | PIM1 | 0.096864 | 20.8155 | 0.478676 | 1.613936 | 1.31E-31 | 6.15E-30 |
| 174 | CCR1 | 0.121371 | 16.89937 | 0.478481 | 1.613621 | 1.39E-31 | 6.50E-30 |
| 175 | S100A6 | 1.204257 | 2.988829 | 0.476375 | 1.610227 | 2.67E-31 | 1.24E-29 |
| 176 | UBE2S | 1.144377 | 3.050675 | 0.469995 | 1.599986 | 1.86E-30 | 8.60E-29 |
| 177 | CLIC3 | 0.102744 | 19.51465 | 0.468895 | 1.598227 | 2.60E-30 | 1.19E-28 |

| | | | | | | | |
|---|---|---|---|---|---|---|---|
| 178 | LARP6 | 0.030293 | 62.67941 | 0.467226 | 1.595561 | 4.29E-30 | 1.96E-28 |
| 179 | RHOB | 0.037854 | 50.16877 | 0.46175 | 1.586849 | 2.19E-29 | 9.95E-28 |
| 180 | ENDOD1 | 0.061682 | 31.27341 | 0.459815 | 1.58378 | 3.89E-29 | 1.75E-27 |
| 181 | NCF2 | 0.171829 | 12.10841 | 0.45802 | 1.580941 | 6.58E-29 | 2.95E-27 |
| 182 | DNASE1L3 | 0.110007 | 18.02134 | 0.452447 | 1.572155 | 3.32E-28 | 1.48E-26 |
| 183 | TNFSF10 | 0.270406 | 8.142165 | 0.450035 | 1.568366 | 6.65E-28 | 2.95E-26 |
| 184 | IGF1 | 0.239267 | 9.012927 | 0.449243 | 1.567126 | 8.35E-28 | 3.68E-26 |
| 185 | HRASLS2 | 0.494781 | 5.025736 | 0.438689 | 1.550673 | 1.64E-26 | 7.17E-25 |
| 186 | RHOV | 0.022894 | 80.15174 | 0.438618 | 1.550564 | 1.67E-26 | 7.28E-25 |
| 187 | CENPF | 0.101166 | 19.16418 | 0.436409 | 1.547141 | 3.08E-26 | 1.33E-24 |
| 188 | ETV7 | 0.161138 | 12.53508 | 0.43567 | 1.545999 | 3.77E-26 | 1.63E-24 |
| 189 | LDHA | 4.687084 | 1.760075 | 0.435634 | 1.545944 | 3.81E-26 | 1.63E-24 |
| 190 | LMO2 | 0.073233 | 25.90068 | 0.434568 | 1.544296 | 5.11E-26 | 2.18E-24 |
| 191 | RASGRP2 | 0.456649 | 5.29038 | 0.431206 | 1.539113 | 1.28E-25 | 5.44E-24 |
| 192 | CLEC4C | 0.077406 | 24.39077 | 0.426895 | 1.532492 | 4.11E-25 | 1.74E-23 |
| 193 | GBP2 | 0.155324 | 12.82726 | 0.425933 | 1.531018 | 5.32E-25 | 2.24E-23 |
| 194 | SPARC | 0.119677 | 16.22031 | 0.424603 | 1.528984 | 7.59E-25 | 3.17E-23 |
| 195 | TRAC | 1.003201 | 3.13008 | 0.42336 | 1.527084 | 1.06E-24 | 4.40E-23 |
| 196 | HHEX | 0.231107 | 9.041972 | 0.423054 | 1.526616 | 1.15E-24 | 4.75E-23 |
| 197 | EGR1 | 0.047577 | 38.59595 | 0.420914 | 1.523354 | 2.02E-24 | 8.33E-23 |
| 198 | NPW | 0.021841 | 82.43198 | 0.420402 | 1.522573 | 2.32E-24 | 9.49E-23 |
| 199 | IFI30 | 0.112906 | 17.01675 | 0.419054 | 1.520522 | 3.30E-24 | 1.35E-22 |
| 200 | MSL3 | 0.621533 | 4.1966 | 0.418454 | 1.51961 | 3.87E-24 | 1.57E-22 |
| 201 | NLRP7 | 0.081491 | 22.99596 | 0.41649 | 1.516629 | 6.46E-24 | 2.61E-22 |
| 202 | LGMN | 0.183227 | 10.97084 | 0.415903 | 1.515739 | 7.53E-24 | 3.03E-22 |
| 203 | PRCAT47 | 0.064778 | 28.55266 | 0.415514 | 1.515149 | 8.34E-24 | 3.33E-22 |
| 204 | TBX21 | 0.128008 | 15.1052 | 0.414845 | 1.514136 | 9.92E-24 | 3.94E-22 |
| 205 | FEZ1 | 0.111439 | 17.14611 | 0.414581 | 1.513736 | 1.06E-23 | 4.20E-22 |
| 206 | P2RX5 | 0.689984 | 3.896822 | 0.413928 | 1.512748 | 1.26E-23 | 4.96E-22 |
| 207 | MID1IP1 | 0.143308 | 13.59738 | 0.411973 | 1.509794 | 2.08E-23 | 8.17E-22 |

| | | | | | | | |
|---|---|---|---|---|---|---|---|
| 208 | OASL | 0.076402 | 24.23655 | 0.408233 | 1.504157 | 5.43E-23 | 2.12E-21 |
| 209 | KIAA0101 | 1.959446 | 2.23137 | 0.40792 | 1.503687 | 5.88E-23 | 2.28E-21 |
| 210 | GGH | 0.375685 | 5.988059 | 0.406935 | 1.502207 | 7.56E-23 | 2.92E-21 |
| 211 | ING1 | 0.131184 | 14.65497 | 0.40688 | 1.502124 | 7.66E-23 | 2.95E-21 |
| 212 | NAPSA | 0.824178 | 3.447388 | 0.404041 | 1.497865 | 1.57E-22 | 6.01E-21 |
| 213 | C16orf54 | 0.519249 | 4.669217 | 0.399882 | 1.491649 | 4.44E-22 | 1.69E-20 |
| 214 | SSPN | 0.086291 | 21.41606 | 0.399092 | 1.490471 | 5.41E-22 | 2.05E-20 |
| 215 | ZFP36L2 | 1.909533 | 2.228711 | 0.396327 | 1.486355 | 1.07E-21 | 4.04E-20 |
| 216 | MKI67 | 0.220156 | 9.101972 | 0.388271 | 1.474429 | 7.58E-21 | 2.85E-19 |
| 217 | WNT10A | 0.135055 | 14.00791 | 0.388119 | 1.474205 | 7.86E-21 | 2.94E-19 |
| 218 | NETO2 | 0.173001 | 11.21829 | 0.387666 | 1.473537 | 8.76E-21 | 3.26E-19 |
| 219 | H1FX | 0.475053 | 4.916984 | 0.387137 | 1.472758 | 9.94E-21 | 3.68E-19 |
| 220 | AC074289.1 | 0.60414 | 4.141616 | 0.385945 | 1.471004 | 1.32E-20 | 4.87E-19 |
| 221 | MT1E | 0.161488 | 11.89233 | 0.384964 | 1.469562 | 1.67E-20 | 6.13E-19 |
| 222 | SULF2 | 0.45236 | 5.081989 | 0.384019 | 1.468173 | 2.09E-20 | 7.63E-19 |
| 223 | HMX2 | 0.064445 | 27.74095 | 0.381772 | 1.464877 | 3.55E-20 | 1.29E-18 |
| 224 | RRM2 | 0.297716 | 7.018564 | 0.38058 | 1.463133 | 4.70E-20 | 1.70E-18 |
| 225 | CTA-384D8.34 | 0.765568 | 3.521795 | 0.379523 | 1.461587 | 6.02E-20 | 2.17E-18 |
| 226 | CCDC180 | 0.069647 | 25.63462 | 0.376643 | 1.457384 | 1.18E-19 | 4.22E-18 |
| 227 | CENPW | 0.371408 | 5.849752 | 0.374662 | 1.4545 | 1.86E-19 | 6.64E-18 |
| 228 | STAC3 | 0.182702 | 10.55001 | 0.374272 | 1.453932 | 2.03E-19 | 7.23E-18 |
| 229 | RP11-47L3.1 | 0.048145 | 36.3838 | 0.373353 | 1.452597 | 2.51E-19 | 8.89E-18 |
| 230 | H1F0 | 0.028551 | 60.4163 | 0.372528 | 1.451399 | 3.03E-19 | 1.07E-17 |
| 231 | ID3 | 9.208026 | 1.473999 | 0.371436 | 1.449815 | 3.89E-19 | 1.37E-17 |
| 232 | ORC6 | 0.283242 | 7.199161 | 0.365213 | 1.440821 | 1.58E-18 | 5.54E-17 |
| 233 | SEPP1 | 0.018414 | 92.28358 | 0.365179 | 1.440773 | 1.59E-18 | 5.55E-17 |
| 234 | S100A11 | 1.326346 | 2.540749 | 0.363032 | 1.437682 | 2.57E-18 | 8.92E-17 |
| 235 | CITED2 | 0.286484 | 7.114511 | 0.362729 | 1.437247 | 2.75E-18 | 9.50E-17 |
| 236 | EFNA5 | 0.051838 | 33.48108 | 0.361395 | 1.43533 | 3.69E-18 | 1.27E-16 |
| 237 | P3H2 | 0.155521 | 11.99528 | 0.360007 | 1.43334 | 5.01E-18 | 1.72E-16 |

| | | | | | | | |
|---|---|---|---|---|---|---|---|
| 238 | IL10 | 0.113198 | 15.98726 | 0.359025 | 1.431933 | 6.22E-18 | 2.12E-16 |
| 239 | IL4R | 0.138878 | 13.25822 | 0.358376 | 1.431003 | 7.17E-18 | 2.43E-16 |
| 240 | ATF5 | 0.893849 | 3.133712 | 0.357613 | 1.429912 | 8.47E-18 | 2.86E-16 |
| 241 | TPRG1 | 0.162031 | 11.53163 | 0.357153 | 1.429254 | 9.36E-18 | 3.15E-16 |
| 242 | CMTM7 | 0.06435 | 27.01145 | 0.35371 | 1.424342 | 1.97E-17 | 6.61E-16 |
| 243 | TXNIP | 1.161736 | 2.689571 | 0.352011 | 1.421924 | 2.84E-17 | 9.48E-16 |
| 244 | RGCC | 0.297523 | 6.788475 | 0.34672 | 1.414421 | 8.73E-17 | 2.90E-15 |
| 245 | CRIP1 | 1.520919 | 2.339655 | 0.346158 | 1.413626 | 9.82E-17 | 3.25E-15 |
| 246 | CDKN3 | 0.244854 | 7.973275 | 0.34615 | 1.413615 | 9.84E-17 | 3.25E-15 |
| 247 | MYL9 | 0.042361 | 39.84131 | 0.340424 | 1.405543 | 3.24E-16 | 1.06E-14 |
| 248 | MAT2A | 0.331387 | 6.162673 | 0.336904 | 1.400605 | 6.65E-16 | 2.18E-14 |
| 249 | CCND2 | 0.465026 | 4.738286 | 0.33443 | 1.397144 | 1.10E-15 | 3.58E-14 |
| 250 | CXCR3 | 1.389463 | 2.409114 | 0.332629 | 1.39463 | 1.58E-15 | 5.12E-14 |
| 251 | SNHG15 | 0.35679 | 5.784948 | 0.332119 | 1.393919 | 1.75E-15 | 5.65E-14 |
| 252 | SERPINB1 | 0.120226 | 14.70177 | 0.330498 | 1.391661 | 2.42E-15 | 7.78E-14 |
| 253 | MYBL2 | 0.527942 | 4.29147 | 0.327373 | 1.387319 | 4.48E-15 | 1.44E-13 |
| 254 | TROAP | 0.143708 | 12.45624 | 0.326833 | 1.38657 | 4.99E-15 | 1.59E-13 |
| 255 | SERPINB9 | 0.172158 | 10.58543 | 0.325285 | 1.384425 | 6.76E-15 | 2.15E-13 |
| 256 | LRRK2 | 0.057138 | 29.4105 | 0.325206 | 1.384316 | 6.86E-15 | 2.17E-13 |
| 257 | PLD4 | 0.255021 | 7.527558 | 0.322776 | 1.380956 | 1.10E-14 | 3.48E-13 |
| 258 | TYW3 | 0.937556 | 2.941808 | 0.322488 | 1.380558 | 1.16E-14 | 3.66E-13 |
| 259 | DDIT3 | 0.531173 | 4.240014 | 0.319651 | 1.376648 | 2.01E-14 | 6.30E-13 |
| 260 | GMNN | 0.260644 | 7.36133 | 0.318663 | 1.375287 | 2.43E-14 | 7.58E-13 |
| 261 | HIST1H2BJ | 0.304336 | 6.476414 | 0.318061 | 1.374461 | 2.72E-14 | 8.47E-13 |
| 262 | PLA2G16 | 0.188759 | 9.687871 | 0.317574 | 1.373791 | 2.99E-14 | 9.26E-13 |
| 263 | CCDC74A | 0.27521 | 7.01014 | 0.314905 | 1.370129 | 4.96E-14 | 1.53E-12 |
| 264 | RARRES3 | 0.744102 | 3.360024 | 0.314475 | 1.36954 | 5.37E-14 | 1.65E-12 |
| 265 | CLEC2B | 0.103312 | 16.58268 | 0.311196 | 1.365056 | 9.92E-14 | 3.04E-12 |
| 266 | ADAM19 | 0.339408 | 5.889239 | 0.310551 | 1.364176 | 1.12E-13 | 3.41E-12 |
| 267 | CD44 | 1.874008 | 2.060936 | 0.310393 | 1.363961 | 1.15E-13 | 3.50E-12 |

| | | | | | | | |
|---|---|---|---|---|---|---|---|
| 268 | TYMS | 0.440555 | 4.798071 | 0.306822 | 1.359099 | 2.22E-13 | 6.73E-12 |
| 269 | PTCRA | 0.029281 | 54.95829 | 0.302529 | 1.353277 | 4.85E-13 | 1.46E-11 |
| 270 | S1PR2 | 0.27225 | 6.984789 | 0.302343 | 1.353025 | 5.01E-13 | 1.51E-11 |
| 271 | BNIP3 | 0.815303 | 3.131733 | 0.301359 | 1.351694 | 5.98E-13 | 1.79E-11 |
| 272 | ZC3H12A | 0.123546 | 13.90203 | 0.299475 | 1.34915 | 8.38E-13 | 2.50E-11 |
| 273 | CD22 | 0.538987 | 4.101396 | 0.296772 | 1.345508 | 1.35E-12 | 4.02E-11 |
| 274 | TUBB2B | 0.025851 | 61.67063 | 0.295761 | 1.344149 | 1.62E-12 | 4.79E-11 |
| 275 | RP3-428L16.2 | 0.069451 | 23.69647 | 0.295351 | 1.343597 | 1.74E-12 | 5.13E-11 |
| 276 | ISYNA1 | 0.036379 | 44.09157 | 0.293997 | 1.341779 | 2.20E-12 | 6.47E-11 |
| 277 | LINC01531 | 0.026813 | 59.3933 | 0.293948 | 1.341714 | 2.22E-12 | 6.50E-11 |
| 278 | GPR183 | 1.552441 | 2.199494 | 0.293808 | 1.341526 | 2.28E-12 | 6.64E-11 |
| 279 | CRELD2 | 5.273557 | 1.488956 | 0.29289 | 1.340295 | 2.67E-12 | 7.76E-11 |
| 280 | NR6A1 | 0.07672 | 21.50155 | 0.29241 | 1.339652 | 2.90E-12 | 8.41E-11 |
| 281 | PIM2 | 1.575018 | 2.181354 | 0.292098 | 1.339234 | 3.06E-12 | 8.85E-11 |
| 282 | ABCA6 | 0.065909 | 24.75805 | 0.289397 | 1.335622 | 4.88E-12 | 1.40E-10 |
| 283 | SRM | 2.786531 | 1.743174 | 0.287396 | 1.332952 | 6.87E-12 | 1.97E-10 |
| 284 | GIMAP7 | 0.176153 | 9.98532 | 0.287164 | 1.332643 | 7.14E-12 | 2.04E-10 |
| 285 | SETP10 | 0.261747 | 7.103375 | 0.286511 | 1.331772 | 7.98E-12 | 2.27E-10 |
| 286 | PCID2 | 0.417661 | 4.890591 | 0.285749 | 1.330758 | 9.08E-12 | 2.58E-10 |
| 287 | SLC2A5 | 0.283454 | 6.644428 | 0.285643 | 1.330617 | 9.25E-12 | 2.61E-10 |
| 288 | MIR3142HG | 0.047236 | 33.90387 | 0.284354 | 1.328904 | 1.15E-11 | 3.24E-10 |
| 289 | ITGB2-AS1 | 0.047324 | 33.786 | 0.282686 | 1.326689 | 1.52E-11 | 4.27E-10 |
| 290 | SELM | 0.604004 | 3.731001 | 0.281382 | 1.324959 | 1.89E-11 | 5.28E-10 |
| 291 | WARS | 1.051316 | 2.644119 | 0.280867 | 1.324278 | 2.06E-11 | 5.74E-10 |
| 292 | LMTK2 | 0.096063 | 17.1918 | 0.279679 | 1.322706 | 2.50E-11 | 6.96E-10 |
| 293 | RP3-395M20.12 | 0.066651 | 24.20264 | 0.277371 | 1.319655 | 3.66E-11 | 1.01E-09 |
| 294 | DMKN | 0.047765 | 33.27008 | 0.276236 | 1.318159 | 4.40E-11 | 1.22E-09 |
| 295 | MKNK2 | 0.759758 | 3.183846 | 0.273832 | 1.314994 | 6.50E-11 | 1.79E-09 |
| 296 | RASSF6 | 0.041744 | 37.77582 | 0.272963 | 1.313852 | 7.48E-11 | 2.05E-09 |
| 297 | RASSF4 | 0.035527 | 44.17664 | 0.272855 | 1.31371 | 7.61E-11 | 2.08E-09 |

| | | | | | | | |
|---|---|---|---|---|---|---|---|
| 298 | PRDM1 | 0.644431 | 3.522823 | 0.267813 | 1.307102 | 1.70E-10 | 4.62E-09 |
| 299 | SAMSN1 | 1.224011 | 2.404877 | 0.267425 | 1.306596 | 1.80E-10 | 4.90E-09 |
| 300 | LDLRAD4 | 0.234995 | 7.62901 | 0.267292 | 1.306422 | 1.84E-10 | 4.98E-09 |
| 301 | DUSP10 | 0.290515 | 6.390739 | 0.266887 | 1.305893 | 1.96E-10 | 5.29E-09 |
| 302 | TRIB3 | 0.319711 | 5.90341 | 0.26521 | 1.303705 | 2.55E-10 | 6.86E-09 |
| 303 | ZBTB32 | 0.317389 | 5.936587 | 0.264954 | 1.303371 | 2.66E-10 | 7.11E-09 |
| 304 | LINC00926 | 0.254169 | 7.123154 | 0.264758 | 1.303115 | 2.74E-10 | 7.31E-09 |
| 305 | HMGN2 | 7.576107 | 1.359058 | 0.263839 | 1.301919 | 3.16E-10 | 8.40E-09 |
| 306 | PMAIP1 | 1.807479 | 1.99637 | 0.263558 | 1.301553 | 3.30E-10 | 8.75E-09 |
| 307 | HNRNPH1 | 1.21034 | 2.408447 | 0.263094 | 1.300949 | 3.54E-10 | 9.36E-09 |
| 308 | CD180 | 0.277111 | 6.617096 | 0.262887 | 1.30068 | 3.66E-10 | 9.64E-09 |
| 309 | ZWINT | 0.266005 | 6.842029 | 0.262451 | 1.300112 | 3.91E-10 | 1.03E-08 |
| 310 | SPSB1 | 0.059494 | 26.54162 | 0.261247 | 1.298549 | 4.71E-10 | 1.23E-08 |
| 311 | HMMR | 0.083075 | 19.32586 | 0.260726 | 1.297872 | 5.10E-10 | 1.33E-08 |
| 312 | MDM2 | 0.425232 | 4.698544 | 0.25927 | 1.295984 | 6.37E-10 | 1.66E-08 |
| 313 | TMEM51 | 0.0585 | 26.91249 | 0.259011 | 1.295649 | 6.62E-10 | 1.72E-08 |
| 314 | NKG7 | 0.283712 | 6.46387 | 0.258839 | 1.295425 | 6.80E-10 | 1.76E-08 |
| 315 | PAQR7 | 0.052783 | 29.67548 | 0.258091 | 1.294457 | 7.61E-10 | 1.96E-08 |
| 316 | BBC3 | 0.150664 | 11.13071 | 0.2568 | 1.292786 | 9.25E-10 | 2.38E-08 |
| 317 | C1orf186 | 0.464827 | 4.374272 | 0.254178 | 1.289402 | 1.37E-09 | 3.50E-08 |
| 318 | APOL1 | 0.350096 | 5.423372 | 0.252671 | 1.28746 | 1.71E-09 | 4.37E-08 |
| 319 | ERN1 | 0.31412 | 5.910346 | 0.252187 | 1.286836 | 1.84E-09 | 4.68E-08 |
| 320 | RP11-138A9.2 | 0.197457 | 8.722265 | 0.251825 | 1.286371 | 1.94E-09 | 4.92E-08 |
| 321 | PHF19 | 0.452447 | 4.451851 | 0.251778 | 1.286311 | 1.95E-09 | 4.94E-08 |
| 322 | SDF2L1 | 9.953629 | 1.292634 | 0.249488 | 1.283369 | 2.73E-09 | 6.89E-08 |
| 323 | TRBC2 | 0.52028 | 4.007111 | 0.248381 | 1.281948 | 3.21E-09 | 8.06E-08 |
| 324 | TBC1D30 | 0.02749 | 55.31664 | 0.247284 | 1.280543 | 3.76E-09 | 9.42E-08 |
| 325 | TLCD1 | 0.048234 | 32.01589 | 0.24724 | 1.280487 | 3.79E-09 | 9.45E-08 |
| 326 | GNG3 | 0.082051 | 19.28027 | 0.246705 | 1.279801 | 4.09E-09 | 1.02E-07 |
| 327 | OSR2 | 0.07772 | 20.26367 | 0.245353 | 1.278073 | 4.97E-09 | 1.23E-07 |

| | | | | | | | |
|---|---|---|---|---|---|---|---|
| 328 | BCAS1 | 0.034081 | 44.69735 | 0.244087 | 1.276455 | 5.95E-09 | 1.47E-07 |
| 329 | FCRL5 | 0.188663 | 9.005044 | 0.244039 | 1.276394 | 5.99E-09 | 1.48E-07 |
| 330 | RAB37 | 0.074813 | 20.93907 | 0.242121 | 1.273948 | 7.87E-09 | 1.93E-07 |
| 331 | AC104024.1 | 0.07801 | 20.09804 | 0.240658 | 1.272086 | 9.67E-09 | 2.37E-07 |
| 332 | SAMD10 | 0.225556 | 7.67493 | 0.23844 | 1.269267 | 1.32E-08 | 3.22E-07 |
| 333 | CAPN10-AS1 | 0.180558 | 9.298014 | 0.23758 | 1.268177 | 1.48E-08 | 3.62E-07 |
| 334 | RP6-159A1.4 | 0.090128 | 17.47114 | 0.236263 | 1.266507 | 1.78E-08 | 4.33E-07 |
| 335 | IRF4 | 2.116037 | 1.822407 | 0.235509 | 1.265553 | 1.97E-08 | 4.78E-07 |
| 336 | CCNB2 | 0.168205 | 9.873941 | 0.235156 | 1.265107 | 2.07E-08 | 5.01E-07 |
| 337 | UNC13C | 0.027744 | 54.12548 | 0.234504 | 1.264281 | 2.27E-08 | 5.46E-07 |
| 338 | TOX2 | 0.047796 | 31.86032 | 0.23356 | 1.263089 | 2.58E-08 | 6.19E-07 |
| 339 | CLDN14 | 0.040711 | 37.00644 | 0.228111 | 1.256224 | 5.36E-08 | 1.28E-06 |
| 340 | RP11-511B23.1 | 0.022967 | 64.63706 | 0.226614 | 1.254346 | 6.53E-08 | 1.56E-06 |
| 341 | PTP4A3 | 0.184372 | 8.988018 | 0.222016 | 1.248592 | 1.19E-07 | 2.83E-06 |
| 342 | LBH | 0.354835 | 5.201236 | 0.221438 | 1.24787 | 1.28E-07 | 3.04E-06 |
| 343 | C1orf228 | 0.076184 | 20.15095 | 0.220918 | 1.247221 | 1.37E-07 | 3.24E-06 |
| 344 | TSC22D3 | 0.202342 | 8.261061 | 0.218697 | 1.244455 | 1.82E-07 | 4.28E-06 |
| 345 | IFIT2 | 0.173193 | 9.458911 | 0.218059 | 1.243661 | 1.97E-07 | 4.63E-06 |
| 346 | OAS3 | 0.073362 | 20.81269 | 0.217532 | 1.243005 | 2.11E-07 | 4.94E-06 |
| 347 | A4GALT | 0.129111 | 12.30088 | 0.217289 | 1.242704 | 2.17E-07 | 5.08E-06 |
| 348 | LMO7 | 0.086689 | 17.75294 | 0.215826 | 1.240887 | 2.61E-07 | 6.08E-06 |
| 349 | S1PR4 | 2.037246 | 1.81232 | 0.215305 | 1.24024 | 2.78E-07 | 6.47E-06 |
| 350 | CASP1 | 0.076087 | 20.03963 | 0.214177 | 1.238842 | 3.20E-07 | 7.43E-06 |
| 351 | UCHL1 | 0.514124 | 3.90098 | 0.213012 | 1.237399 | 3.70E-07 | 8.55E-06 |
| 352 | RGS3 | 0.142748 | 11.18005 | 0.212704 | 1.237018 | 3.84E-07 | 8.86E-06 |
| 353 | SPIB | 0.862988 | 2.761272 | 0.210077 | 1.233774 | 5.30E-07 | 1.22E-05 |
| 354 | MYH10 | 0.022644 | 64.46674 | 0.210037 | 1.233724 | 5.33E-07 | 1.22E-05 |
| 355 | TNFSF4 | 0.019291 | 75.46696 | 0.209855 | 1.233499 | 5.45E-07 | 1.25E-05 |
| 356 | MNDA | 0.744246 | 3.018694 | 0.207475 | 1.230567 | 7.26E-07 | 1.66E-05 |
| 357 | IPCEF1 | 0.10439 | 14.80309 | 0.207288 | 1.230337 | 7.43E-07 | 1.69E-05 |

| | | | | | | | |
|---|---|---|---|---|---|---|---|
| 358 | KCNQ1OT1 | 0.288871 | 6.048929 | 0.207273 | 1.230318 | 7.44E-07 | 1.69E-05 |
| 359 | RP11-1151B14.4 | 0.103325 | 14.93821 | 0.206872 | 1.229825 | 7.81E-07 | 1.76E-05 |
| 360 | IER2 | 1.231438 | 2.255063 | 0.206223 | 1.229027 | 8.44E-07 | 1.90E-05 |
| 361 | ZFP36 | 2.594001 | 1.643485 | 0.205095 | 1.227642 | 9.65E-07 | 2.17E-05 |
| 362 | LST1 | 0.074144 | 20.3479 | 0.204985 | 1.227507 | 9.78E-07 | 2.19E-05 |
| 363 | TNFSF14 | 0.185232 | 8.798172 | 0.204745 | 1.227212 | 1.01E-06 | 2.25E-05 |
| 364 | RAB13 | 0.940303 | 2.610178 | 0.204584 | 1.227015 | 1.03E-06 | 2.29E-05 |
| 365 | NME8 | 0.055862 | 26.63093 | 0.204278 | 1.226639 | 1.06E-06 | 2.36E-05 |
| 366 | CD1C | 0.02866 | 50.86385 | 0.204163 | 1.226498 | 1.08E-06 | 2.39E-05 |
| 367 | RP4-555D20.2 | 0.036283 | 40.39424 | 0.203839 | 1.226101 | 1.12E-06 | 2.48E-05 |
| 368 | CBX6 | 0.809157 | 2.848714 | 0.201974 | 1.223817 | 1.39E-06 | 3.07E-05 |
| 369 | MGLL | 0.690025 | 3.152212 | 0.20191 | 1.223738 | 1.40E-06 | 3.09E-05 |
| 370 | SQLE | 0.57565 | 3.55706 | 0.200513 | 1.222029 | 1.65E-06 | 3.62E-05 |
| 371 | ARHGAP24 | 0.044679 | 32.87635 | 0.199825 | 1.221189 | 1.79E-06 | 3.91E-05 |
| 372 | ANP32E | 0.550862 | 3.6608 | 0.198504 | 1.219577 | 2.08E-06 | 4.54E-05 |
| 373 | RP11-138I18.2 | 0.103733 | 14.75873 | 0.198437 | 1.219496 | 2.10E-06 | 4.57E-05 |
| 374 | BCL2L1 | 0.221488 | 7.488833 | 0.198354 | 1.219394 | 2.12E-06 | 4.60E-05 |
| 375 | QPRT | 1.032586 | 2.456413 | 0.197231 | 1.218025 | 2.41E-06 | 5.21E-05 |
| 376 | CCL17 | 0.019474 | 73.78878 | 0.196692 | 1.217369 | 2.56E-06 | 5.53E-05 |
| 377 | IRF8 | 1.073633 | 2.4012 | 0.196081 | 1.216626 | 2.75E-06 | 5.91E-05 |
| 378 | NLK | 0.135637 | 11.51444 | 0.195989 | 1.216514 | 2.78E-06 | 5.96E-05 |
| 379 | FCGR2A | 0.070126 | 21.2508 | 0.195599 | 1.216039 | 2.90E-06 | 6.21E-05 |
| 380 | NAB2 | 0.10616 | 14.40449 | 0.195549 | 1.215978 | 2.92E-06 | 6.22E-05 |
| 381 | CHPF | 0.418929 | 4.458511 | 0.195546 | 1.215975 | 2.92E-06 | 6.22E-05 |
| 382 | NOP16 | 0.535848 | 3.718889 | 0.194731 | 1.214984 | 3.20E-06 | 6.80E-05 |
| 383 | SH3TC1 | 0.149007 | 10.56121 | 0.194362 | 1.214536 | 3.34E-06 | 7.07E-05 |
| 384 | GPT2 | 0.073485 | 20.29654 | 0.194003 | 1.214099 | 3.47E-06 | 7.34E-05 |
| 385 | LRRC75A | 0.148419 | 10.59186 | 0.193708 | 1.213741 | 3.59E-06 | 7.56E-05 |
| 386 | BEX5 | 0.16353 | 9.712392 | 0.193645 | 1.213666 | 3.61E-06 | 7.58E-05 |
| 387 | PLG | 0.071025 | 20.95442 | 0.19364 | 1.213659 | 3.62E-06 | 7.58E-05 |

| | | | | | | | |
|---|---|---|---|---|---|---|---|
| 388 | LEUTX | 0.042039 | 34.62616 | 0.192737 | 1.212564 | 4.00E-06 | 8.37E-05 |
| 389 | C2CD2L | 0.063747 | 23.20026 | 0.192637 | 1.212443 | 4.04E-06 | 8.44E-05 |
| 390 | LIPH | 0.192782 | 8.390735 | 0.19223 | 1.211949 | 4.23E-06 | 8.81E-05 |
| 391 | CCL25 | 0.321597 | 5.46099 | 0.192032 | 1.21171 | 4.33E-06 | 8.98E-05 |
| 392 | KCTD1 | 0.063944 | 23.08245 | 0.190493 | 1.209846 | 5.13E-06 | 0.000106 |
| 393 | HCK | 0.118925 | 12.90572 | 0.19023 | 1.209528 | 5.28E-06 | 0.000109 |
| 394 | TCL1B | 0.049975 | 29.21629 | 0.189911 | 1.209142 | 5.47E-06 | 0.000113 |
| 395 | NGFRAP1 | 0.061395 | 23.96011 | 0.188992 | 1.208031 | 6.05E-06 | 0.000124 |
| 396 | GPER1 | 0.028028 | 51.17853 | 0.188507 | 1.207446 | 6.38E-06 | 0.000131 |
| 397 | UPP1 | 0.061379 | 23.93328 | 0.187626 | 1.206382 | 7.02E-06 | 0.000143 |
| 398 | TNFSF11 | 0.024817 | 57.55387 | 0.186612 | 1.205159 | 7.83E-06 | 0.00016 |
| 399 | BATF3 | 0.055038 | 26.52765 | 0.186146 | 1.204599 | 8.23E-06 | 0.000167 |
| 400 | CCR10 | 1.062348 | 2.38981 | 0.185513 | 1.203836 | 8.81E-06 | 0.000179 |
| 401 | MIAT | 0.129774 | 11.85153 | 0.184733 | 1.202897 | 9.58E-06 | 0.000194 |
| 402 | CCR6 | 0.109526 | 13.81517 | 0.182609 | 1.200345 | 1.20E-05 | 0.000242 |
| 403 | RP11-111A22.1 | 0.086653 | 17.13928 | 0.180259 | 1.197527 | 1.54E-05 | 0.000309 |
| 404 | GPR82 | 0.077213 | 19.08673 | 0.179335 | 1.196422 | 1.69E-05 | 0.000339 |
| 405 | FAM46C | 0.207171 | 7.779921 | 0.179077 | 1.196113 | 1.74E-05 | 0.000348 |
| 406 | RP1-34B20.21 | 0.099667 | 14.99524 | 0.177237 | 1.193915 | 2.10E-05 | 0.000419 |
| 407 | CLSPN | 0.047436 | 30.32488 | 0.176873 | 1.19348 | 2.18E-05 | 0.000434 |
| 408 | LAMP3 | 0.143473 | 10.7304 | 0.176217 | 1.192697 | 2.33E-05 | 0.000463 |
| 409 | C3orf58 | 0.147515 | 10.45248 | 0.174976 | 1.191218 | 2.64E-05 | 0.000524 |
| 410 | TRIB1 | 0.242509 | 6.761171 | 0.173142 | 1.189035 | 3.18E-05 | 0.000629 |
| 411 | HIST1H2BC | 0.174421 | 8.984401 | 0.172828 | 1.188661 | 3.28E-05 | 0.000647 |
| 412 | CDCA7 | 0.152983 | 10.09441 | 0.17276 | 1.188581 | 3.30E-05 | 0.00065 |
| 413 | RP11-693J15.5 | 0.798823 | 2.788458 | 0.172638 | 1.188436 | 3.34E-05 | 0.000656 |
| 414 | NT5E | 0.140953 | 10.8415 | 0.170534 | 1.185938 | 4.12E-05 | 0.000807 |
| 415 | EVI2A | 1.459279 | 2.000117 | 0.170032 | 1.185343 | 4.33E-05 | 0.000846 |
| 416 | YPEL2 | 0.069847 | 20.76582 | 0.168725 | 1.183794 | 4.92E-05 | 0.000959 |
| 417 | MANF | 9.270245 | 1.201482 | 0.167846 | 1.182755 | 5.35E-05 | 0.001042 |

| | | | | | | | |
|---|---|---|---|---|---|---|---|
| 418 | IGKV4-1 | 0.134145 | 11.28164 | 0.165541 | 1.180031 | 6.69E-05 | 0.001298 |
| 419 | HSP90B1 | 10.60661 | 1.179469 | 0.165038 | 1.179438 | 7.02E-05 | 0.001359 |
| 420 | ALDOC | 0.21594 | 7.396479 | 0.164217 | 1.17847 | 7.59E-05 | 0.001466 |
| 421 | MZB1 | 20.20867 | 1.115055 | 0.162384 | 1.176312 | 9.03E-05 | 0.001739 |
| 422 | DPEP2 | 0.074694 | 19.36265 | 0.16234 | 1.17626 | 9.06E-05 | 0.001742 |
| 423 | C12orf75 | 4.157763 | 1.376232 | 0.162258 | 1.176164 | 9.13E-05 | 0.001752 |
| 424 | BCL7A | 0.112674 | 13.18525 | 0.162064 | 1.175935 | 9.30E-05 | 0.00178 |
| 425 | PSTPIP2 | 0.049371 | 28.7344 | 0.161552 | 1.175334 | 9.76E-05 | 0.001863 |
| 426 | IL6R | 0.084365 | 17.23815 | 0.160888 | 1.174553 | 0.000104 | 0.001977 |
| 427 | EIF4EBP1 | 0.757814 | 2.847972 | 0.160729 | 1.174366 | 0.000105 | 0.002002 |
| 428 | PFN2 | 0.093833 | 15.59545 | 0.160328 | 1.173895 | 0.000109 | 0.002074 |
| 429 | C4orf32 | 0.121531 | 12.27862 | 0.160266 | 1.173823 | 0.00011 | 0.002081 |
| 430 | RP3-325F22.5 | 0.149707 | 10.15513 | 0.159357 | 1.172757 | 0.00012 | 0.002258 |
| 431 | SESN2 | 0.188701 | 8.263364 | 0.158263 | 1.171474 | 0.000132 | 0.002491 |
| 432 | KCNK1 | 0.084263 | 17.21163 | 0.158217 | 1.171421 | 0.000133 | 0.002496 |
| 433 | BIK | 0.971004 | 2.444782 | 0.157677 | 1.170788 | 0.00014 | 0.002617 |
| 434 | DNAJC5B | 0.093662 | 15.56696 | 0.156793 | 1.169754 | 0.000151 | 0.00283 |
| 435 | ADAMTS12 | 0.207518 | 7.592748 | 0.156171 | 1.169027 | 0.00016 | 0.002987 |
| 436 | RP11-326C3.12 | 0.174496 | 8.830517 | 0.155932 | 1.168747 | 0.000164 | 0.003045 |
| 437 | TLR1 | 0.126137 | 11.81517 | 0.15577 | 1.168557 | 0.000166 | 0.003078 |
| 438 | LILRB4 | 0.218237 | 7.267996 | 0.155764 | 1.16855 | 0.000166 | 0.003078 |
| 439 | INSR | 0.296254 | 5.624794 | 0.155194 | 1.167885 | 0.000175 | 0.003233 |
| 440 | CD3E | 0.121496 | 12.21901 | 0.15514 | 1.167822 | 0.000176 | 0.003241 |
| 441 | CD82 | 0.641958 | 3.153805 | 0.154583 | 1.167171 | 0.000185 | 0.0034 |
| 442 | ANG | 0.110251 | 13.34904 | 0.154374 | 1.166928 | 0.000188 | 0.003456 |
| 443 | CENPV | 0.377819 | 4.631045 | 0.154347 | 1.166895 | 0.000189 | 0.003457 |
| 444 | IFNG-AS1 | 0.136641 | 10.96859 | 0.154105 | 1.166614 | 0.000193 | 0.003524 |
| 445 | MYO1F | 0.540306 | 3.547935 | 0.15354 | 1.165954 | 0.000203 | 0.003699 |
| 446 | IGHG4 | 0.137777 | 10.87999 | 0.153487 | 1.165892 | 0.000204 | 0.003708 |
| 447 | AL109761.5 | 0.110389 | 13.31301 | 0.152823 | 1.165118 | 0.000216 | 0.003924 |

| | | | | | | | |
|---|---|---|---|---|---|---|---|
| 448 | LINC00886 | 0.049703 | 28.26315 | 0.151477 | 1.163551 | 0.000243 | 0.00441 |
| 449 | GSTM3 | 0.040412 | 34.51904 | 0.15137 | 1.163427 | 0.000246 | 0.004441 |
| 450 | KLHL6 | 0.088776 | 16.25661 | 0.150061 | 1.161905 | 0.000276 | 0.00497 |
| 451 | XBP1 | 4.744628 | 1.318401 | 0.149345 | 1.161073 | 0.000293 | 0.005278 |
| 452 | LINC01307 | 0.137883 | 10.82238 | 0.148877 | 1.16053 | 0.000305 | 0.005484 |
| 453 | RGL4 | 0.038033 | 36.51086 | 0.148564 | 1.160167 | 0.000314 | 0.005622 |
| 454 | MOXD1 | 0.044626 | 31.23002 | 0.147297 | 1.158698 | 0.00035 | 0.006255 |
| 455 | PHACTR1 | 1.043839 | 2.319355 | 0.145857 | 1.157031 | 0.000396 | 0.007057 |
| 456 | AUTS2 | 0.089533 | 16.05881 | 0.145767 | 1.156926 | 0.000399 | 0.007096 |
| 457 | GLUL | 0.148157 | 10.10625 | 0.14519 | 1.156259 | 0.000419 | 0.007434 |
| 458 | KDM2B | 0.215781 | 7.255553 | 0.144345 | 1.155282 | 0.00045 | 0.007966 |
| 459 | GCSAM | 0.785029 | 2.739985 | 0.144258 | 1.155182 | 0.000453 | 0.008007 |
| 460 | FTH1 | 59.37972 | 1.050792 | 0.143899 | 1.154768 | 0.000467 | 0.008234 |
| 461 | SORL1 | 0.400944 | 4.376459 | 0.143571 | 1.154389 | 0.00048 | 0.008445 |
| 462 | HIP1 | 0.050498 | 27.60886 | 0.143337 | 1.154119 | 0.000489 | 0.008592 |
| 463 | MIB2 | 0.215455 | 7.255598 | 0.143056 | 1.153794 | 0.000501 | 0.008778 |
| 464 | SMYD3 | 0.151255 | 9.897882 | 0.142925 | 1.153644 | 0.000506 | 0.008854 |
| 465 | GPR155 | 0.212498 | 7.334812 | 0.14204 | 1.152623 | 0.000545 | 0.009509 |

**Supplementary Table S2. HVGs of lung airway epithelial cells (FDR<0.01).**

| id | genes | u | cv2 | residualcv2 | fitratio | pval | fdr |
|---|---|---|---|---|---|---|---|
| 1 | IGFBP5 | 0.3422 | 366.8569 | 4.521678 | 91.98979 | 0 | 0 |
| 2 | COL1A2 | 0.402753 | 162.9321 | 3.852893 | 47.12919 | 0 | 0 |
| 3 | POSTN | 0.15356 | 253.3906 | 3.415588 | 30.43484 | 0 | 0 |
| 4 | COL1A1 | 0.68134 | 62.38933 | 3.331199 | 27.97186 | 0 | 0 |
| 5 | SPARC | 0.30297 | 106.0357 | 3.171943 | 23.85378 | 0 | 0 |
| 6 | TSLP | 0.658173 | 43.62917 | 2.945928 | 19.02831 | 0 | 0 |
| 7 | IGFL1 | 0.334309 | 75.65533 | 2.92221 | 18.5823 | 0 | 0 |
| 8 | IFNL2 | 0.520793 | 50.31697 | 2.896921 | 18.11827 | 0 | 0 |
| 9 | IFNL3 | 0.442755 | 52.73862 | 2.806557 | 16.55282 | 0 | 0 |
| 10 | CCL2 | 1.451734 | 15.0024 | 2.454099 | 11.63595 | 0 | 0 |
| 11 | IL23A | 1.439021 | 14.56089 | 2.418541 | 11.22946 | 0 | 0 |
| 12 | SERPINB4 | 1.837869 | 12.3673 | 2.406701 | 11.09729 | 0 | 0 |
| 13 | S100A7 | 0.457983 | 29.3218 | 2.248435 | 9.472897 | 0 | 0 |
| 14 | COL3A1 | 0.10598 | 111.0061 | 2.236555 | 9.361029 | 0 | 0 |
| 15 | COL6A3 | 0.044814 | 234.9718 | 2.148192 | 8.569349 | 0 | 0 |
| 16 | PI3 | 3.386293 | 6.972353 | 2.147977 | 8.56751 | 0 | 0 |
| 17 | RP11-338I21.1 | 0.035711 | 286.7962 | 2.123833 | 8.363136 | 0 | 0 |
| 18 | TEX26-AS1 | 0.030836 | 330.5665 | 2.12091 | 8.338725 | 0 | 0 |
| 19 | OVOS2 | 0.1025 | 101.2763 | 2.1127 | 8.270542 | 0 | 0 |
| 20 | GREM1 | 0.038824 | 256.9982 | 2.096538 | 8.13795 | 0 | 0 |
| 21 | AMTN | 0.481061 | 20.84226 | 1.948835 | 7.020506 | 0 | 0 |
| 22 | TNFAIP6 | 1.47615 | 8.760052 | 1.926826 | 6.867676 | 0 | 0 |
| 23 | S100A8 | 1.074926 | 10.35836 | 1.879684 | 6.551435 | 0 | 0 |
| 24 | MSMB | 1.007618 | 10.74332 | 1.869777 | 6.486847 | 0 | 0 |
| 25 | IL17C | 2.239176 | 6.274484 | 1.839883 | 6.295805 | 0 | 0 |
| 26 | LY6D | 0.186798 | 43.30096 | 1.832953 | 6.252323 | 0 | 0 |
| 27 | SLC15A2 | 0.295966 | 28.34831 | 1.831725 | 6.244652 | 0 | 0 |

| | | | | | | | |
|---|---|---|---|---|---|---|---|
| 28 | KRT81 | 0.094953 | 80.34071 | 1.807393 | 6.09454 | 0 | 0 |
| 29 | AREG | 9.075432 | 3.566721 | 1.799062 | 6.043975 | 0 | 0 |
| 30 | CXCL6 | 0.678661 | 13.03051 | 1.761965 | 5.823868 | 0 | 0 |
| 31 | CTGF | 0.146548 | 50.33998 | 1.755226 | 5.784755 | 0 | 0 |
| 32 | CRCT1 | 0.47111 | 17.12071 | 1.734429 | 5.665694 | 0 | 0 |
| 33 | PPBP | 0.758688 | 11.41749 | 1.717488 | 5.570517 | 0 | 0 |
| 34 | MMP1 | 0.698346 | 11.95643 | 1.698626 | 5.466434 | 0 | 0 |
| 35 | COL6A1 | 0.24869 | 29.02535 | 1.6975 | 5.460278 | 0 | 0 |
| 36 | SERPINB2 | 2.356929 | 5.216303 | 1.682605 | 5.379553 | 0 | 0 |
| 37 | SUGCT | 2.210794 | 5.232406 | 1.651336 | 5.213943 | 0 | 0 |
| 38 | S100A9 | 24.53591 | 2.468123 | 1.584215 | 4.875464 | 0 | 0 |
| 39 | MEG3 | 0.030171 | 190.7004 | 1.549247 | 4.707925 | 0 | 0 |
| 40 | IFNL1 | 0.906759 | 8.333716 | 1.538342 | 4.656862 | 0 | 0 |
| 41 | CXCL5 | 5.059604 | 3.155575 | 1.511859 | 4.535152 | 0 | 0 |
| 42 | FBXO32 | 0.837573 | 8.458314 | 1.49349 | 4.452609 | 0 | 0 |
| 43 | KRT6B | 2.948961 | 3.709274 | 1.453863 | 4.279615 | 0 | 0 |
| 44 | KRT14 | 0.823119 | 8.221108 | 1.451798 | 4.270787 | 0 | 0 |
| 45 | CCL5 | 15.97882 | 2.228143 | 1.431141 | 4.183469 | 0 | 0 |
| 46 | HAS2 | 0.0571 | 90.29242 | 1.429499 | 4.176607 | 0 | 0 |
| 47 | ANKRD1 | 0.126662 | 41.07686 | 1.413192 | 4.10905 | 0 | 0 |
| 48 | SULF1 | 0.046107 | 108.738 | 1.405627 | 4.078084 | 0 | 0 |
| 49 | SERPINB3 | 0.841328 | 7.692464 | 1.401977 | 4.063225 | 0 | 0 |
| 50 | CCL7 | 0.024243 | 202.5249 | 1.39286 | 4.026348 | 0 | 0 |
| 51 | NEXN | 0.469267 | 12.0642 | 1.381055 | 3.979096 | 0 | 0 |
| 52 | LCE3D | 0.054741 | 86.82733 | 1.34906 | 3.853802 | 0 | 0 |
| 53 | IL20 | 0.066254 | 71.97708 | 1.34811 | 3.850142 | 0 | 0 |
| 54 | THBS2 | 0.073408 | 64.69675 | 1.341375 | 3.8243 | 0 | 0 |
| 55 | SBSN | 0.03048 | 153.0234 | 1.339206 | 3.816012 | 2.96e-323 | 5.90-321 |
| 56 | GDF15 | 2.710583 | 3.426464 | 1.333822 | 3.795521 | 5.22e-320 | 1.02e-317 |
| 57 | COL6A2 | 0.031812 | 144.0976 | 1.321402 | 3.748673 | 1.56e-312 | 3.00-310 |

| | | | | | | | |
|---|---|---|---|---|---|---|---|
| 58 | IL13RA2 | 0.543652 | 9.653596 | 1.281669 | 3.602647 | 1.89E-289 | 3.57E-287 |
| 59 | LUM | 0.016842 | 259.2768 | 1.278459 | 3.5911 | 1.22E-287 | 2.27E-285 |
| 60 | VIM | 0.640769 | 8.331497 | 1.268731 | 3.556335 | 3.34E-282 | 6.10E-280 |
| 61 | CGA | 0.044896 | 97.29511 | 1.26828 | 3.554734 | 5.93E-282 | 1.07E-279 |
| 62 | DKK1 | 0.303146 | 15.37342 | 1.241329 | 3.460209 | 2.81E-267 | 4.92E-265 |
| 63 | ADAMTS2 | 0.015892 | 264.6734 | 1.241323 | 3.46019 | 2.83E-267 | 4.92E-265 |
| 64 | CCL3L3 | 0.036405 | 116.2132 | 1.239485 | 3.453833 | 2.71E-266 | 4.64E-264 |
| 65 | IFNB1 | 0.075163 | 56.06027 | 1.221074 | 3.390829 | 1.31E-256 | 2.21E-254 |
| 66 | CSF2 | 3.525373 | 2.669806 | 1.205479 | 3.338359 | 1.34E-248 | 2.22E-246 |
| 67 | SFTA1P | 1.283625 | 4.638519 | 1.199123 | 3.317207 | 2.19E-245 | 3.58E-243 |
| 68 | TIMP1 | 9.432114 | 1.940592 | 1.198977 | 3.316721 | 2.59E-245 | 4.18E-243 |
| 69 | OLR1 | 0.427302 | 10.84251 | 1.194176 | 3.300838 | 6.62E-243 | 1.05E-240 |
| 70 | RGS2 | 0.769233 | 6.588071 | 1.17831 | 3.24888 | 4.57E-235 | 7.16E-233 |
| 71 | ITGA1 | 0.17072 | 24.20949 | 1.167215 | 3.213031 | 1.09E-229 | 1.68E-227 |
| 72 | FST | 7.060735 | 1.9896 | 1.15296 | 3.167554 | 6.59E-223 | 1.00E-220 |
| 73 | DHRS9 | 0.730404 | 6.605607 | 1.140614 | 3.128688 | 3.80E-217 | 5.71E-215 |
| 74 | CYP24A1 | 0.510087 | 8.833089 | 1.1397 | 3.12583 | 1.00E-216 | 1.49E-214 |
| 75 | TAGLN | 4.963836 | 2.143319 | 1.118444 | 3.060088 | 4.64E-207 | 6.78E-205 |
| 76 | PTX3 | 0.295949 | 13.82639 | 1.113685 | 3.045559 | 6.14E-205 | 8.86E-203 |
| 77 | MMP13 | 0.205037 | 19.29789 | 1.111514 | 3.038956 | 5.64E-204 | 8.03E-202 |
| 78 | AQP3 | 4.433787 | 2.187756 | 1.098256 | 2.998931 | 3.69E-198 | 5.18E-196 |
| 79 | CLNK | 0.291392 | 13.62221 | 1.08484 | 2.958968 | 2.16E-192 | 3.00E-190 |
| 80 | IL6 | 6.026135 | 1.911218 | 1.067052 | 2.906797 | 6.38E-185 | 8.74E-183 |
| 81 | LTB | 11.41982 | 1.609612 | 1.050832 | 2.860031 | 2.77E-178 | 3.74E-176 |
| 82 | SRGN | 0.380583 | 10.37734 | 1.049867 | 2.857271 | 6.79E-178 | 9.07E-176 |
| 83 | SPP1 | 0.068844 | 50.73216 | 1.03572 | 2.817133 | 3.02E-172 | 3.98E-170 |
| 84 | NUPR1 | 16.61788 | 1.452156 | 1.008497 | 2.741476 | 1.01E-161 | 1.32E-159 |
| 85 | CYP1B1 | 0.107756 | 31.41651 | 0.990274 | 2.691971 | 6.35E-155 | 8.20E-153 |
| 86 | IL36G | 9.119815 | 1.584866 | 0.989014 | 2.688581 | 1.84E-154 | 2.35E-152 |
| 87 | MMP9 | 0.212473 | 16.50988 | 0.988497 | 2.687194 | 2.85E-154 | 3.59E-152 |

| | | | | | | | |
|---|---|---|---|---|---|---|---|
| 88 | IL1RL1 | 0.235331 | 14.97132 | 0.984882 | 2.677495 | 5.99E-153 | 7.46E-151 |
| 89 | COL4A1 | 0.422932 | 8.838681 | 0.980983 | 2.667077 | 1.56E-151 | 1.93E-149 |
| 90 | DUOXA2 | 3.124207 | 2.233415 | 0.973434 | 2.647019 | 8.19E-149 | 9.97E-147 |
| 91 | CXCL9 | 0.067621 | 48.11255 | 0.965227 | 2.625383 | 6.79E-146 | 8.18E-144 |
| 92 | KRT23 | 0.507769 | 7.444874 | 0.964902 | 2.62453 | 8.85E-146 | 1.05E-143 |
| 93 | PLAT | 1.240323 | 3.724778 | 0.956505 | 2.602585 | 7.79E-143 | 9.18E-141 |
| 94 | RRAD | 0.062065 | 51.77931 | 0.954983 | 2.598627 | 2.64E-142 | 3.08E-140 |
| 95 | SPRR2D | 0.108205 | 29.41605 | 0.928475 | 2.530648 | 2.73E-133 | 3.15E-131 |
| 96 | LINC00958 | 2.04908 | 2.648251 | 0.92828 | 2.530155 | 3.17E-133 | 3.62E-131 |
| 97 | NKX3-1 | 1.471711 | 3.226156 | 0.925982 | 2.524346 | 1.84E-132 | 2.08E-130 |
| 98 | INHBA | 6.242686 | 1.619937 | 0.912333 | 2.490125 | 5.46E-128 | 6.10E-126 |
| 99 | CYR61 | 1.479118 | 3.165918 | 0.910356 | 2.485206 | 2.38E-127 | 2.63E-125 |
| 100 | CCNA1 | 1.037619 | 3.977149 | 0.897215 | 2.452762 | 3.74E-123 | 4.11E-121 |
| 101 | CKS2 | 6.927309 | 1.547587 | 0.896489 | 2.450983 | 6.35E-123 | 6.89E-121 |
| 102 | STC1 | 1.893739 | 2.680656 | 0.895264 | 2.447983 | 1.54E-122 | 1.66E-120 |
| 103 | BAMBI | 0.125974 | 24.58496 | 0.894682 | 2.446557 | 2.35E-122 | 2.51E-120 |
| 104 | CLEC2D | 0.634284 | 5.727787 | 0.885821 | 2.424974 | 1.37E-119 | 1.45E-117 |
| 105 | CX3CL1 | 1.06441 | 3.850671 | 0.883138 | 2.418477 | 9.28E-119 | 9.69E-117 |
| 106 | IL19 | 0.051476 | 56.77675 | 0.863981 | 2.372587 | 6.04E-113 | 6.24E-111 |
| 107 | SPRR1B | 0.552169 | 6.270775 | 0.863115 | 2.370534 | 1.09E-112 | 1.12E-110 |
| 108 | NREP | 0.178627 | 17.03359 | 0.85813 | 2.358746 | 3.30E-111 | 3.35E-109 |
| 109 | CLEC2B | 0.493746 | 6.846158 | 0.857518 | 2.357302 | 5.01E-111 | 5.04E-109 |
| 110 | KRTAP2-3 | 0.110306 | 26.6673 | 0.848835 | 2.336924 | 1.75E-108 | 1.74E-106 |
| 111 | CTD-2017D11.1 | 0.16016 | 18.31027 | 0.827838 | 2.288367 | 1.73E-102 | 1.71E-100 |
| 112 | CH25H | 0.063969 | 44.2392 | 0.827112 | 2.286706 | 2.76E-102 | 2.70E-100 |
| 113 | TNF | 0.364563 | 8.612859 | 0.825805 | 2.283719 | 6.41E-102 | 6.22E-100 |
| 114 | S100A4 | 0.937216 | 3.982929 | 0.824546 | 2.280846 | 1.44E-101 | 1.38E-99 |
| 115 | PTHLH | 0.720085 | 4.866944 | 0.823995 | 2.279589 | 2.05E-101 | 1.95E-99 |
| 116 | AKAP12 | 1.428491 | 2.966611 | 0.82286 | 2.277003 | 4.24E-101 | 4.00E-99 |
| 117 | ALDH3A1 | 0.461551 | 6.911097 | 0.809842 | 2.247553 | 1.59E-97 | 1.49E-95 |

| | | | | | | | |
|---|---|---|---|---|---|---|---|
| 118 | CCL20 | 46.66523 | 1.083928 | 0.808579 | 2.244715 | 3.50E-97 | 3.25E-95 |
| 119 | LY6K | 0.453381 | 7.003775 | 0.807938 | 2.243277 | 5.22E-97 | 4.81E-95 |
| 120 | FN1 | 11.32735 | 1.24032 | 0.788674 | 2.200478 | 6.89E-92 | 6.30E-90 |
| 121 | CCDC71L | 2.827852 | 1.944631 | 0.788055 | 2.199115 | 1.00E-91 | 9.06E-90 |
| 122 | FBN1 | 0.029648 | 90.62215 | 0.787948 | 2.198881 | 1.07E-91 | 9.58E-90 |
| 123 | AKR1C2 | 0.315185 | 9.380832 | 0.782226 | 2.186333 | 3.26E-90 | 2.90E-88 |
| 124 | KRT80 | 0.092043 | 29.52914 | 0.776431 | 2.173701 | 1.00E-88 | 8.86E-87 |
| 125 | APOE | 1.117765 | 3.340811 | 0.77573 | 2.172178 | 1.51E-88 | 1.33E-86 |
| 126 | CXCL3 | 17.61087 | 1.137975 | 0.772467 | 2.165101 | 1.02E-87 | 8.90E-86 |
| 127 | SDC2 | 0.055933 | 47.57607 | 0.768567 | 2.156673 | 9.89E-87 | 8.53E-85 |
| 128 | CLIC3 | 0.029454 | 88.71605 | 0.760192 | 2.138687 | 1.22E-84 | 1.05E-82 |
| 129 | EREG | 1.559601 | 2.614265 | 0.752548 | 2.122402 | 9.28E-83 | 7.89E-81 |
| 130 | IL33 | 7.242763 | 1.309399 | 0.741465 | 2.099008 | 4.44E-80 | 3.74E-78 |
| 131 | C8orf4 | 0.25284 | 10.90537 | 0.733702 | 2.082776 | 3.10E-78 | 2.59E-76 |
| 132 | FLJ22447 | 0.059563 | 43.1313 | 0.732023 | 2.079284 | 7.69E-78 | 6.38E-76 |
| 133 | KIAA0101 | 0.498177 | 5.96862 | 0.727869 | 2.070663 | 7.21E-77 | 5.94E-75 |
| 134 | SLC2A3 | 0.041903 | 60.03886 | 0.717636 | 2.049583 | 1.66E-74 | 1.36E-72 |
| 135 | PTGS2 | 1.72444 | 2.358165 | 0.711458 | 2.036959 | 4.20E-73 | 3.41E-71 |
| 136 | ANGPTL4 | 1.884057 | 2.196767 | 0.6932 | 2.000106 | 4.71E-69 | 3.79E-67 |
| 137 | CLDN11 | 0.192502 | 13.31843 | 0.681992 | 1.977813 | 1.22E-66 | 9.78E-65 |
| 138 | EDN1 | 3.563444 | 1.574349 | 0.681905 | 1.977641 | 1.27E-66 | 1.01E-64 |
| 139 | EGR1 | 1.149755 | 2.981462 | 0.68168 | 1.977197 | 1.42E-66 | 1.12E-64 |
| 140 | CDH11 | 0.062147 | 39.06004 | 0.674382 | 1.962819 | 4.96E-65 | 3.88E-63 |
| 141 | ADAMTS1 | 0.855461 | 3.668829 | 0.67422 | 1.962502 | 5.36E-65 | 4.17E-63 |
| 142 | TESC | 0.072753 | 33.2997 | 0.668483 | 1.951275 | 8.42E-64 | 6.50E-62 |
| 143 | NR4A3 | 0.12814 | 19.27864 | 0.667812 | 1.949966 | 1.16E-63 | 8.89E-62 |
| 144 | RP11-357H14.17 | 0.032716 | 72.71821 | 0.665186 | 1.944852 | 4.04E-63 | 3.07E-61 |
| 145 | CCK | 0.317478 | 8.259602 | 0.661407 | 1.937517 | 2.41E-62 | 1.82E-60 |
| 146 | RBP1 | 1.219065 | 2.785477 | 0.654117 | 1.923443 | 7.23E-61 | 5.43E-59 |
| 147 | SERPINE1 | 0.54234 | 5.16081 | 0.653428 | 1.922119 | 9.95E-61 | 7.42E-59 |

| | | | | | | | |
|---|---|---|---|---|---|---|---|
| 148 | FOLR1 | 0.437277 | 6.150304 | 0.647036 | 1.909872 | 1.88E-59 | 1.39E-57 |
| 149 | SLC6A14 | 3.244538 | 1.577538 | 0.642914 | 1.902016 | 1.22E-58 | 9.00E-57 |
| 150 | TNIP3 | 0.877111 | 3.467474 | 0.6366 | 1.890044 | 2.09E-57 | 1.53E-55 |
| 151 | CTSC | 5.947212 | 1.245266 | 0.634621 | 1.886307 | 5.05E-57 | 3.67E-55 |
| 152 | TGM2 | 6.803 | 1.196253 | 0.633951 | 1.885043 | 6.81E-57 | 4.91E-55 |
| 153 | FGFBP1 | 0.674075 | 4.239445 | 0.633704 | 1.884578 | 7.60E-57 | 5.44E-55 |
| 154 | COL5A1 | 0.086037 | 27.26816 | 0.631485 | 1.880401 | 2.03E-56 | 1.44E-54 |
| 155 | MAP1B | 0.064541 | 36.02835 | 0.630511 | 1.878571 | 3.12E-56 | 2.21E-54 |
| 156 | CTHRC1 | 0.049044 | 46.89721 | 0.625302 | 1.868811 | 3.06E-55 | 2.15E-53 |
| 157 | SERPINB7 | 2.149317 | 1.901199 | 0.623492 | 1.865431 | 6.74E-55 | 4.70E-53 |
| 158 | SLC34A2 | 0.422726 | 6.168513 | 0.620883 | 1.86057 | 2.09E-54 | 1.45E-52 |
| 159 | MMP7 | 0.362101 | 7.057947 | 0.620746 | 1.860316 | 2.21E-54 | 1.53E-52 |
| 160 | BPGM | 3.264412 | 1.526346 | 0.612663 | 1.845339 | 7.05E-53 | 4.83E-51 |
| 161 | CD55 | 1.280787 | 2.581217 | 0.611498 | 1.843191 | 1.15E-52 | 7.86E-51 |
| 162 | MT1E | 16.43354 | 0.971858 | 0.605363 | 1.831917 | 1.52E-51 | 1.03E-49 |
| 163 | C1orf56 | 2.449275 | 1.737199 | 0.603226 | 1.828007 | 3.71E-51 | 2.50E-49 |
| 164 | MYL9 | 2.717297 | 1.647673 | 0.602878 | 1.82737 | 4.29E-51 | 2.87E-49 |
| 165 | AC004791.2 | 0.051829 | 43.32719 | 0.600334 | 1.822728 | 1.23E-50 | 8.18E-49 |
| 166 | THBS1 | 1.939852 | 1.965333 | 0.598813 | 1.819956 | 2.31E-50 | 1.52E-48 |
| 167 | KRBOX1 | 0.038551 | 57.80122 | 0.597541 | 1.817643 | 3.89E-50 | 2.55E-48 |
| 168 | MROH8 | 0.073895 | 30.49126 | 0.595539 | 1.814009 | 8.83E-50 | 5.76E-48 |
| 169 | ESM1 | 0.039579 | 55.92396 | 0.590442 | 1.804786 | 7.01E-49 | 4.55E-47 |
| 170 | TNFSF15 | 0.200596 | 11.69562 | 0.590394 | 1.8047 | 7.15E-49 | 4.61E-47 |
| 171 | S100P | 2.163384 | 1.829495 | 0.58864 | 1.801537 | 1.45E-48 | 9.30E-47 |
| 172 | RHCG | 0.67339 | 4.0092 | 0.577053 | 1.780783 | 1.45E-46 | 9.24E-45 |
| 173 | TNFAIP2 | 9.340519 | 1.041381 | 0.574387 | 1.776041 | 4.11E-46 | 2.61E-44 |
| 174 | WNT5A | 0.051098 | 42.71092 | 0.572079 | 1.771947 | 1.01E-45 | 6.36E-44 |
| 175 | AIM2 | 0.853244 | 3.310872 | 0.569598 | 1.767557 | 2.63E-45 | 1.65E-43 |
| 176 | CKB | 0.787468 | 3.50894 | 0.566464 | 1.762026 | 8.79E-45 | 5.47E-43 |
| 177 | TIMP3 | 2.067739 | 1.834166 | 0.566068 | 1.761327 | 1.02E-44 | 6.33E-43 |

| | | | | | | | |
|---|---|---|---|---|---|---|---|
| 178 | LAYN | 0.783748 | 3.511114 | 0.563433 | 1.756694 | 2.79E-44 | 1.72E-42 |
| 179 | ARL4C | 1.175102 | 2.604544 | 0.561672 | 1.753602 | 5.45E-44 | 3.34E-42 |
| 180 | NR4A2 | 0.332347 | 7.151168 | 0.558069 | 1.747295 | 2.12E-43 | 1.29E-41 |
| 181 | TUBA1A | 0.788711 | 3.455106 | 0.552218 | 1.737102 | 1.88E-42 | 1.14E-40 |
| 182 | NR2F2 | 2.114002 | 1.781414 | 0.549242 | 1.73194 | 5.65E-42 | 3.40E-40 |
| 183 | ADAMTS9 | 0.263164 | 8.721966 | 0.546755 | 1.727638 | 1.41E-41 | 8.42E-40 |
| 184 | AQP5 | 0.063895 | 33.43544 | 0.545999 | 1.726331 | 1.85E-41 | 1.10E-39 |
| 185 | FAM43A | 0.295108 | 7.855156 | 0.545716 | 1.725843 | 2.06E-41 | 1.22E-39 |
| 186 | SLC7A11 | 0.757805 | 3.53998 | 0.545557 | 1.725569 | 2.18E-41 | 1.28E-39 |
| 187 | ATP2B1 | 1.462482 | 2.212451 | 0.544745 | 1.724169 | 2.93E-41 | 1.72E-39 |
| 188 | GAL | 2.522055 | 1.612636 | 0.543916 | 1.72274 | 3.96E-41 | 2.31E-39 |
| 189 | LGALSL | 0.782529 | 3.421232 | 0.5363 | 1.70967 | 6.15E-40 | 3.56E-38 |
| 190 | EPB41L3 | 0.019131 | 108.645 | 0.535203 | 1.707796 | 9.08E-40 | 5.24E-38 |
| 191 | KCNJ2 | 0.058154 | 35.67521 | 0.518815 | 1.680035 | 2.77E-37 | 1.59E-35 |
| 192 | COL5A2 | 0.231167 | 9.519108 | 0.515639 | 1.674709 | 8.17E-37 | 4.67E-35 |
| 193 | NR2F1 | 0.037995 | 54.00307 | 0.515252 | 1.674059 | 9.32E-37 | 5.29E-35 |
| 194 | UCHL1 | 0.972363 | 2.83803 | 0.512688 | 1.669773 | 2.22E-36 | 1.25E-34 |
| 195 | NPW | 0.225273 | 9.716505 | 0.512389 | 1.669274 | 2.45E-36 | 1.38E-34 |
| 196 | IGF2 | 0.063136 | 32.59572 | 0.508889 | 1.663442 | 7.93E-36 | 4.44E-34 |
| 197 | HIST1H2AC | 1.538272 | 2.060375 | 0.50577 | 1.658262 | 2.24E-35 | 1.25E-33 |
| 198 | DDIT3 | 0.988941 | 2.778773 | 0.503919 | 1.655196 | 4.13E-35 | 2.28E-33 |
| 199 | MT1X | 8.212007 | 0.998223 | 0.502192 | 1.652339 | 7.28E-35 | 4.01E-33 |
| 200 | VCAN | 0.29549 | 7.493307 | 0.499719 | 1.648258 | 1.64E-34 | 8.97E-33 |
| 201 | ZNF778 | 0.139839 | 14.97458 | 0.498324 | 1.645961 | 2.58E-34 | 1.41E-32 |
| 202 | LOX | 0.070606 | 28.91498 | 0.498134 | 1.645648 | 2.74E-34 | 1.49E-32 |
| 203 | LIF | 2.178442 | 1.658488 | 0.494316 | 1.639376 | 9.43E-34 | 5.09E-32 |
| 204 | NUMBL | 0.138755 | 14.98382 | 0.491547 | 1.634843 | 2.29E-33 | 1.23E-31 |
| 205 | ELF3 | 2.404905 | 1.564647 | 0.48908 | 1.630816 | 5.03E-33 | 2.69E-31 |
| 206 | SERPINB1 | 10.10383 | 0.939698 | 0.48845 | 1.629789 | 6.14E-33 | 3.27E-31 |
| 207 | ID2 | 0.272889 | 7.950052 | 0.487036 | 1.627485 | 9.61E-33 | 5.09E-31 |

| | | | | | | | |
|---|---|---|---|---|---|---|---|
| 208 | SCD | 0.230175 | 9.28196 | 0.486457 | 1.626544 | 1.15E-32 | 6.08E-31 |
| 209 | TFPI | 0.752566 | 3.33136 | 0.47942 | 1.615137 | 1.04E-31 | 5.46E-30 |
| 210 | GJB2 | 5.533565 | 1.086293 | 0.475298 | 1.608493 | 3.71E-31 | 1.94E-29 |
| 211 | TPM2 | 4.692049 | 1.149045 | 0.475088 | 1.608155 | 3.96E-31 | 2.06E-29 |
| 212 | CYGB | 0.343181 | 6.380953 | 0.472557 | 1.604091 | 8.57E-31 | 4.43E-29 |
| 213 | LAMA4 | 0.028931 | 67.51106 | 0.469353 | 1.598959 | 2.26E-30 | 1.17E-28 |
| 214 | GJB6 | 0.139544 | 14.5656 | 0.468626 | 1.597797 | 2.82E-30 | 1.44E-28 |
| 215 | EMP3 | 1.770688 | 1.813282 | 0.464647 | 1.591453 | 9.29E-30 | 4.73E-28 |
| 216 | RP11-297N6.4 | 0.049143 | 39.71304 | 0.461021 | 1.585692 | 2.72E-29 | 1.38E-27 |
| 217 | EFNA1 | 1.658897 | 1.870167 | 0.455969 | 1.577702 | 1.20E-28 | 6.05E-27 |
| 218 | GATA3 | 0.048338 | 39.90209 | 0.449554 | 1.567612 | 7.64E-28 | 3.84E-26 |
| 219 | IGFBP2 | 8.149336 | 0.948242 | 0.448954 | 1.566673 | 9.06E-28 | 4.54E-26 |
| 220 | SLPI | 3.186187 | 1.300741 | 0.441794 | 1.555495 | 6.88E-27 | 3.43E-25 |
| 221 | RARRES3 | 7.609456 | 0.951232 | 0.434871 | 1.544764 | 4.70E-26 | 2.33E-24 |
| 222 | AKR1C1 | 0.762154 | 3.150766 | 0.433526 | 1.542688 | 6.79E-26 | 3.35E-24 |
| 223 | NCCRP1 | 0.055646 | 34.19336 | 0.433228 | 1.542228 | 7.37E-26 | 3.62E-24 |
| 224 | IRG1 | 0.07257 | 26.3601 | 0.432336 | 1.540853 | 9.41E-26 | 4.60E-24 |
| 225 | SERPINB5 | 1.063493 | 2.453899 | 0.431955 | 1.540265 | 1.04E-25 | 5.09E-24 |
| 226 | SERPINA1 | 0.475836 | 4.614346 | 0.431777 | 1.539991 | 1.10E-25 | 5.32E-24 |
| 227 | NCF2 | 1.593752 | 1.866297 | 0.429091 | 1.535861 | 2.27E-25 | 1.10E-23 |
| 228 | BIRC3 | 6.481638 | 0.985544 | 0.426414 | 1.531755 | 4.67E-25 | 2.25E-23 |
| 229 | CNKSR3 | 0.178748 | 11.05336 | 0.426311 | 1.531598 | 4.80E-25 | 2.30E-23 |
| 230 | CLMP | 0.267812 | 7.605456 | 0.425685 | 1.530638 | 5.68E-25 | 2.71E-23 |
| 231 | POU2F2 | 2.582179 | 1.410221 | 0.421781 | 1.524675 | 1.61E-24 | 7.63E-23 |
| 232 | ATF3 | 0.253987 | 7.948156 | 0.421518 | 1.524274 | 1.72E-24 | 8.15E-23 |
| 233 | SYT8 | 0.173626 | 11.29342 | 0.42054 | 1.522784 | 2.23E-24 | 1.05E-22 |
| 234 | KRT6A | 11.94816 | 0.849444 | 0.420014 | 1.521983 | 2.56E-24 | 1.20E-22 |
| 235 | SPRR2A | 0.02985 | 62.28018 | 0.41962 | 1.521383 | 2.85E-24 | 1.33E-22 |
| 236 | G0S2 | 151.6594 | 0.706781 | 0.418782 | 1.520109 | 3.55E-24 | 1.65E-22 |
| 237 | ZFP36 | 1.36719 | 2.033134 | 0.416323 | 1.516376 | 6.75E-24 | 3.12E-22 |

| | | | | | | | |
|---|---|---|---|---|---|---|---|
| 238 | FILIP1L | 0.313922 | 6.51299 | 0.413768 | 1.512505 | 1.31E-23 | 6.04E-22 |
| 239 | CDKN2A | 2.438334 | 1.437392 | 0.411455 | 1.509012 | 2.38E-23 | 1.09E-21 |
| 240 | C1orf68 | 0.043623 | 42.41633 | 0.409769 | 1.50647 | 3.67E-23 | 1.68E-21 |
| 241 | CA2 | 0.388176 | 5.372408 | 0.408766 | 1.50496 | 4.74E-23 | 2.16E-21 |
| 242 | BEX1 | 0.021307 | 85.83769 | 0.406497 | 1.501548 | 8.45E-23 | 3.83E-21 |
| 243 | SERPINE2 | 0.46447 | 4.558695 | 0.399116 | 1.490506 | 5.37E-22 | 2.42E-20 |
| 244 | LRRC75A | 0.865917 | 2.760278 | 0.398844 | 1.490102 | 5.75E-22 | 2.58E-20 |
| 245 | PNRC1 | 5.125812 | 1.031089 | 0.397746 | 1.488466 | 7.54E-22 | 3.37E-20 |
| 246 | NDRG4 | 0.035654 | 51.12058 | 0.397658 | 1.488335 | 7.71E-22 | 3.44E-20 |
| 247 | S100A12 | 0.071562 | 25.74158 | 0.394986 | 1.484364 | 1.49E-21 | 6.60E-20 |
| 248 | MMP10 | 0.040301 | 45.17012 | 0.394701 | 1.48394 | 1.59E-21 | 7.05E-20 |
| 249 | LINC01436 | 0.118083 | 15.8465 | 0.393663 | 1.482402 | 2.05E-21 | 9.05E-20 |
| 250 | PELI1 | 1.324096 | 2.030156 | 0.393632 | 1.482355 | 2.07E-21 | 9.08E-20 |
| 251 | DUSP1 | 2.609113 | 1.358562 | 0.389698 | 1.476535 | 5.38E-21 | 2.35E-19 |
| 252 | KCNK5 | 0.075029 | 24.45176 | 0.389616 | 1.476414 | 5.48E-21 | 2.39E-19 |
| 253 | MT1M | 0.048469 | 37.44835 | 0.388729 | 1.475105 | 6.79E-21 | 2.94E-19 |
| 254 | NEDD9 | 0.297969 | 6.61253 | 0.382188 | 1.465487 | 3.22E-20 | 1.39E-18 |
| 255 | CTC-425F1.4 | 0.220765 | 8.682703 | 0.381254 | 1.46412 | 4.01E-20 | 1.72E-18 |
| 256 | IGFBP3 | 7.798741 | 0.894954 | 0.380164 | 1.462524 | 5.18E-20 | 2.22E-18 |
| 257 | NEURL3 | 1.28723 | 2.040229 | 0.379677 | 1.461813 | 5.80E-20 | 2.48E-18 |
| 258 | ASB2 | 1.115353 | 2.250322 | 0.379071 | 1.460927 | 6.69E-20 | 2.84E-18 |
| 259 | PRDM8 | 0.467133 | 4.43531 | 0.376537 | 1.45723 | 1.20E-19 | 5.10E-18 |
| 260 | SLC44A4 | 0.3689 | 5.429445 | 0.374771 | 1.454658 | 1.81E-19 | 7.64E-18 |
| 261 | HRASLS2 | 0.341377 | 5.798069 | 0.372097 | 1.450773 | 3.34E-19 | 1.40E-17 |
| 262 | ZFP36L2 | 1.156332 | 2.178909 | 0.372063 | 1.450724 | 3.37E-19 | 1.41E-17 |
| 263 | TNNT1 | 0.305202 | 6.392372 | 0.369855 | 1.447525 | 5.57E-19 | 2.32E-17 |
| 264 | BMP2 | 0.740948 | 3.01978 | 0.369092 | 1.44642 | 6.62E-19 | 2.75E-17 |
| 265 | SNAPC1 | 2.35789 | 1.397541 | 0.365746 | 1.441589 | 1.41E-18 | 5.80E-17 |
| 266 | TPM1 | 4.607126 | 1.036865 | 0.365736 | 1.441575 | 1.41E-18 | 5.80E-17 |
| 267 | ADIRF | 3.889909 | 1.106156 | 0.365357 | 1.441028 | 1.53E-18 | 6.29E-17 |

| | | | | | | | |
|---|---|---|---|---|---|---|---|
| 268 | MAP3K8 | 0.830245 | 2.738063 | 0.358911 | 1.431769 | 6.38E-18 | 2.61E-16 |
| 269 | CSF1 | 0.966038 | 2.443224 | 0.358121 | 1.430638 | 7.58E-18 | 3.09E-16 |
| 270 | CHST2 | 0.353214 | 5.544193 | 0.357455 | 1.429686 | 8.77E-18 | 3.56E-16 |
| 271 | MSC | 0.070821 | 24.98791 | 0.355123 | 1.426356 | 1.45E-17 | 5.88E-16 |
| 272 | MRGPRX3 | 0.28048 | 6.796186 | 0.355058 | 1.426263 | 1.47E-17 | 5.94E-16 |
| 273 | IL20RB | 0.263512 | 7.189783 | 0.354772 | 1.425855 | 1.57E-17 | 6.30E-16 |
| 274 | HMGB3 | 2.201927 | 1.43393 | 0.354694 | 1.425744 | 1.60E-17 | 6.38E-16 |
| 275 | CXCL1 | 33.92938 | 0.701017 | 0.352835 | 1.423097 | 2.38E-17 | 9.49E-16 |
| 276 | SERPINB13 | 0.059137 | 29.68804 | 0.351496 | 1.421192 | 3.17E-17 | 1.26E-15 |
| 277 | TAGLN3 | 0.434703 | 4.592979 | 0.349995 | 1.419061 | 4.37E-17 | 1.73E-15 |
| 278 | SNPH | 0.2158 | 8.58522 | 0.348953 | 1.417583 | 5.45E-17 | 2.15E-15 |
| 279 | CHAC1 | 0.147935 | 12.22514 | 0.348841 | 1.417424 | 5.58E-17 | 2.19E-15 |
| 280 | CITED2 | 0.94815 | 2.450936 | 0.347549 | 1.415594 | 7.33E-17 | 2.87E-15 |
| 281 | SLC39A8 | 2.129204 | 1.438423 | 0.339357 | 1.404045 | 4.03E-16 | 1.57E-14 |
| 282 | DSG3 | 0.191989 | 9.434106 | 0.334688 | 1.397504 | 1.04E-15 | 4.05E-14 |
| 283 | INSIG1 | 0.046571 | 36.8735 | 0.334031 | 1.396586 | 1.19E-15 | 4.61E-14 |
| 284 | TM4SF1 | 0.463672 | 4.271415 | 0.332562 | 1.394537 | 1.60E-15 | 6.17E-14 |
| 285 | PNMA2 | 0.108412 | 16.17942 | 0.332506 | 1.394458 | 1.62E-15 | 6.22E-14 |
| 286 | PLA2G4D | 0.071294 | 24.1807 | 0.32877 | 1.389258 | 3.40E-15 | 1.30E-13 |
| 287 | SNCG | 0.912039 | 2.46822 | 0.32585 | 1.385207 | 6.05E-15 | 2.31E-13 |
| 288 | KCCAT211 | 0.778525 | 2.776122 | 0.323394 | 1.38181 | 9.77E-15 | 3.72E-13 |
| 289 | CAMK2N1 | 2.388932 | 1.326909 | 0.320777 | 1.378198 | 1.62E-14 | 6.14E-13 |
| 290 | C11orf96 | 0.039416 | 42.87329 | 0.320648 | 1.37802 | 1.66E-14 | 6.28E-13 |
| 291 | GK | 0.127195 | 13.68867 | 0.318317 | 1.374812 | 2.59E-14 | 9.77E-13 |
| 292 | ID3 | 3.980683 | 1.045515 | 0.318246 | 1.374715 | 2.63E-14 | 9.87E-13 |
| 293 | PPP1R14A | 0.427204 | 4.510392 | 0.316889 | 1.37285 | 3.41E-14 | 1.27E-12 |
| 294 | ALPL | 0.083571 | 20.39646 | 0.312952 | 1.367456 | 7.15E-14 | 2.67E-12 |
| 295 | CD200 | 0.184276 | 9.583786 | 0.312147 | 1.366356 | 8.31E-14 | 3.09E-12 |
| 296 | TK1 | 0.1427 | 12.1853 | 0.311427 | 1.365372 | 9.51E-14 | 3.52E-12 |
| 297 | GS1-124K5.4 | 0.084753 | 20.07528 | 0.310693 | 1.36437 | 1.09E-13 | 4.02E-12 |

| | | | | | | | |
|---|---|---|---|---|---|---|---|
| 298 | PTGER4 | 0.340551 | 5.45887 | 0.309669 | 1.362974 | 1.32E-13 | 4.85E-12 |
| 299 | KRT16 | 0.113131 | 15.13806 | 0.306877 | 1.359174 | 2.20E-13 | 8.07E-12 |
| 300 | EGFLAM | 0.049386 | 33.84237 | 0.305886 | 1.357827 | 2.64E-13 | 9.64E-12 |
| 301 | ERRFI1 | 2.50938 | 1.272827 | 0.304705 | 1.356225 | 3.27E-13 | 1.19E-11 |
| 302 | ALMS1 | 0.113365 | 15.06028 | 0.303709 | 1.354874 | 3.92E-13 | 1.42E-11 |
| 303 | DUSP4 | 2.46182 | 1.283288 | 0.303027 | 1.353951 | 4.43E-13 | 1.60E-11 |
| 304 | TLR2 | 0.198269 | 8.818481 | 0.297181 | 1.346059 | 1.26E-12 | 4.54E-11 |
| 305 | TRAF1 | 0.15672 | 10.99093 | 0.296964 | 1.345767 | 1.31E-12 | 4.70E-11 |
| 306 | ICAM1 | 5.886176 | 0.891101 | 0.296788 | 1.34553 | 1.35E-12 | 4.83E-11 |
| 307 | NPPC | 0.037736 | 43.69501 | 0.296684 | 1.34539 | 1.37E-12 | 4.91E-11 |
| 308 | CRYAB | 0.051442 | 32.20685 | 0.296373 | 1.344972 | 1.45E-12 | 5.17E-11 |
| 309 | PVRL4 | 0.279204 | 6.425056 | 0.294778 | 1.342828 | 1.92E-12 | 6.81E-11 |
| 310 | CD70 | 0.038632 | 42.50302 | 0.292168 | 1.339328 | 3.03E-12 | 1.07E-10 |
| 311 | ADAMTS14 | 0.030981 | 52.83218 | 0.29186 | 1.338915 | 3.19E-12 | 1.13E-10 |
| 312 | GPX3 | 0.058662 | 28.1882 | 0.291769 | 1.338794 | 3.24E-12 | 1.14E-10 |
| 313 | ST3GAL4-AS1 | 0.05928 | 27.7986 | 0.288104 | 1.333896 | 6.09E-12 | 2.13E-10 |
| 314 | FOS | 0.241338 | 7.287793 | 0.288067 | 1.333846 | 6.13E-12 | 2.14E-10 |
| 315 | KLK6 | 0.118569 | 14.18307 | 0.286688 | 1.332009 | 7.75E-12 | 2.70E-10 |
| 316 | RRP12 | 0.483735 | 3.93246 | 0.285806 | 1.330835 | 8.99E-12 | 3.12E-10 |
| 317 | SAA2 | 5.195694 | 0.916728 | 0.284774 | 1.329461 | 1.07E-11 | 3.70E-10 |
| 318 | MAFB | 0.065628 | 24.942 | 0.279056 | 1.321881 | 2.78E-11 | 9.57E-10 |
| 319 | TRIB3 | 0.363744 | 4.994575 | 0.278923 | 1.321706 | 2.84E-11 | 9.75E-10 |
| 320 | HLA-DPA1 | 0.069501 | 23.55107 | 0.277581 | 1.319933 | 3.54E-11 | 1.21E-09 |
| 321 | HIST1H2BD | 0.858356 | 2.455664 | 0.275297 | 1.316922 | 5.13E-11 | 1.75E-09 |
| 322 | PRKCDBP | 0.330595 | 5.41177 | 0.274674 | 1.316102 | 5.68E-11 | 1.93E-09 |
| 323 | STC2 | 0.629737 | 3.120475 | 0.27267 | 1.313467 | 7.84E-11 | 2.66E-09 |
| 324 | BDNF | 0.326558 | 5.457673 | 0.272192 | 1.31284 | 8.46E-11 | 2.86E-09 |
| 325 | WNT5B | 0.033403 | 48.04385 | 0.271225 | 1.31157 | 9.88E-11 | 3.33E-09 |
| 326 | ANKRD37 | 0.380366 | 4.761479 | 0.270302 | 1.31036 | 1.14E-10 | 3.85E-09 |
| 327 | LGALS7B | 0.17042 | 9.885979 | 0.269934 | 1.309878 | 1.21E-10 | 4.07E-09 |

| | | | | | | | |
|---|---|---|---|---|---|---|---|
| 328 | ITGB8 | 0.502394 | 3.748529 | 0.269805 | 1.309709 | 1.24E-10 | 4.14E-09 |
| 329 | RARRES1 | 0.400649 | 4.546977 | 0.269475 | 1.309277 | 1.31E-10 | 4.35E-09 |
| 330 | PRR15 | 0.363436 | 4.946073 | 0.268421 | 1.307898 | 1.54E-10 | 5.12E-09 |
| 331 | HIST1H4H | 0.082653 | 19.68727 | 0.266863 | 1.305861 | 1.97E-10 | 6.53E-09 |
| 332 | AF127936.7 | 0.464304 | 3.993267 | 0.266384 | 1.305237 | 2.12E-10 | 7.01E-09 |
| 333 | TRBC2 | 0.116216 | 14.13743 | 0.264278 | 1.30249 | 2.95E-10 | 9.71E-09 |
| 334 | HLA-G | 0.983843 | 2.193368 | 0.263584 | 1.301586 | 3.28E-10 | 1.08E-08 |
| 335 | FLRT3 | 0.161829 | 10.28549 | 0.260884 | 1.298078 | 4.98E-10 | 1.63E-08 |
| 336 | GBP5 | 0.731093 | 2.733533 | 0.259028 | 1.29567 | 6.60E-10 | 2.15E-08 |
| 337 | ICAM2 | 0.118063 | 13.83926 | 0.258062 | 1.29442 | 7.65E-10 | 2.49E-08 |
| 338 | CST6 | 7.528953 | 0.797232 | 0.255505 | 1.291114 | 1.12E-09 | 3.64E-08 |
| 339 | DUOX2 | 1.624878 | 1.54823 | 0.254271 | 1.289522 | 1.35E-09 | 4.37E-08 |
| 340 | APLN | 0.105928 | 15.29065 | 0.253736 | 1.288831 | 1.46E-09 | 4.72E-08 |
| 341 | TGFBI | 4.53309 | 0.931167 | 0.25228 | 1.286956 | 1.81E-09 | 5.83E-08 |
| 342 | IDO1 | 2.363911 | 1.243936 | 0.25066 | 1.284873 | 2.30E-09 | 7.38E-08 |
| 343 | SEMA3B | 0.252939 | 6.720662 | 0.24999 | 1.284012 | 2.54E-09 | 8.12E-08 |
| 344 | CRABP2 | 1.385018 | 1.706046 | 0.249444 | 1.283312 | 2.75E-09 | 8.76E-08 |
| 345 | FHL1 | 0.135608 | 12.00953 | 0.248467 | 1.282058 | 3.17E-09 | 1.01E-07 |
| 346 | NDRG1 | 1.050598 | 2.060071 | 0.248301 | 1.281846 | 3.25E-09 | 1.03E-07 |
| 347 | SLC7A2 | 0.116518 | 13.85619 | 0.246669 | 1.279755 | 4.11E-09 | 1.30E-07 |
| 348 | ZNF667-AS1 | 0.862764 | 2.376576 | 0.246425 | 1.279443 | 4.26E-09 | 1.34E-07 |
| 349 | RP11-384O8.1 | 0.161237 | 10.16862 | 0.246004 | 1.278905 | 4.52E-09 | 1.42E-07 |
| 350 | HIP1R | 0.250703 | 6.748031 | 0.245948 | 1.278833 | 4.56E-09 | 1.43E-07 |
| 351 | FGF2 | 2.018302 | 1.348742 | 0.244999 | 1.27762 | 5.23E-09 | 1.63E-07 |
| 352 | KIT | 0.030184 | 51.67617 | 0.24396 | 1.276293 | 6.06E-09 | 1.89E-07 |
| 353 | C1QTNF1-AS1 | 0.10998 | 14.60129 | 0.243673 | 1.275927 | 6.31E-09 | 1.96E-07 |
| 354 | KISS1 | 0.050088 | 31.29963 | 0.241629 | 1.273321 | 8.43E-09 | 2.61E-07 |
| 355 | RP11-95M15.1 | 0.438258 | 4.087299 | 0.240342 | 1.271684 | 1.01E-08 | 3.12E-07 |
| 356 | FAM110C | 0.875345 | 2.334729 | 0.239556 | 1.270685 | 1.13E-08 | 3.47E-07 |
| 357 | CAPN12 | 0.24435 | 6.863296 | 0.239415 | 1.270505 | 1.15E-08 | 3.53E-07 |

| | | | | | | | |
|---|---|---|---|---|---|---|---|
| 358 | AKR1B10 | 0.133243 | 12.09926 | 0.239168 | 1.270192 | 1.19E-08 | 3.65E-07 |
| 359 | PCDH17 | 0.13024 | 12.36324 | 0.239039 | 1.270028 | 1.21E-08 | 3.70E-07 |
| 360 | LXN | 0.208754 | 7.928182 | 0.238607 | 1.26948 | 1.29E-08 | 3.92E-07 |
| 361 | TREM1 | 0.147034 | 10.99752 | 0.237228 | 1.26773 | 1.56E-08 | 4.73E-07 |
| 362 | CYP3A5 | 0.042828 | 36.34043 | 0.237073 | 1.267534 | 1.59E-08 | 4.82E-07 |
| 363 | SOX4 | 1.445921 | 1.638446 | 0.237045 | 1.267498 | 1.60E-08 | 4.83E-07 |
| 364 | PAG1 | 0.130057 | 12.33936 | 0.235761 | 1.265872 | 1.91E-08 | 5.75E-07 |
| 365 | PRDM1 | 0.821944 | 2.438345 | 0.235323 | 1.265317 | 2.03E-08 | 6.08E-07 |
| 366 | EXOC3L4 | 0.045003 | 34.48427 | 0.233365 | 1.262842 | 2.65E-08 | 7.93E-07 |
| 367 | DKK3 | 0.604114 | 3.099196 | 0.232146 | 1.261303 | 3.12E-08 | 9.32E-07 |
| 368 | RP11-456O19.2 | 0.035672 | 43.29901 | 0.232099 | 1.261244 | 3.14E-08 | 9.36E-07 |
| 369 | RP11-402P6.11 | 0.140747 | 11.39774 | 0.231528 | 1.260525 | 3.39E-08 | 1.01E-06 |
| 370 | S1PR3 | 0.285436 | 5.911877 | 0.231483 | 1.260468 | 3.41E-08 | 1.01E-06 |
| 371 | PLPP3 | 0.308894 | 5.506427 | 0.231439 | 1.260412 | 3.43E-08 | 1.01E-06 |
| 372 | LINC00936 | 0.065044 | 23.93239 | 0.229004 | 1.257347 | 4.76E-08 | 1.40E-06 |
| 373 | PTPRR | 0.060492 | 25.68214 | 0.228708 | 1.256976 | 4.95E-08 | 1.45E-06 |
| 374 | DNAJB4 | 0.298351 | 5.644892 | 0.225125 | 1.252479 | 7.94E-08 | 2.33E-06 |
| 375 | HSPA1A | 0.780794 | 2.50876 | 0.224375 | 1.25154 | 8.75E-08 | 2.56E-06 |
| 376 | ALOX15B | 0.174924 | 9.203326 | 0.222871 | 1.249659 | 1.06E-07 | 3.10E-06 |
| 377 | TGFB2 | 0.341724 | 4.963851 | 0.217655 | 1.243158 | 2.07E-07 | 6.03E-06 |
| 378 | NRG1 | 0.221369 | 7.348226 | 0.216904 | 1.242224 | 2.28E-07 | 6.61E-06 |
| 379 | CEACAM6 | 0.106941 | 14.5925 | 0.216152 | 1.241291 | 2.50E-07 | 7.24E-06 |
| 380 | PHKG1 | 0.423745 | 4.106761 | 0.216136 | 1.241271 | 2.51E-07 | 7.24E-06 |
| 381 | ACKR3 | 1.335012 | 1.687814 | 0.214416 | 1.239138 | 3.11E-07 | 8.95E-06 |
| 382 | TNC | 0.464867 | 3.779681 | 0.212445 | 1.236698 | 3.97E-07 | 1.14E-05 |
| 383 | IRS2 | 0.360933 | 4.702302 | 0.211801 | 1.235902 | 4.30E-07 | 1.23E-05 |
| 384 | ZBED2 | 0.92081 | 2.185897 | 0.211486 | 1.235513 | 4.46E-07 | 1.27E-05 |
| 385 | RP11-538P18.2 | 0.217347 | 7.416996 | 0.209288 | 1.2328 | 5.84E-07 | 1.66E-05 |
| 386 | ACTA2 | 0.430981 | 4.016452 | 0.208476 | 1.2318 | 6.44E-07 | 1.83E-05 |
| 387 | GALNT5 | 0.284717 | 5.777392 | 0.206193 | 1.228991 | 8.47E-07 | 2.40E-05 |

| | | | | | | | |
|---|---|---|---|---|---|---|---|
| 388 | UPP1 | 2.860597 | 1.080433 | 0.205892 | 1.22862 | 8.78E-07 | 2.48E-05 |
| 389 | PTN | 0.013831 | 107.6242 | 0.203325 | 1.225471 | 1.19E-06 | 3.35E-05 |
| 390 | PSG5 | 0.06628 | 22.87868 | 0.202349 | 1.224276 | 1.33E-06 | 3.75E-05 |
| 391 | CSF3 | 6.526953 | 0.78594 | 0.202114 | 1.223988 | 1.37E-06 | 3.84E-05 |
| 392 | TMPRSS2 | 0.086917 | 17.56346 | 0.201441 | 1.223164 | 1.48E-06 | 4.15E-05 |
| 393 | NKAIN4 | 0.039678 | 37.78772 | 0.2009 | 1.222502 | 1.58E-06 | 4.40E-05 |
| 394 | KCNQ4 | 0.265915 | 6.113639 | 0.200885 | 1.222485 | 1.58E-06 | 4.40E-05 |
| 395 | CDC20 | 0.148177 | 10.50203 | 0.198465 | 1.219529 | 2.09E-06 | 5.81E-05 |
| 396 | FAM107B | 0.548891 | 3.237891 | 0.197209 | 1.217999 | 2.42E-06 | 6.69E-05 |
| 397 | CITED4 | 0.976057 | 2.063307 | 0.196659 | 1.217329 | 2.57E-06 | 7.10E-05 |
| 398 | SELM | 2.020536 | 1.283855 | 0.196321 | 1.216917 | 2.67E-06 | 7.36E-05 |
| 399 | ALDH1A1 | 0.115046 | 13.33221 | 0.195942 | 1.216457 | 2.79E-06 | 7.67E-05 |
| 400 | CTD-2008A1.3 | 0.048088 | 31.07767 | 0.194506 | 1.214711 | 3.28E-06 | 8.99E-05 |
| 401 | BBC3 | 3.01046 | 1.042614 | 0.194462 | 1.214658 | 3.30E-06 | 9.01E-05 |
| 402 | CCAT1 | 0.047565 | 31.36429 | 0.192954 | 1.212827 | 3.90E-06 | 0.000106 |
| 403 | UNC5B-AS1 | 0.044147 | 33.74847 | 0.192923 | 1.212789 | 3.92E-06 | 0.000107 |
| 404 | PSAT1 | 0.493906 | 3.521049 | 0.192862 | 1.212715 | 3.94E-06 | 0.000107 |
| 405 | ALDH1A3 | 2.339425 | 1.179703 | 0.192131 | 1.211829 | 4.28E-06 | 0.000116 |
| 406 | PGF | 1.417993 | 1.584508 | 0.190918 | 1.210361 | 4.89E-06 | 0.000132 |
| 407 | SGCE | 0.057693 | 25.82901 | 0.18806 | 1.206906 | 6.69E-06 | 0.00018 |
| 408 | HIST1H2BC | 0.427892 | 3.956009 | 0.187126 | 1.20578 | 7.41E-06 | 0.000199 |
| 409 | IL11 | 0.061827 | 24.11506 | 0.187078 | 1.205721 | 7.45E-06 | 0.0002 |
| 410 | NAV3 | 0.243016 | 6.545191 | 0.186944 | 1.20556 | 7.55E-06 | 0.000202 |
| 411 | BBOX1-AS1 | 0.022716 | 64.64566 | 0.186463 | 1.20498 | 7.96E-06 | 0.000212 |
| 412 | PAQR7 | 0.573716 | 3.085834 | 0.185599 | 1.203939 | 8.73E-06 | 0.000232 |
| 413 | LGALS7 | 0.385105 | 4.325182 | 0.185014 | 1.203235 | 9.30E-06 | 0.000247 |
| 414 | NDRG2 | 0.127444 | 11.93478 | 0.183077 | 1.200907 | 1.14E-05 | 0.000302 |
| 415 | C6orf58 | 0.044903 | 32.85737 | 0.182857 | 1.200642 | 1.17E-05 | 0.000309 |
| 416 | LINC01133 | 0.43713 | 3.867352 | 0.182817 | 1.200595 | 1.17E-05 | 0.000309 |
| 417 | CAPNS2 | 1.118822 | 1.841574 | 0.180802 | 1.198178 | 1.45E-05 | 0.000381 |

| | | | | | | | |
|---|---|---|---|---|---|---|---|
| 418 | MGAM | 0.059328 | 24.93516 | 0.180188 | 1.197442 | 1.55E-05 | 0.000406 |
| 419 | NINJ1 | 7.43887 | 0.740216 | 0.178159 | 1.195015 | 1.91E-05 | 0.000499 |
| 420 | MRGPRX4 | 0.03993 | 36.60537 | 0.17536 | 1.191676 | 2.54E-05 | 0.000663 |
| 421 | DUSP2 | 0.181145 | 8.487587 | 0.174653 | 1.190833 | 2.73E-05 | 0.000711 |
| 422 | TMEM47 | 0.08298 | 17.84214 | 0.172279 | 1.188009 | 3.46E-05 | 0.000899 |
| 423 | S100A14 | 27.21845 | 0.595567 | 0.17215 | 1.187856 | 3.51E-05 | 0.000909 |
| 424 | SLCO4A1 | 0.421254 | 3.948741 | 0.171814 | 1.187457 | 3.63E-05 | 0.000938 |
| 425 | SDPR | 0.188655 | 8.141208 | 0.170954 | 1.186437 | 3.95E-05 | 0.001019 |
| 426 | ZNF19 | 0.029981 | 48.33648 | 0.170492 | 1.185889 | 4.13E-05 | 0.001064 |
| 427 | SSPN | 0.415413 | 3.988089 | 0.169674 | 1.184918 | 4.48E-05 | 0.00115 |
| 428 | GPX2 | 1.037469 | 1.921467 | 0.169634 | 1.184871 | 4.50E-05 | 0.001152 |
| 429 | RRM2B | 0.766095 | 2.405969 | 0.167835 | 1.182741 | 5.36E-05 | 0.00137 |
| 430 | MFAP2 | 0.226233 | 6.844233 | 0.16589 | 1.180444 | 6.47E-05 | 0.001649 |
| 431 | SOX7 | 1.74119 | 1.355254 | 0.163403 | 1.177511 | 8.20E-05 | 0.002085 |
| 432 | ZFP57 | 0.147321 | 10.19509 | 0.163317 | 1.17741 | 8.27E-05 | 0.002096 |
| 433 | MRAS | 0.156751 | 9.613986 | 0.163299 | 1.177388 | 8.28E-05 | 0.002096 |
| 434 | FA2H | 0.104581 | 14.10617 | 0.160795 | 1.174444 | 0.000105 | 0.002645 |
| 435 | TMPRSS4 | 0.51973 | 3.266204 | 0.1605 | 1.174098 | 0.000108 | 0.002712 |
| 436 | LRRC8B | 0.342132 | 4.682916 | 0.160451 | 1.17404 | 0.000108 | 0.002719 |
| 437 | HEG1 | 0.394798 | 4.128502 | 0.160138 | 1.173673 | 0.000111 | 0.002792 |
| 438 | IZUMO4 | 0.59429 | 2.922203 | 0.159974 | 1.173481 | 0.000113 | 0.002829 |
| 439 | CSTA | 16.60954 | 0.621542 | 0.159826 | 1.173307 | 0.000115 | 0.002861 |
| 440 | GPAT3 | 0.067051 | 21.66174 | 0.158973 | 1.172306 | 0.000124 | 0.003089 |
| 441 | KISS1R | 0.092626 | 15.81792 | 0.158294 | 1.171511 | 0.000132 | 0.00328 |
| 442 | KLK10 | 3.26289 | 0.969177 | 0.158269 | 1.171482 | 0.000132 | 0.00328 |
| 443 | CDKN2D | 0.515721 | 3.274923 | 0.15669 | 1.169632 | 0.000153 | 0.003781 |
| 444 | INA | 0.204828 | 7.428829 | 0.155941 | 1.168757 | 0.000164 | 0.004037 |
| 445 | SRGAP3 | 0.143018 | 10.40815 | 0.1559 | 1.168709 | 0.000164 | 0.004043 |
| 446 | IGFBP6 | 14.4719 | 0.631035 | 0.154903 | 1.167544 | 0.00018 | 0.004406 |
| 447 | B3GNT7 | 0.177034 | 8.502417 | 0.154896 | 1.167537 | 0.00018 | 0.004406 |

| | | | | | | | |
|---|---|---|---|---|---|---|---|
| 448 | GLIPR1 | 0.502554 | 3.340048 | 0.154694 | 1.167301 | 0.000183 | 0.004476 |
| 449 | GPRC5A | 3.332971 | 0.956042 | 0.154081 | 1.166585 | 0.000193 | 0.004719 |
| 450 | PLAC8 | 0.566019 | 3.018372 | 0.152382 | 1.164605 | 0.000225 | 0.005476 |
| 451 | CORO1A | 0.401575 | 4.03517 | 0.152065 | 1.164236 | 0.000231 | 0.005619 |
| 452 | ENC1 | 0.578869 | 2.960992 | 0.151643 | 1.163745 | 0.00024 | 0.005819 |
| 453 | CXCL14 | 0.037151 | 38.38116 | 0.151611 | 1.163708 | 0.000241 | 0.005823 |
| 454 | LIMD2 | 0.521447 | 3.225531 | 0.150724 | 1.162676 | 0.00026 | 0.00628 |
| 455 | LYPD3 | 0.145256 | 10.19833 | 0.150256 | 1.162132 | 0.000271 | 0.006529 |
| 456 | TGM1 | 0.341657 | 4.637465 | 0.149467 | 1.161216 | 0.00029 | 0.006977 |
| 457 | RASD1 | 0.058231 | 24.62141 | 0.149272 | 1.160989 | 0.000295 | 0.007081 |
| 458 | VAMP5 | 1.002364 | 1.92891 | 0.148659 | 1.160277 | 0.000311 | 0.007451 |
| 459 | EDNRA | 0.192267 | 7.821068 | 0.148528 | 1.160125 | 0.000315 | 0.007519 |
| 460 | RGMB | 0.731571 | 2.445817 | 0.148324 | 1.159889 | 0.00032 | 0.007636 |
| 461 | ALOX5AP | 0.034922 | 40.65659 | 0.148158 | 1.159696 | 0.000325 | 0.007729 |
| 462 | WNT7B | 0.968194 | 1.97704 | 0.148035 | 1.159554 | 0.000328 | 0.007794 |
| 463 | KANK3 | 0.028442 | 49.77033 | 0.147615 | 1.159066 | 0.000341 | 0.008064 |
| 464 | HES1 | 3.062057 | 0.985963 | 0.146505 | 1.157781 | 0.000374 | 0.008847 |
| 465 | STMN1 | 1.257285 | 1.640465 | 0.145703 | 1.156853 | 0.000401 | 0.009451 |
| 466 | HILPDA | 0.558703 | 3.030023 | 0.145507 | 1.156625 | 0.000408 | 0.00959 |

**Supplementary Table S3. HVGs of dermal fibroblasts (FDR<0.01).**

| id | genes | u | cv2 | residualcv2 | fitratio | pval | fdr |
|---|---|---|---|---|---|---|---|
| 1 | LCE1F | 0.116966 | 256.1778 | 3.191764 | 24.33132 | 0 | 0 |
| 2 | KCNMA1 | 2.723042 | 11.75273 | 2.434902 | 11.4147 | 0 | 0 |
| 3 | PI16 | 1.340869 | 14.24888 | 2.272039 | 9.699153 | 0 | 0 |
| 4 | ACAN | 0.146075 | 74.86154 | 2.169615 | 8.754916 | 0 | 0 |
| 5 | IGFBP5 | 0.170898 | 58.27489 | 2.064172 | 7.878774 | 0 | 0 |
| 6 | COMP | 1.341755 | 10.97805 | 2.011647 | 7.475621 | 0 | 0 |
| 7 | KRTAP2-3 | 2.096382 | 7.463947 | 1.864205 | 6.450805 | 0 | 0 |
| 8 | SFRP2 | 1.45289 | 8.632065 | 1.817349 | 6.155521 | 0 | 0 |
| 9 | KRT19 | 0.768361 | 12.18379 | 1.751429 | 5.762833 | 0 | 0 |
| 10 | KRT81 | 0.171725 | 40.45415 | 1.703603 | 5.493706 | 0 | 0 |
| 11 | FLG | 0.020781 | 283.5322 | 1.613618 | 5.020943 | 0 | 0 |
| 12 | KRT14 | 0.062542 | 94.9548 | 1.600267 | 4.954355 | 0 | 0 |
| 13 | G0S2 | 0.086609 | 67.41123 | 1.57121 | 4.81247 | 0 | 0 |
| 14 | CXCL1 | 0.148654 | 36.08072 | 1.455989 | 4.288723 | 0 | 0 |
| 15 | MMP3 | 2.014441 | 4.978754 | 1.440019 | 4.220776 | 0 | 0 |
| 16 | RGCC | 0.310434 | 17.34869 | 1.384953 | 3.994639 | 0 | 0 |
| 17 | CLU | 1.392854 | 5.490137 | 1.340555 | 3.821165 | 4.94e-324 | 1.95e-321 |
| 18 | KRT17 | 0.028573 | 154.7059 | 1.322274 | 3.751943 | 4.71e-313 | 1.75e-310 |
| 19 | STMN2 | 0.184151 | 25.7279 | 1.31496 | 3.724603 | 0.00E+00 | 3.71E-306 |
| 20 | PTX3 | 0.662505 | 8.612073 | 1.296361 | 3.655968 | 7.71E-298 | 2.59E-295 |
| 21 | MGP | 0.053374 | 75.07596 | 1.211488 | 3.358478 | 1.15E-251 | 3.68E-249 |
| 22 | CTSC | 0.624386 | 8.183875 | 1.200948 | 3.323264 | 2.63E-246 | 8.03E-244 |
| 23 | TFPI2 | 0.477804 | 9.92611 | 1.185612 | 3.272689 | 1.19E-238 | 3.46E-236 |
| 24 | PCP4 | 0.025177 | 152.2137 | 1.181254 | 3.258459 | 1.66E-236 | 4.63E-234 |
| 25 | MMP1 | 0.624137 | 7.973244 | 1.174572 | 3.236757 | 3.03E-233 | 8.13E-231 |
| 26 | MT1X | 0.735322 | 7.020708 | 1.168581 | 3.217422 | 2.40E-230 | 6.19E-228 |
| 27 | ACTC1 | 2.807422 | 3.119233 | 1.120927 | 3.067698 | 3.57E-208 | 8.86E-206 |

| | | | | | | | |
|---|---|---|---|---|---|---|---|
| 28 | GAL | 0.413089 | 10.41782 | 1.115735 | 3.051809 | 7.52E-206 | 1.80E-203 |
| 29 | ACTA2 | 0.768912 | 6.432964 | 1.11327 | 3.044297 | 9.39E-205 | 2.17E-202 |
| 30 | HIST1H4C | 0.721375 | 6.265193 | 1.040835 | 2.831579 | 2.83E-174 | 6.33E-172 |
| 31 | LUM | 1.052713 | 4.781408 | 1.030536 | 2.802569 | 3.30E-170 | 7.13E-168 |
| 32 | PPP1R14A | 0.239632 | 14.66802 | 0.99043 | 2.692391 | 5.57E-155 | 1.17E-152 |
| 33 | RARRES2 | 0.279066 | 12.36985 | 0.954309 | 2.596875 | 4.52E-142 | 9.19E-140 |
| 34 | PENK | 0.163029 | 19.10614 | 0.905635 | 2.473502 | 7.85E-126 | 1.55E-123 |
| 35 | DCN | 8.58271 | 1.811251 | 0.897098 | 2.452476 | 4.08E-123 | 7.81E-121 |
| 36 | DKK1 | 6.606759 | 1.901539 | 0.892414 | 2.441016 | 1.21E-121 | 2.26E-119 |
| 37 | SCG5 | 0.408511 | 8.077002 | 0.852053 | 2.344455 | 2.02E-109 | 3.66E-107 |
| 38 | CEMIP | 1.330268 | 3.447448 | 0.848309 | 2.335694 | 2.49E-108 | 4.40E-106 |
| 39 | OLFM2 | 0.475435 | 7.09264 | 0.845517 | 2.329182 | 1.60E-107 | 2.76E-105 |
| 40 | SERPINE2 | 10.74543 | 1.641158 | 0.836042 | 2.307217 | 8.36E-105 | 1.40E-102 |
| 41 | SFRP4 | 0.027489 | 95.64457 | 0.803269 | 2.232829 | 9.44E-96 | 1.54E-93 |
| 42 | IGFBP7 | 4.49476 | 1.913554 | 0.797971 | 2.221029 | 2.45E-94 | 3.91E-92 |
| 43 | PTTG1 | 0.662184 | 5.207442 | 0.792924 | 2.209849 | 5.30E-93 | 8.26E-91 |
| 44 | IL1RL1 | 0.121001 | 22.47245 | 0.790123 | 2.203667 | 2.88E-92 | 4.39E-90 |
| 45 | NMB | 0.471719 | 6.729194 | 0.786619 | 2.19596 | 2.37E-91 | 3.53E-89 |
| 46 | KRT34 | 0.163024 | 16.91028 | 0.783517 | 2.189159 | 1.51E-90 | 2.20E-88 |
| 47 | KRTAP1-5 | 0.41936 | 7.369849 | 0.782 | 2.18584 | 3.73E-90 | 5.32E-88 |
| 48 | TNFRSF11B | 0.829388 | 4.267988 | 0.756483 | 2.130769 | 1.01E-83 | 1.41E-81 |
| 49 | POSTN | 0.485973 | 6.365959 | 0.754967 | 2.127541 | 2.37E-83 | 3.25E-81 |
| 50 | PGF | 0.731107 | 4.594283 | 0.740367 | 2.096704 | 8.12E-80 | 1.09E-77 |
| 51 | AKR1C1 | 0.555307 | 5.554263 | 0.723576 | 2.061793 | 7.15E-76 | 9.41E-74 |
| 52 | STC1 | 0.064104 | 37.78824 | 0.702752 | 2.019302 | 3.74E-71 | 4.82E-69 |
| 53 | MT1E | 1.592697 | 2.611449 | 0.673083 | 1.960271 | 9.28E-65 | 1.17E-62 |
| 54 | ATF5 | 0.536604 | 5.377746 | 0.66459 | 1.943694 | 5.36E-63 | 6.65E-61 |
| 55 | PSG5 | 2.242421 | 2.171606 | 0.661254 | 1.93722 | 2.59E-62 | 3.15E-60 |
| 56 | ANKRD1 | 0.039149 | 58.59 | 0.660827 | 1.936394 | 3.16E-62 | 3.78E-60 |
| 57 | CRIP1 | 1.568983 | 2.595968 | 0.658901 | 1.932667 | 7.79E-62 | 9.17E-60 |

| | | | | | | | |
|---|---|---|---|---|---|---|---|
| 58 | KRT7 | 2.353586 | 2.108295 | 0.653721 | 1.922682 | 8.69E-61 | 1.00E-58 |
| 59 | ADH1B | 0.013303 | 168.2453 | 0.649516 | 1.914614 | 6.03E-60 | 6.86E-58 |
| 60 | FST | 0.638735 | 4.595838 | 0.641039 | 1.898453 | 2.85E-58 | 3.19E-56 |
| 61 | CDKN3 | 0.404235 | 6.596568 | 0.640894 | 1.898178 | 3.05E-58 | 3.35E-56 |
| 62 | STMN1 | 1.024111 | 3.279027 | 0.635456 | 1.887883 | 3.48E-57 | 3.77E-55 |
| 63 | SMYD3 | 0.481787 | 5.679795 | 0.634004 | 1.885144 | 6.65E-57 | 7.08E-55 |
| 64 | ADIRF | 7.288053 | 1.433879 | 0.631446 | 1.880328 | 2.06E-56 | 2.16E-54 |
| 65 | HSPB7 | 2.085822 | 2.170168 | 0.626505 | 1.871059 | 1.81E-55 | 1.87E-53 |
| 66 | SPON2 | 0.377548 | 6.789996 | 0.613023 | 1.846003 | 6.05E-53 | 6.15E-51 |
| 67 | NEFM | 0.044378 | 49.36846 | 0.612282 | 1.844636 | 8.28E-53 | 8.29E-51 |
| 68 | IGF1 | 0.057596 | 37.99746 | 0.604505 | 1.830347 | 2.18E-51 | 2.15E-49 |
| 69 | DPP4 | 0.086195 | 25.74107 | 0.603904 | 1.829245 | 2.80E-51 | 2.72E-49 |
| 70 | UCHL1 | 0.927765 | 3.391709 | 0.603664 | 1.828807 | 3.09E-51 | 2.96E-49 |
| 71 | CRLF1 | 0.497742 | 5.367295 | 0.603384 | 1.828296 | 3.48E-51 | 3.28E-49 |
| 72 | S100A2 | 0.308775 | 7.866426 | 0.589423 | 1.802947 | 1.06E-48 | 9.86E-47 |
| 73 | SERPINB2 | 0.111824 | 19.66877 | 0.582494 | 1.790499 | 1.69E-47 | 1.56E-45 |
| 74 | BAMBI | 0.184133 | 12.32803 | 0.579172 | 1.784561 | 6.30E-47 | 5.71E-45 |
| 75 | CYP1B1 | 0.752623 | 3.819961 | 0.576729 | 1.780206 | 1.65E-46 | 1.47E-44 |
| 76 | ADAMTS1 | 1.108483 | 2.917772 | 0.569684 | 1.767708 | 2.55E-45 | 2.25E-43 |
| 77 | IGFBP2 | 0.242419 | 9.486536 | 0.564912 | 1.759293 | 1.59E-44 | 1.39E-42 |
| 78 | TGM2 | 0.206411 | 10.87677 | 0.557626 | 1.746521 | 2.50E-43 | 2.15E-41 |
| 79 | NQO1 | 2.988034 | 1.704918 | 0.541747 | 1.719007 | 8.69E-41 | 7.38E-39 |
| 80 | TNNT1 | 0.101315 | 20.71061 | 0.540592 | 1.717024 | 1.32E-40 | 1.10E-38 |
| 81 | TUBA1B | 10.17762 | 1.219544 | 0.530678 | 1.700085 | 4.51E-39 | 3.73E-37 |
| 82 | CYTL1 | 0.033776 | 59.41298 | 0.529892 | 1.698748 | 5.95E-39 | 4.86E-37 |
| 83 | PRR15 | 0.029641 | 67.08565 | 0.522872 | 1.686865 | 6.86E-38 | 5.55E-36 |
| 84 | CSRP2 | 0.363502 | 6.389714 | 0.52047 | 1.682818 | 1.57E-37 | 1.25E-35 |
| 85 | HIST1H1C | 0.441478 | 5.434599 | 0.519395 | 1.68101 | 2.27E-37 | 1.79E-35 |
| 86 | OSR2 | 0.365721 | 6.328806 | 0.516009 | 1.675328 | 7.21E-37 | 5.62E-35 |
| 87 | CCL26 | 0.369738 | 6.233917 | 0.510073 | 1.665414 | 5.34E-36 | 4.11E-34 |

| | | | | | | | |
|---|---|---|---|---|---|---|---|
| 88 | WNT2 | 0.105214 | 19.19358 | 0.500362 | 1.649318 | 1.33E-34 | 1.01E-32 |
| 89 | PSG4 | 0.6222 | 4.07092 | 0.500002 | 1.648724 | 1.49E-34 | 1.12E-32 |
| 90 | MT2A | 16.69458 | 1.10569 | 0.496754 | 1.643379 | 4.29E-34 | 3.20E-32 |
| 91 | CFD | 0.260042 | 8.293227 | 0.492551 | 1.636486 | 1.66E-33 | 1.22E-31 |
| 92 | LSP1 | 0.098332 | 20.25492 | 0.489927 | 1.632198 | 3.84E-33 | 2.80E-31 |
| 93 | CLIC3 | 1.356285 | 2.363507 | 0.482228 | 1.619679 | 4.35E-32 | 3.14E-30 |
| 94 | MFGE8 | 3.511804 | 1.500153 | 0.473999 | 1.606406 | 5.52E-31 | 3.94E-29 |
| 95 | COL15A1 | 0.079472 | 24.42339 | 0.473493 | 1.605592 | 6.45E-31 | 4.55E-29 |
| 96 | NDUFA4L2 | 0.113649 | 17.19726 | 0.463506 | 1.589638 | 1.30E-29 | 9.11E-28 |
| 97 | UBE2C | 0.188285 | 10.70774 | 0.458592 | 1.581846 | 5.56E-29 | 3.85E-27 |
| 98 | DPT | 0.828859 | 3.142406 | 0.449884 | 1.56813 | 6.95E-28 | 4.75E-26 |
| 99 | LY6K | 0.199234 | 10.07901 | 0.449436 | 1.567428 | 7.90E-28 | 5.35E-26 |
| 100 | CSTA | 0.04902 | 37.94674 | 0.446284 | 1.562495 | 1.94E-27 | 1.30E-25 |
| 101 | APOE | 0.020572 | 88.8042 | 0.44277 | 1.557015 | 5.23E-27 | 3.47E-25 |
| 102 | PTGDS | 0.11966 | 16.02967 | 0.441787 | 1.555485 | 6.89E-27 | 4.53E-25 |
| 103 | HAPLN1 | 0.055166 | 33.63672 | 0.440725 | 1.553833 | 9.28E-27 | 6.04E-25 |
| 104 | SULF1 | 1.864905 | 1.904322 | 0.440538 | 1.553542 | 9.77E-27 | 6.30E-25 |
| 105 | IFI27 | 0.153928 | 12.63966 | 0.439391 | 1.551762 | 1.35E-26 | 8.59E-25 |
| 106 | TAGLN | 12.45646 | 1.078974 | 0.437734 | 1.549193 | 2.13E-26 | 1.35E-24 |
| 107 | THBD | 0.144933 | 13.32631 | 0.436417 | 1.547154 | 3.07E-26 | 1.92E-24 |
| 108 | FBXO32 | 0.183337 | 10.67678 | 0.431413 | 1.539431 | 1.21E-25 | 7.51E-24 |
| 109 | CDKN1A | 2.499433 | 1.634699 | 0.4259 | 1.530968 | 5.36E-25 | 3.30E-23 |
| 110 | EFEMP1 | 0.357933 | 5.839951 | 0.417501 | 1.518163 | 4.96E-24 | 3.03E-22 |
| 111 | CRABP2 | 2.275213 | 1.681688 | 0.412263 | 1.510232 | 1.93E-23 | 1.17E-21 |
| 112 | PSMC3IP | 0.074969 | 24.25637 | 0.410548 | 1.507644 | 3.01E-23 | 1.80E-21 |
| 113 | GLRX | 1.783268 | 1.876981 | 0.403094 | 1.496447 | 1.99E-22 | 1.18E-20 |
| 114 | CEBPD | 0.460806 | 4.666954 | 0.401829 | 1.494555 | 2.74E-22 | 1.61E-20 |
| 115 | SOCS2 | 0.621805 | 3.690415 | 0.401392 | 1.493903 | 3.05E-22 | 1.78E-20 |
| 116 | BIRC5 | 0.305734 | 6.549906 | 0.397736 | 1.488451 | 7.56E-22 | 4.37E-20 |
| 117 | ITPRIP | 0.073584 | 24.34996 | 0.396453 | 1.486542 | 1.04E-21 | 5.95E-20 |

| | | | | | | | |
|---|---|---|---|---|---|---|---|
| 118 | CLDN11 | 0.618094 | 3.678601 | 0.393658 | 1.482394 | 2.06E-21 | 1.17E-19 |
| 119 | NTF3 | 0.105791 | 17.07268 | 0.388449 | 1.474692 | 7.26E-21 | 4.09E-19 |
| 120 | FGF7 | 3.712581 | 1.341029 | 0.3812 | 1.46404 | 4.06E-20 | 2.27E-18 |
| 121 | EDN1 | 0.090762 | 19.58641 | 0.380012 | 1.462303 | 5.37E-20 | 2.97E-18 |
| 122 | PRSS3 | 0.115772 | 15.53405 | 0.379252 | 1.461191 | 6.41E-20 | 3.52E-18 |
| 123 | SOX4 | 0.408288 | 5.025598 | 0.377127 | 1.458089 | 1.05E-19 | 5.73E-18 |
| 124 | CD9 | 0.129711 | 13.80666 | 0.368246 | 1.445198 | 8.01E-19 | 4.33E-17 |
| 125 | FABP5 | 0.168682 | 10.79306 | 0.365912 | 1.441829 | 1.35E-18 | 7.26E-17 |
| 126 | HSPB6 | 1.603451 | 1.897319 | 0.357295 | 1.429458 | 9.08E-18 | 4.83E-16 |
| 127 | QPRT | 0.133888 | 13.25079 | 0.356816 | 1.428773 | 1.01E-17 | 5.32E-16 |
| 128 | F2R | 0.185986 | 9.776816 | 0.356445 | 1.428243 | 1.09E-17 | 5.72E-16 |
| 129 | ELN | 7.269759 | 1.089008 | 0.355805 | 1.427329 | 1.25E-17 | 6.52E-16 |
| 130 | PARD3 | 0.110247 | 15.87226 | 0.354607 | 1.42562 | 1.63E-17 | 8.39E-16 |
| 131 | RSPO3 | 0.068744 | 24.90264 | 0.353275 | 1.423723 | 2.17E-17 | 1.11E-15 |
| 132 | CDC20 | 0.140204 | 12.62953 | 0.351828 | 1.421664 | 2.95E-17 | 1.50E-15 |
| 133 | CXCL6 | 0.065969 | 25.82698 | 0.349909 | 1.418939 | 4.45E-17 | 2.24E-15 |
| 134 | OLFML2B | 0.104431 | 16.6117 | 0.348803 | 1.417369 | 5.62E-17 | 2.81E-15 |
| 135 | HMOX1 | 0.485552 | 4.234541 | 0.346585 | 1.41423 | 8.98E-17 | 4.46E-15 |
| 136 | MELK | 0.070098 | 24.24357 | 0.34528 | 1.412386 | 1.18E-16 | 5.80E-15 |
| 137 | SOD3 | 0.692254 | 3.220629 | 0.345263 | 1.412362 | 1.18E-16 | 5.80E-15 |
| 138 | SFRP1 | 0.074174 | 22.85911 | 0.340965 | 1.406304 | 2.89E-16 | 1.41E-14 |
| 139 | THBS1 | 2.106952 | 1.620426 | 0.339214 | 1.403843 | 4.15E-16 | 2.00E-14 |
| 140 | CH25H | 0.235784 | 7.751028 | 0.338178 | 1.40239 | 5.13E-16 | 2.46E-14 |
| 141 | CLSPN | 0.029833 | 55.41425 | 0.33808 | 1.402252 | 5.23E-16 | 2.49E-14 |
| 142 | SCUBE3 | 0.055799 | 29.93365 | 0.33518 | 1.398191 | 9.44E-16 | 4.46E-14 |
| 143 | SERPINE1 | 8.954891 | 1.020789 | 0.331297 | 1.392774 | 2.06E-15 | 9.66E-14 |
| 144 | OXTR | 0.431599 | 4.586052 | 0.331181 | 1.392612 | 2.11E-15 | 9.82E-14 |
| 145 | CENPF | 0.166528 | 10.54733 | 0.33106 | 1.392443 | 2.16E-15 | 9.99E-14 |
| 146 | CDA | 0.822067 | 2.8028 | 0.329755 | 1.390628 | 2.80E-15 | 1.29E-13 |
| 147 | TSTD1 | 0.122262 | 14.01322 | 0.32759 | 1.38762 | 4.30E-15 | 1.96E-13 |

| | | | | | | | |
|---|---|---|---|---|---|---|---|
| 148 | PTN | 0.190965 | 9.251261 | 0.325253 | 1.384381 | 6.80E-15 | 3.08E-13 |
| 149 | ITGA2 | 0.131702 | 13.01681 | 0.323599 | 1.382093 | 9.38E-15 | 4.22E-13 |
| 150 | SLC35E3 | 0.084439 | 19.83462 | 0.32353 | 1.381997 | 9.51E-15 | 4.25E-13 |
| 151 | TINAGL1 | 0.061423 | 26.91049 | 0.321901 | 1.379748 | 1.30E-14 | 5.79E-13 |
| 152 | GADD45B | 1.458878 | 1.925182 | 0.319227 | 1.376064 | 2.18E-14 | 9.62E-13 |
| 153 | IGFBP3 | 28.67086 | 0.883752 | 0.31684 | 1.372783 | 3.44E-14 | 1.51E-12 |
| 154 | IL32 | 0.099455 | 16.77956 | 0.312495 | 1.366832 | 7.79E-14 | 3.39E-12 |
| 155 | SNCG | 0.169912 | 10.15118 | 0.311274 | 1.365164 | 9.78E-14 | 4.23E-12 |
| 156 | PEG10 | 0.04158 | 38.89141 | 0.31003 | 1.363466 | 1.23E-13 | 5.30E-12 |
| 157 | UBE2S | 0.481724 | 4.105929 | 0.309417 | 1.362631 | 1.38E-13 | 5.90E-12 |
| 158 | AKAP12 | 0.295165 | 6.17727 | 0.30873 | 1.361695 | 1.57E-13 | 6.65E-12 |
| 159 | GLIPR1 | 1.496973 | 1.870152 | 0.304807 | 1.356364 | 3.21E-13 | 1.35E-11 |
| 160 | HAPLN3 | 0.271393 | 6.618928 | 0.304588 | 1.356067 | 3.34E-13 | 1.40E-11 |
| 161 | H2AFZ | 5.106357 | 1.123548 | 0.302066 | 1.352651 | 5.27E-13 | 2.20E-11 |
| 162 | COL8A1 | 1.049108 | 2.311263 | 0.301377 | 1.351719 | 5.97E-13 | 2.47E-11 |
| 163 | ADRA2A | 0.139963 | 12.02453 | 0.301136 | 1.351393 | 6.23E-13 | 2.56E-11 |
| 164 | AP5B1 | 0.086426 | 18.87444 | 0.296183 | 1.344716 | 1.50E-12 | 6.14E-11 |
| 165 | FBN2 | 0.146127 | 11.45696 | 0.292899 | 1.340307 | 2.67E-12 | 1.08E-10 |
| 166 | MOK | 0.136193 | 12.16953 | 0.287648 | 1.333288 | 6.58E-12 | 2.66E-10 |
| 167 | TM4SF1 | 0.038617 | 40.80833 | 0.285731 | 1.330734 | 9.11E-12 | 3.66E-10 |
| 168 | TSPAN13 | 0.20417 | 8.317419 | 0.279486 | 1.32245 | 2.59E-11 | 1.03E-09 |
| 169 | PCOLCE | 1.662039 | 1.721325 | 0.279364 | 1.322288 | 2.64E-11 | 1.05E-09 |
| 170 | MARCKSL1 | 0.406172 | 4.575988 | 0.279116 | 1.321961 | 2.75E-11 | 1.08E-09 |
| 171 | SMC6 | 0.147188 | 11.22071 | 0.278787 | 1.321525 | 2.90E-11 | 1.14E-09 |
| 172 | F3 | 0.426011 | 4.39831 | 0.278727 | 1.321446 | 2.93E-11 | 1.14E-09 |
| 173 | DKK2 | 0.072676 | 21.89876 | 0.278385 | 1.320995 | 3.10E-11 | 1.20E-09 |
| 174 | SUGCT | 0.224016 | 7.637998 | 0.277753 | 1.32016 | 3.44E-11 | 1.33E-09 |
| 175 | LNPEP | 0.088196 | 18.11197 | 0.27434 | 1.315662 | 5.99E-11 | 2.30E-09 |
| 176 | HAS2 | 0.403106 | 4.555848 | 0.268443 | 1.307927 | 1.54E-10 | 5.86E-09 |
| 177 | HES1 | 0.190143 | 8.765556 | 0.2674 | 1.306563 | 1.81E-10 | 6.86E-09 |

| | | | | | | | |
|---|---|---|---|---|---|---|---|
| 178 | SPEN | 0.132215 | 12.25523 | 0.26695 | 1.305975 | 1.94E-10 | 7.32E-09 |
| 179 | LMCD1 | 0.358028 | 5.017507 | 0.265935 | 1.30465 | 2.28E-10 | 8.54E-09 |
| 180 | DIRAS3 | 0.052595 | 29.53257 | 0.26417 | 1.30235 | 3.00E-10 | 1.12E-08 |
| 181 | NREP | 1.655881 | 1.696241 | 0.262691 | 1.300425 | 3.77E-10 | 1.40E-08 |
| 182 | MEDAG | 0.15925 | 10.22609 | 0.258929 | 1.295542 | 6.71E-10 | 2.47E-08 |
| 183 | DDX10 | 0.082304 | 19.0348 | 0.257826 | 1.294114 | 7.92E-10 | 2.90E-08 |
| 184 | BCL2L12 | 0.195616 | 8.454101 | 0.257011 | 1.29306 | 8.96E-10 | 3.27E-08 |
| 185 | CKB | 0.499683 | 3.783806 | 0.256882 | 1.292892 | 9.14E-10 | 3.31E-08 |
| 186 | CCL2 | 0.066974 | 23.18539 | 0.256639 | 1.292578 | 9.48E-10 | 3.42E-08 |
| 187 | PRSS23 | 4.554094 | 1.105978 | 0.253652 | 1.288723 | 1.48E-09 | 5.31E-08 |
| 188 | CXCL14 | 0.030536 | 49.58458 | 0.249863 | 1.283849 | 2.59E-09 | 9.23E-08 |
| 189 | STC2 | 1.741791 | 1.630169 | 0.249829 | 1.283805 | 2.60E-09 | 9.23E-08 |
| 190 | CLEC2B | 0.315111 | 5.498446 | 0.248763 | 1.282438 | 3.04E-09 | 1.07E-07 |
| 191 | ISG20 | 0.041884 | 36.27899 | 0.247637 | 1.280995 | 3.58E-09 | 1.26E-07 |
| 192 | HSPA5 | 1.487566 | 1.770381 | 0.246432 | 1.279452 | 4.25E-09 | 1.49E-07 |
| 193 | CENPM | 0.174566 | 9.269392 | 0.245208 | 1.277886 | 5.07E-09 | 1.76E-07 |
| 194 | MLLT11 | 0.218646 | 7.524312 | 0.240995 | 1.272515 | 9.22E-09 | 3.19E-07 |
| 195 | ZNF232 | 0.03174 | 47.2785 | 0.240276 | 1.2716 | 1.02E-08 | 3.51E-07 |
| 196 | ID4 | 0.259466 | 6.444308 | 0.238356 | 1.269161 | 1.33E-08 | 4.56E-07 |
| 197 | APBB1IP | 0.186335 | 8.671516 | 0.238184 | 1.268942 | 1.37E-08 | 4.64E-07 |
| 198 | TRIB3 | 0.452109 | 4.023826 | 0.238155 | 1.268906 | 1.37E-08 | 4.64E-07 |
| 199 | C11orf96 | 0.076395 | 20.0185 | 0.236665 | 1.267016 | 1.68E-08 | 5.68E-07 |
| 200 | PTGER1 | 0.081713 | 18.76255 | 0.236511 | 1.266821 | 1.72E-08 | 5.77E-07 |
| 201 | FAM43A | 0.149546 | 10.58316 | 0.235044 | 1.264965 | 2.10E-08 | 7.02E-07 |
| 202 | CD70 | 0.026954 | 55.23288 | 0.23479 | 1.264643 | 2.18E-08 | 7.24E-07 |
| 203 | S1PR3 | 0.057546 | 26.25956 | 0.234165 | 1.263853 | 2.37E-08 | 7.84E-07 |
| 204 | PALM | 0.127561 | 12.25871 | 0.233668 | 1.263225 | 2.54E-08 | 8.35E-07 |
| 205 | INHBA | 0.500122 | 3.688165 | 0.231977 | 1.261091 | 3.19E-08 | 1.04E-06 |
| 206 | MFAP4 | 0.695052 | 2.864022 | 0.230878 | 1.259706 | 3.70E-08 | 1.21E-06 |
| 207 | TACC3 | 0.054128 | 27.77632 | 0.230809 | 1.259619 | 3.74E-08 | 1.21E-06 |

| | | | | | | | |
|---|---|---|---|---|---|---|---|
| 208 | FMN1 | 0.075192 | 20.12972 | 0.226928 | 1.25474 | 6.26E-08 | 2.02E-06 |
| 209 | TROAP | 0.089908 | 16.95662 | 0.226802 | 1.254582 | 6.37E-08 | 2.04E-06 |
| 210 | SERPINF1 | 0.316803 | 5.354254 | 0.226789 | 1.254565 | 6.38E-08 | 2.04E-06 |
| 211 | TPX2 | 0.125577 | 12.33973 | 0.225543 | 1.253003 | 7.51E-08 | 2.39E-06 |
| 212 | KLHL24 | 0.07263 | 20.7529 | 0.224028 | 1.251106 | 9.16E-08 | 2.90E-06 |
| 213 | CADM1 | 0.042234 | 35.10161 | 0.222781 | 1.249547 | 1.08E-07 | 3.39E-06 |
| 214 | PRC1 | 0.098006 | 15.51308 | 0.220054 | 1.246144 | 1.53E-07 | 4.79E-06 |
| 215 | SYNPO2 | 0.284788 | 5.831065 | 0.219951 | 1.246015 | 1.55E-07 | 4.83E-06 |
| 216 | DNAJC18 | 0.093585 | 16.20261 | 0.219579 | 1.245552 | 1.62E-07 | 5.04E-06 |
| 217 | IER3 | 0.660822 | 2.939496 | 0.219541 | 1.245505 | 1.63E-07 | 5.04E-06 |
| 218 | FOLR3 | 0.071212 | 21.03396 | 0.218482 | 1.244187 | 1.87E-07 | 5.74E-06 |
| 219 | PHACTR3 | 0.173161 | 9.051554 | 0.214012 | 1.238637 | 3.27E-07 | 1.00E-05 |
| 220 | BAALC | 1.15632 | 1.990736 | 0.213968 | 1.238584 | 3.29E-07 | 1.00E-05 |
| 221 | CCDC8 | 0.079766 | 18.767 | 0.213603 | 1.23813 | 3.44E-07 | 1.04E-05 |
| 222 | PLXNA2 | 0.080055 | 18.64171 | 0.210376 | 1.234142 | 5.11E-07 | 1.54E-05 |
| 223 | SRGN | 0.056725 | 26.00138 | 0.210337 | 1.234094 | 5.14E-07 | 1.55E-05 |
| 224 | CARD9 | 0.079272 | 18.79929 | 0.20936 | 1.232888 | 5.79E-07 | 1.73E-05 |
| 225 | DDIT4 | 0.556021 | 3.311077 | 0.207281 | 1.230329 | 7.43E-07 | 2.22E-05 |
| 226 | VTA1 | 0.134431 | 11.35709 | 0.206386 | 1.229227 | 8.28E-07 | 2.46E-05 |
| 227 | CHN1 | 0.292911 | 5.605179 | 0.204894 | 1.227395 | 9.88E-07 | 2.92E-05 |
| 228 | PDE1A | 0.045933 | 31.71032 | 0.20326 | 1.225391 | 1.20E-06 | 3.53E-05 |
| 229 | LBH | 0.864537 | 2.383763 | 0.202843 | 1.224881 | 1.26E-06 | 3.69E-05 |
| 230 | MSX1 | 0.382491 | 4.454362 | 0.202318 | 1.224238 | 1.34E-06 | 3.90E-05 |
| 231 | IL7R | 0.153586 | 9.991675 | 0.202244 | 1.224146 | 1.35E-06 | 3.92E-05 |
| 232 | YLPM1 | 0.135055 | 11.25992 | 0.202121 | 1.223997 | 1.37E-06 | 3.96E-05 |
| 233 | PAPPA | 0.450149 | 3.892331 | 0.201409 | 1.223125 | 1.49E-06 | 4.29E-05 |
| 234 | KCTD5 | 0.109951 | 13.6406 | 0.200541 | 1.222063 | 1.65E-06 | 4.72E-05 |
| 235 | SYNGR2 | 0.050179 | 28.99905 | 0.200144 | 1.221579 | 1.72E-06 | 4.92E-05 |
| 236 | NGFR | 0.041255 | 35.0491 | 0.198326 | 1.219359 | 2.13E-06 | 6.04E-05 |
| 237 | GTSE1 | 0.046375 | 31.2468 | 0.197884 | 1.218821 | 2.24E-06 | 6.33E-05 |

| | | | | | | | |
|---|---|---|---|---|---|---|---|
| 238 | SCARA3 | 0.438764 | 3.950977 | 0.195553 | 1.215984 | 2.92E-06 | 8.22E-05 |
| 239 | TK1 | 0.447681 | 3.883521 | 0.194685 | 1.214928 | 3.22E-06 | 9.03E-05 |
| 240 | TUBB4B | 1.058007 | 2.062913 | 0.19316 | 1.213077 | 3.82E-06 | 0.000107 |
| 241 | MAFB | 0.100469 | 14.7371 | 0.192341 | 1.212084 | 4.18E-06 | 0.000116 |
| 242 | CYGB | 0.265271 | 6.033427 | 0.191947 | 1.211607 | 4.37E-06 | 0.000121 |
| 243 | SBSN | 0.313188 | 5.221232 | 0.19177 | 1.211392 | 4.45E-06 | 0.000123 |
| 244 | HMGCS1 | 0.064039 | 22.6313 | 0.189107 | 1.20817 | 5.97E-06 | 0.000164 |
| 245 | TRIP13 | 0.062291 | 23.22981 | 0.188408 | 1.207326 | 6.44E-06 | 0.000176 |
| 246 | TNFAIP6 | 0.084425 | 17.31688 | 0.187631 | 1.206389 | 7.01E-06 | 0.000191 |
| 247 | TOP2A | 0.094227 | 15.57072 | 0.186309 | 1.204794 | 8.09E-06 | 0.00022 |
| 248 | SLC20A1 | 0.178562 | 8.55768 | 0.186049 | 1.204482 | 8.32E-06 | 0.000225 |
| 249 | WNT5B | 0.736797 | 2.624161 | 0.185928 | 1.204335 | 8.43E-06 | 0.000227 |
| 250 | TNFRSF19 | 0.082629 | 17.6417 | 0.185603 | 1.203944 | 8.73E-06 | 0.000234 |
| 251 | TUBA1A | 2.204813 | 1.356384 | 0.182762 | 1.200529 | 1.18E-05 | 0.000316 |
| 252 | TSPAN2 | 0.153199 | 9.804128 | 0.180958 | 1.198364 | 1.43E-05 | 0.00038 |
| 253 | MKI67 | 0.115862 | 12.72036 | 0.180153 | 1.197401 | 1.55E-05 | 0.000412 |
| 254 | CST6 | 0.096073 | 15.18758 | 0.179887 | 1.197082 | 1.60E-05 | 0.000421 |
| 255 | SLC7A11 | 0.373219 | 4.44277 | 0.179201 | 1.196261 | 1.71E-05 | 0.000451 |
| 256 | FECH | 0.106813 | 13.69997 | 0.177469 | 1.194191 | 2.05E-05 | 0.000537 |
| 257 | SMIM3 | 0.43349 | 3.917822 | 0.177267 | 1.19395 | 2.09E-05 | 0.000546 |
| 258 | LYPD6B | 0.117119 | 12.52006 | 0.174458 | 1.1906 | 2.78E-05 | 0.000724 |
| 259 | LMOD1 | 0.04085 | 34.53324 | 0.173831 | 1.189855 | 2.96E-05 | 0.000768 |
| 260 | MARCH4 | 0.230617 | 6.705067 | 0.173452 | 1.189404 | 3.08E-05 | 0.000794 |
| 261 | BDKRB1 | 0.099371 | 14.60773 | 0.173079 | 1.188959 | 3.20E-05 | 0.000822 |
| 262 | GALE | 0.08952 | 16.13519 | 0.173009 | 1.188877 | 3.22E-05 | 0.000824 |
| 263 | NRG1 | 0.394998 | 4.210279 | 0.172733 | 1.188549 | 3.31E-05 | 0.000844 |
| 264 | CNN1 | 0.185268 | 8.15301 | 0.17129 | 1.186835 | 3.82E-05 | 0.00097 |
| 265 | SLC4A7 | 0.094727 | 15.24876 | 0.170465 | 1.185857 | 4.14E-05 | 0.001049 |
| 266 | PHF7 | 0.039204 | 35.8134 | 0.169946 | 1.18524 | 4.36E-05 | 0.0011 |
| 267 | ALDH1A3 | 0.296679 | 5.347959 | 0.169004 | 1.184125 | 4.78E-05 | 0.001202 |

| | | | | | | | |
|---|---|---|---|---|---|---|---|
| 268 | FAH | 0.28295 | 5.567127 | 0.167987 | 1.182922 | 5.28E-05 | 0.001317 |
| 269 | LAPTM5 | 0.05272 | 26.7627 | 0.167987 | 1.182922 | 5.28E-05 | 0.001317 |
| 270 | COL3A1 | 8.112822 | 0.88267 | 0.167725 | 1.182611 | 5.42E-05 | 0.001346 |
| 271 | WNT16 | 0.05521 | 25.57278 | 0.167412 | 1.182241 | 5.59E-05 | 0.001382 |
| 272 | ANG | 0.10557 | 13.71186 | 0.167248 | 1.182047 | 5.67E-05 | 0.001399 |
| 273 | FGF9 | 0.05543 | 25.40319 | 0.164615 | 1.178939 | 7.31E-05 | 0.001795 |
| 274 | WDYHV1 | 0.162206 | 9.145461 | 0.164215 | 1.178467 | 7.59E-05 | 0.001858 |
| 275 | NPW | 0.134186 | 10.89801 | 0.163416 | 1.177527 | 8.19E-05 | 0.001997 |
| 276 | PIP5KL1 | 0.139531 | 10.50623 | 0.163275 | 1.17736 | 8.30E-05 | 0.002017 |
| 277 | FLNC | 1.273442 | 1.778797 | 0.160569 | 1.174179 | 0.000107 | 0.002589 |
| 278 | ADM | 3.436383 | 1.100532 | 0.156487 | 1.169395 | 0.000156 | 0.003755 |
| 279 | PINLYP | 0.083038 | 17.04981 | 0.156214 | 1.169076 | 0.00016 | 0.003835 |
| 280 | TNXB | 0.15871 | 9.254122 | 0.155916 | 1.168728 | 0.000164 | 0.003925 |
| 281 | RCAN1 | 0.270406 | 5.720378 | 0.155498 | 1.168239 | 0.00017 | 0.004062 |
| 282 | CEP55 | 0.053625 | 25.96573 | 0.15432 | 1.166864 | 0.000189 | 0.0045 |
| 283 | PRELP | 0.091147 | 15.5618 | 0.154034 | 1.16653 | 0.000194 | 0.0046 |
| 284 | CCNB1 | 0.099318 | 14.33446 | 0.153685 | 1.166123 | 0.0002 | 0.004729 |
| 285 | TMEM200A | 0.088745 | 15.9209 | 0.151329 | 1.163379 | 0.000247 | 0.005805 |
| 286 | ASPM | 0.05845 | 23.79178 | 0.150615 | 1.162549 | 0.000263 | 0.006147 |
| 287 | ERRFI1 | 0.083704 | 16.82512 | 0.150597 | 1.162528 | 0.000263 | 0.006147 |
| 288 | APOD | 0.138159 | 10.46919 | 0.150526 | 1.162446 | 0.000265 | 0.006163 |
| 289 | PLAU | 0.061375 | 22.59552 | 0.146374 | 1.15763 | 0.000379 | 0.008789 |
| 290 | NUDCD1 | 0.042305 | 32.4575 | 0.146102 | 1.157314 | 0.000388 | 0.008964 |
| 291 | HLA-DPB1 | 0.203201 | 7.302613 | 0.145065 | 1.156115 | 0.000423 | 0.009753 |

**Supplementary Table S4. Significantly enriched motifs found in HVGs.**

Results were generated by GREAT (version 3.0.0) using hg19 human genome reference with association rule: Basal+extension: 5000 bp upstream, 1000 bp downstream, 1000000 bp max extension, curated regulatory domains included.

| Cell type | # Term Name | Binom Rank | Binom Raw P-Value | Binom FDR Q-Val | Binom Fold Enrichment | Binom Obs. Region Hits | Binom Region Set Cov. |
|---|---|---|---|---|---|---|---|
| LCL | Motif NNNNKGGRAANTCCCN (no known TF) | 1 | 3.35E-06 | 0.002058 | 2.685652 | 28 | 0.066038 |
| LCL | Motif GGGAMTTYCC matches NFKB; RELA: v-rel reticuloendotheliosis viral oncogene homolog A, nuclear factor of kappa light polypeptide gene enhancer in B-cells 3, p65 (avian) | 2 | 2.51E-05 | 0.007732 | 2.500147 | 26 | 0.061321 |
| LCL | Motif NGGGACTTTCCA (no known TF) | 3 | 7.05E-05 | 0.014461 | 2.346414 | 26 | 0.061321 |
| LCL | Motif SRTGAGTCANC matches BACH2: BTB and CNC homology 1, basic leucine zipper transcription factor 2 | 5 | 0.000357 | 0.043963 | 2.236156 | 23 | 0.054245 |
| LAEC | Motif STATAAAWRNNNNNN matches TAF<br> TATA | 1 | 1.89E-05 | 0.011642 | 2.441259 | 28 | 0.06422 |
| LAEC | Motif NNNNKGGRAANTCCCN (no known TF) | 3 | 0.000107 | 0.022002 | 2.331906 | 25 | 0.057339 |
| LAEC | Motif NGGGGAMTTTCCNN (no known TF) | 4 | 0.000218 | 0.033477 | 2.1836 | 26 | 0.059633 |
| LAEC | Motif GGGAMTTYCC matches NFKB; RELA: v-rel reticuloendotheliosis viral oncogene homolog A, nuclear factor of kappa light polypeptide gene enhancer in B-cells 3, p65 (avian) | 5 | 0.000256 | 0.031449 | 2.24431 | 24 | 0.055046 |
| LAEC | Motif TTGCWCAAY matches CEBPB: CCAAT/enhancer binding protein (C/EBP), beta | 6 | 0.000471 | 0.048241 | 3.215633 | 12 | 0.027523 |
