## Supplementary material for "Extent, heritability, and functional relevance of single cell expression variability in highly homogeneous populations of human cells": Suppl Figures

### Supplementary Figures

**Supplementary Fig. S1. Flowchart of selection of highly homogeneous population of cells.**

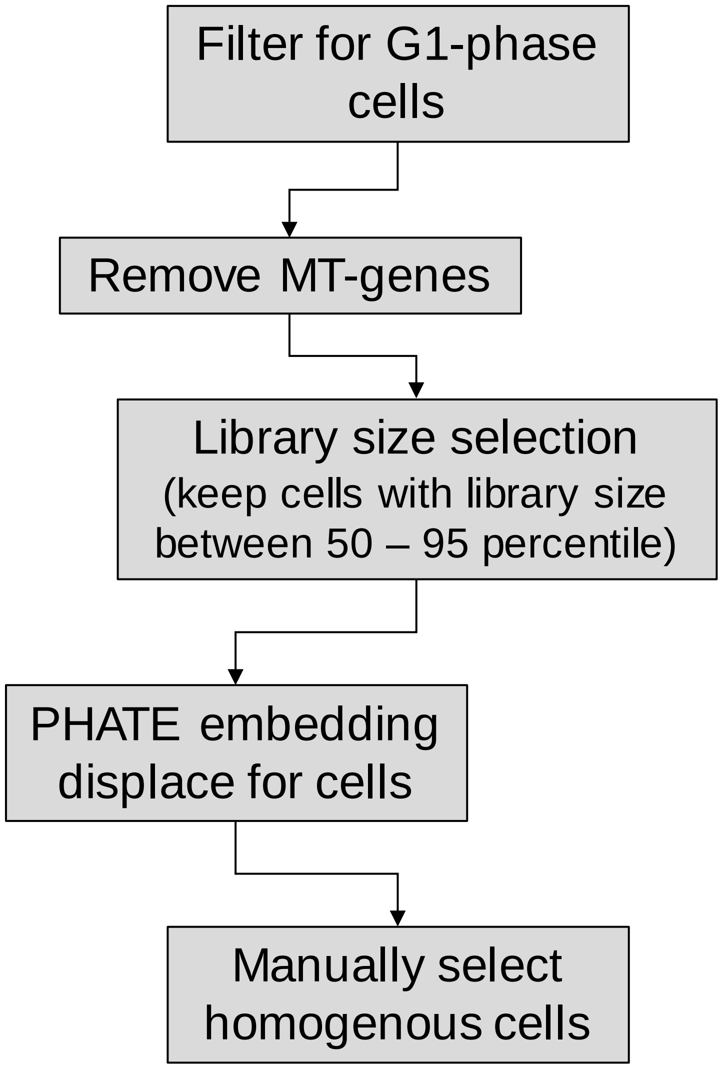

**Supplementary Fig. S2. Correlation analyses of scEV between two genetically related LCL samples and between two genetically unrelated LCL samples.**

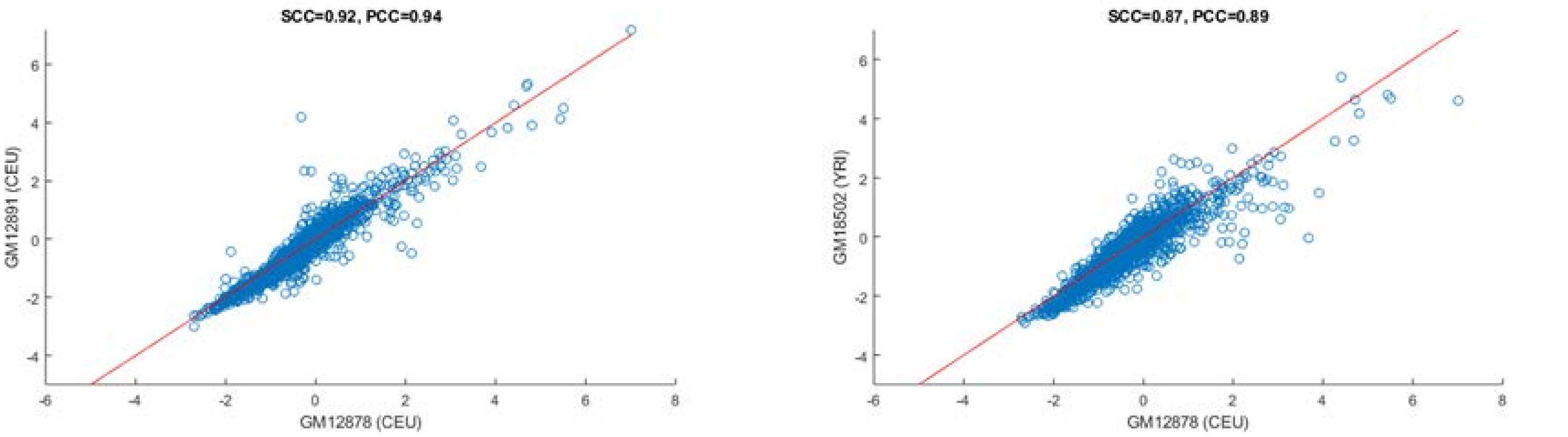

**Supplementary Fig. S3. Correlation between scEV and population-level expression variability across genes of functional sets.**

1-8 gene sets are defined by GO terms for biological process (BP); 9-16 molecular function (MF). Four subplots highlighted with an orange box are those also shown in the figure in the main text.

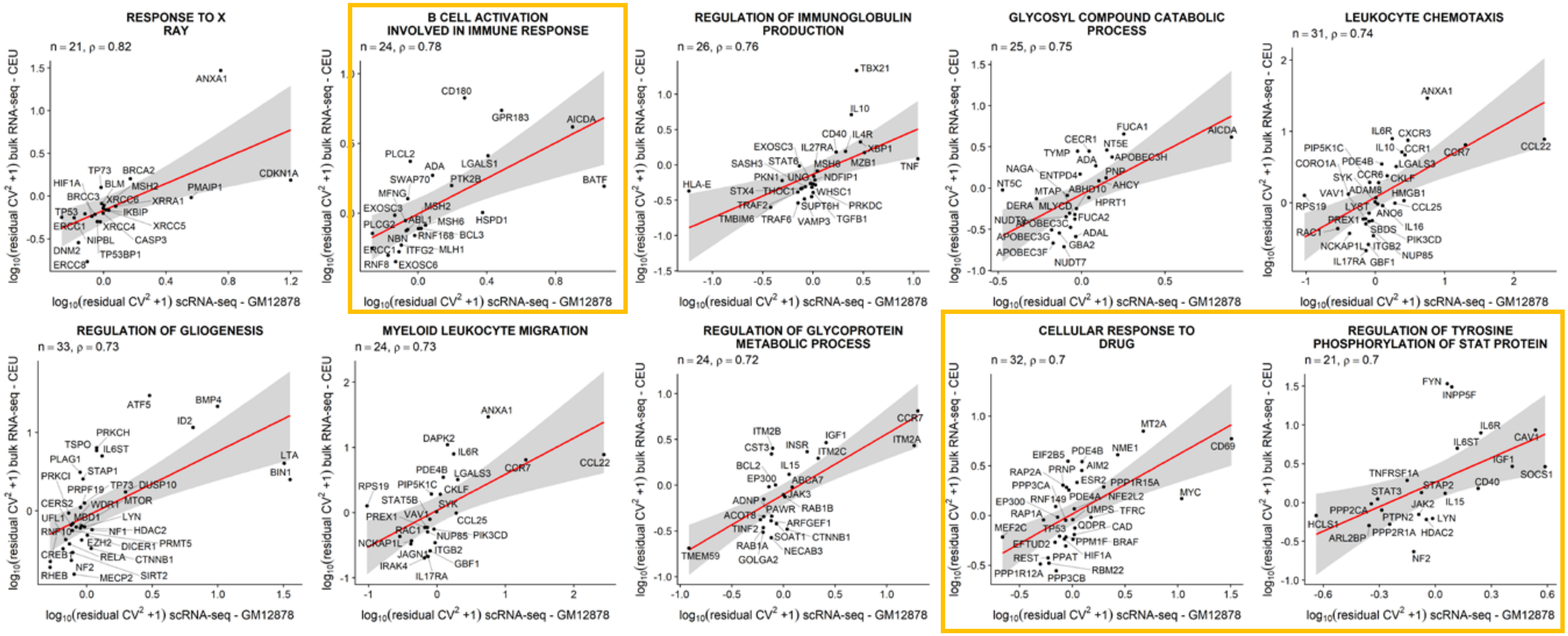

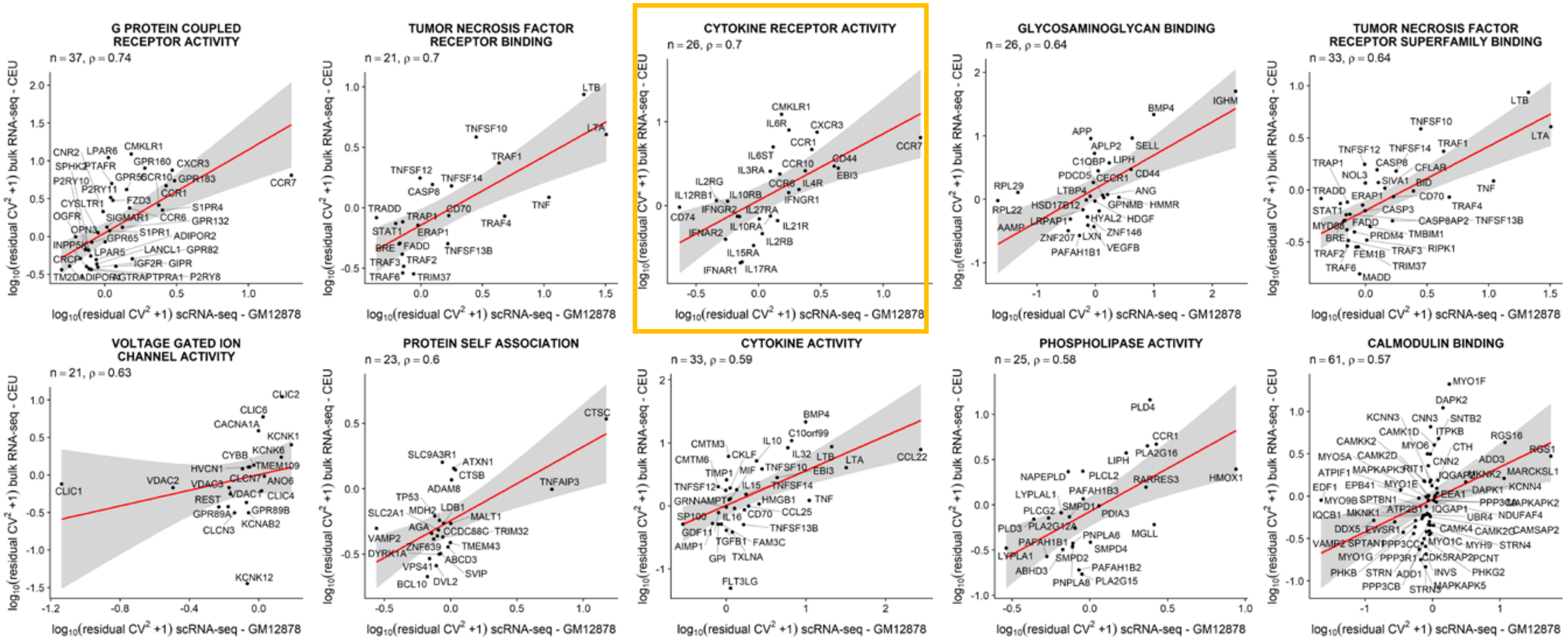
G PROTEIN COUPLED RECEPTOR ACTIVITY
n = 37, ρ = 0.74
TUMOR NECROSIS FACTOR RECEPTOR BINDING
n = 21, ρ = 0.7
CYTOKINE RECEPTOR ACTIVITY
n = 26, ρ = 0.7
GLYCOSAMINOGLYCAN BINDING
n = 26, ρ = 0.64
TUMOR NECROSIS FACTOR RECEPTOR SUPERFAMILY BINDING
n = 33, ρ = 0.64
VOLTAGE GATED ION CHANNEL ACTIVITY
n = 21, ρ = 0.63
PROTEIN SELF ASSOCIATION
n = 23, ρ = 0.6
CYTOKINE ACTIVITY
n = 33, ρ = 0.59
PHOSPHOLIPASE ACTIVITY
n = 25, ρ = 0.58
CALMODULIN BINDING
n = 61, ρ = 0.57
log10(residual CV² +1) bulk RNA-seq - CEU
log10(residual CV² +1) scRNA-seq - GM12878

**Supplementary Fig. S4. PHATE 3-D embedding plot for cells colored according to IGLC2 expression level in cells.**

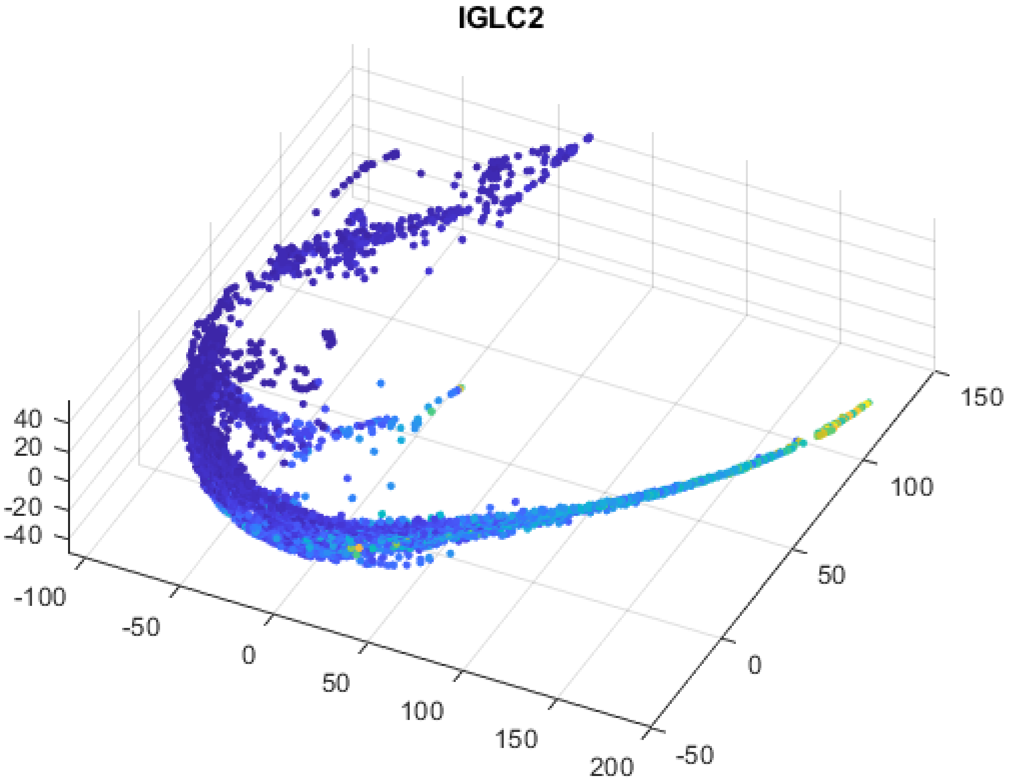

**Supplementary Fig. S5. The relationship between CV2 and mean expression of genes in human iPSCs.**

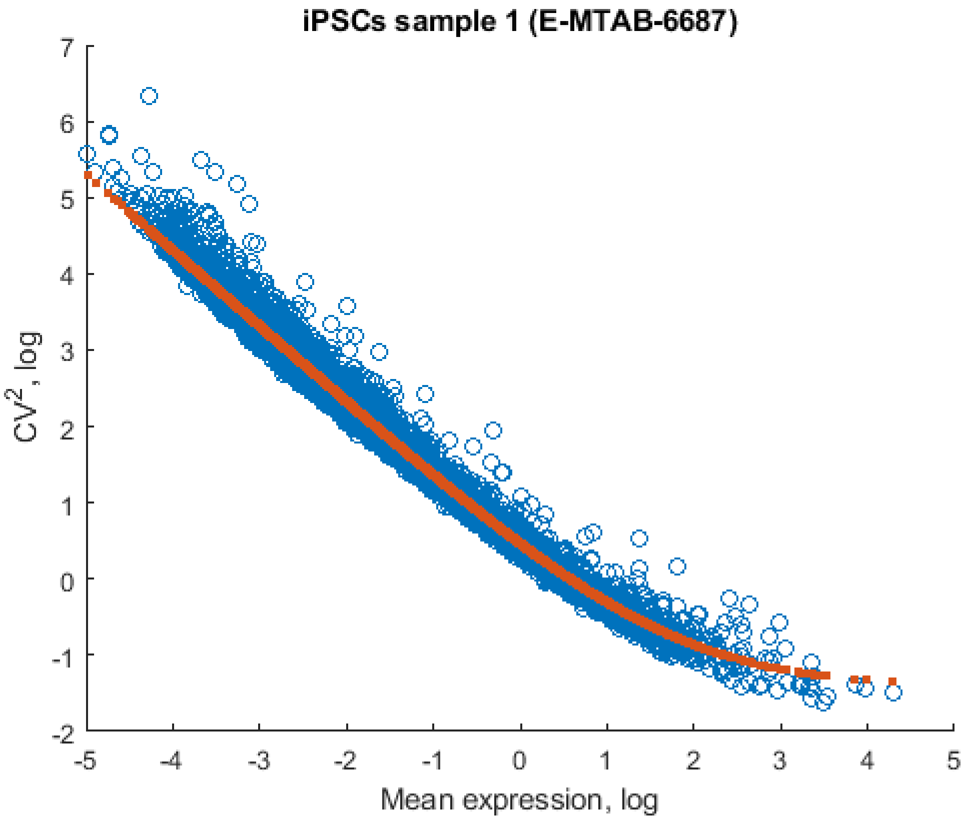
